# A *Very Oil Yellow1* modifier of the *Oil Yellow1-N1989* allele uncovers a cryptic phenotypic impact of cis-regulatory variation in maize

**DOI:** 10.1101/230375

**Authors:** Rajdeep S. Khangura, Sandeep Marla, Bala P. Venkata, Nicholas J. Heller, Gurmukh S. Johal, Brian P. Dilkes

## Abstract

Forward genetics determines the function of genes underlying trait variation by identifying the change in DNA responsible for changes in phenotype. Detecting phenotypically-relevant variation outside protein coding sequences and distinguishing this from neutral variants is not trivial; partly because the mechanisms by which DNA polymorphisms in the intergenic regions affect gene regulation are poorly understood. Here we utilized a dominant genetic marker with a convenient phenotype to investigate the effect of cis and trans-acting regulatory variation. We performed a forward genetic screen for natural variation that suppress or enhance the semi-dominant mutant allele *Oy1*-*N1989,* encoding the magnesium chelatase subunit I of maize. This mutant permits rapid phenotyping of leaf color as a reporter for chlorophyll accumulation, and mapping of natural variation in maize affecting chlorophyll metabolism. We identified a single modifier locus segregating between B73 and Mo17 that was linked to the reporter gene itself, which we call *very oil yellow1*. Based on the variation in OY1 transcript abundance and genome-wide association data, *vey1* is predicted to consist of multiple cis-acting regulatory sequence polymorphisms encoded at the wild-type *oy1* alleles. The *vey1* allele appears to be a common polymorphism in the maize germplasm that alters the expression level of a key gene in chlorophyll biosynthesis. These *vey1* alleles have no discernable impact on leaf chlorophyll in the absence of the *Oy1-N1989* reporter. Thus, use of a mutant as a simple and efficient reporter for magnesium chelatase activity resulted in the detection of expression-level polymorphisms not readily visible in the laboratory.

## Introduction

Genetic variation in the coding or regulatory sequences causes differences between genotypes and is fundamental to crop improvement (Springer and Stupar 2007). Gene function discovery via mutant analyses focuses on linking phenotype alterations to gene variants. As a result, forward genetics has been of great value to understand biological systems but is predominantly useful for determining functions for genes with alleles that cause large phenotypic impacts. Natural variants, including those that encode alleles of relevance to adaptation and fitness of the organism, are often of small effect (Fisher 1930; Orr 1998, 2005). Mutant alleles with conditional impacts on phenotypes, through genetic interactions or modifiers, as well as alleles of small individual effect are difficult to study. Further, even if we identify loci that have not been previously associated with a biological process, it can be difficult to validate and associate such natural variants with physiological and biochemical mechanisms.

A forward genetics approach that uses a mutant phenotype as a reporter for genetic interactions can be used to detect modifiers in natural populations and expose cryptic variation affecting traits of interest (Johal *et al*. 2008). This Mutant-Assisted Gene Identification and Characterization (MAGIC) approach is particularly efficient in species where outcrossing is easy, such as maize (*Zea mays*). It can be applied to any mutant with any quantifiable phenotype to expand our understanding of the process disrupted by mutation. This approach is convenient when a dominant mutant allele is used as a reporter because natural variants that encode enhancers or suppressors of a given mutant phenotype can be detected in F_1_ crosses. Thus, it enables easy detection of dominant enhancers and suppressors. Any germplasm collection, diversity panel, or line-cross population that can serve as the variable parents in these crosses, can be utilized to detect and map loci that alter mutant phenotype expression. Natural variants discovered this way have an experimental link (genetic modifiers) of the processes affected by the mutant reporter allele (induced variation). Thus, this approach speeds the assignment of mechanism(s) to natural variation in the germplasm. Indeed, one can consider this as a screen for epistatic or contingent gene action. MAGIC screens have identified loci from maize involved in the hypersensitive response (Chintamanani *et al*. 2010; Penning, Johal, and McMullen 2004; Olukolu *et al*. 2013, 2014, 2016) and plant development (Buescher *et al*. 2014), among others. In each case, in the absence of a mutant allele, no phenotype was previously associated to the modifier loci demonstrating the remarkable efficiency of this genetic screen to detect epistatic interactions between mutant and modifier alleles. Thus, this approach is a powerful way to both characterize cryptic variation in genomes and construct genetic pathways affecting phenotypes of interest.

The easy visual scoring and simplicity of quantifying chlorophyll make chlorophyll biosynthetic mutants an excellent reporter for MAGIC screens. Chlorophyll is a major component of central metabolism in plants, which can produce phototoxic intermediates during both synthesis (Hu *et al.* 1998; Huang *et al.* 2009) and breakdown (Gray *et al*. 1997; Gray *et al*. 2002; Mach *et al*. 2001; Yang et al. 2004), and its levels are carefully regulated in plants (Meskauskiene *et al.* 2001). Enzymatic conversion of protoporphyrin IX into magnesium protoporphyrin IX by Magnesium Chelatase (MgChl) is the first committed step in chlorophyll biosynthesis (Wettstein *et al.* 1995). MgChl is a hetero-oligomeric enzyme consisting of three subunits (I, D, and H) that are conserved from prokaryotes to plants. The MgChl subunit I is encoded by the *oil yellow1* (*oy1*; GRMZM2G419806) gene in maize and encodes the AAA^+^-type ATPase subunit that energizes the complex (Fodje *et al.* 2001). Weak loss-of-function alleles of *oy1* result in recessive yellow-green plants while complete loss-of-function alleles result in a recessive yellow seedling-lethal phenotype (Sawers *et al.* 2006). The semi-dominant *Oy1-N1989* allele carries a leucine (L) to phenylalanine (F) change at amino acid position 176 (L176F) that hinders the formation of a functional complex between the OY1 protein and other subunits of MgChl. Heterozygous plants carrying one *Oy1-N1989* allele and one wild-type allele are oil-yellow, but homozygous *Oy1-N1989* plants lack any MgChl activity, resulting in a recessive yellow seedling-lethal phenotype with no chlorophyll accumulation (Sawers *et al.* 2006). Consistent with the conservation of this protein complex, the orthologous L->F mutation (encoded by *Oy1-N1989*) was identified in barley (L161F) and also results in a recessive yellow seedling-lethal phenotype with no detectable chlorophyll in homozygous condition and a pale-green phenotype as a heterozygote (Hansson *et al.* 1999). The biochemical basis of this semi-dominant mutant allele was studied by creating a mutant MgChl subunit I in *Rhodobacter* (BCHI), with the orthologous amino acid change at position 111 of wild-type BCHI (Hansson *et al*. 2002). The L111F mutation converted BCHI into a competitive inhibitor of MgChl that reduced enzyme activity by 4-fold when mixed 1:1 with wild-type BCHI (Hansson *et al*. 2002; Lundqvist *et al*. 2013). This conserved leucine residue is between the ATP-binding fold of MgChl subunit I created by the Walker A and B motifs (Hansson *et al*. 2002; Sawers *et al*. 2006; Lundqvist *et al*. 2013), and its substitution with phenylalanine was deleterious to dephosphorylation activity. The ATPase activity of wild-type MgChl complex is directly proportional to the magnesium chelation reaction. However, complexes assembled from MgChl subunit I with the L->F change exhibit reduced ATPase activity and no ability to chelate Mg^2+^ ions into protoporphyrin IX (Hansson *et al*. 1999, 2002; Sawers *et al*. 2006). The absence of MgChl activity displayed by the mutant BCHI carrying L111F substitution (BCHI^L111F^) demonstrates that this amino acid change results in a dominant-negative subunit which disrupts the coupling of ATPase activity to magnesium chelation (Hansson *et al*. 2002; Lundqvist *et al*. 2013).

We screened maize germplasm for cryptic variation using the *Oy1-N1989* mutant as a dominant reporter for chlorophyll biosynthesis. We hypothesized that alteration in the quantity of biochemically active MgChl complex should read out as a change in chlorophyll content and plant color in heterozygous *Oy1-N1989* mutants. For instance, in a population of heterozygous *Oy1-N1989/oy1* F_1_ plants, an amino acid change in the wild-type OY1 protein that alters the dissociation constant (k_D_) of OY1 for the other protein subunits of MgChl should contribute to variance in chlorophyll biogenesis. Similarly, a cis-acting expression QTL (eQTL) that increases expression of the wild-type *oy1* allele should result in assembly of more active MgChl complexes and increase chlorophyll content in F_1_ mutant plants. Thus, chlorophyll content of *Oy1-N1989* mutants should be modulated by the stoichiometry of both the wild-type and mutant OY1 proteins present in the MgChl complex in heterozygous *Oy1-N1989/oy1* plants either by protein structure or abundance changes.

We introgressed the maize *Oy1-N1989* mutant allele into the B73 inbred background and maintained it as a heterozygote (*Oy1*-*N1989*/*oy1*:B73). While crossing this mutant to multiple backgrounds, we detected genetic variation in the *Oy1*-*N1989/oy1* mutant phenotype expression between the maize inbred lines B73 and Mo17. The phenotype of the *Oy1*-*N1989* mutant heterozygotes was suppressed in the B73 background. However, F_1_ hybrids with Mo17 dramatically enhanced it. In an attempt to determine the genetic basis of this modification, we carried out genetic mapping experiments using five F_1_ populations. In each of these mapping experiments, the *Oy1-N1989/oy1*:B73 parents were crossed with (1) the intermated B73 x Mo17 recombinant inbred lines (IBM), (2) B73 x Mo17 doubled haploid lines (Syn10), (3) Mo17 x B73 F_1_ hybrids, (4) B73 x Mo17 near-isogenic lines (BM-NILs), and (5) a genome-wide association mapping panel of diverse maize inbred lines (MDL). In each case, we identified a quantitative trait locus (QTL) of large effect on chromosome 10 linked to the *oy1* locus itself. In each of the B73 x Mo17 line-cross populations, the B73 wild-type allele at *oy1* suppressed the *Oy1-N1989/oy1* mutant phenotype. This QTL, which we call *very oil yellow1* (*vey1*), was not associated with changes in protein sequence at *oy1* and did not correlate with the chlorophyll content of the wild-type mapping parents. However, a cis-acting eQTL causing higher expression of the B73 allele of *oy1* in the IBM lines was consistent with the observed phenotypic variation in the mutant siblings. Consistently, the allele-specific expression at *oy1* in the mutant heterozygotes was biased towards the *Oy1-N1989* allele in the hybrids comprising wild-type *oy1* allele from Mo17. The inheritance of the traits, proposed allele expression bias at *oy1* due to a putative cis-acting regulatory element, implications of the discovered cryptic variation, and the utility of this study in general are discussed.

## Materials and Methods

### Plant materials

The *Oy1-N1989* mutant allele was acquired from the Maize Genetics Cooperation Stock Center (University of Illinois, Urbana-Champaign, IL). The original allele of the *Oy1-N1989* mutation was isolated from a *r1 c1* colorless synthetic stock of mixed parentage (G. Neuffer, personal communication). This allele was introgressed into B73 for eight generations by repeated backcrossing of B73 ear-parents with *Oy1-N1989/oy1* pollen-parents and is maintained as a heterozygote (*Oy1-N1989*/*oy1*:B73) by crossing to wild-type siblings.

Line-cross QTL mapping: For these experiments, *Oy1-N1989*/*oy1*:B73 plants were crossed as pollen-parents to 216 Intermated B73 x Mo17 recombinant inbred lines (IBM; Lee *et al*, 2002) and 251 Syn10 doubled haploid lines (Syn10; Hussain *et al*, 2007). The QTL validation was done using F_1_ progenies derived from the cross of 35 B73-Mo17 Near-Isogenic lines (BM-NILs) ears with pollen from *Oy1*-*N1989*/*oy1*:B73. These BM-NILs consisted of 22 B73-like NILs and 13 Mo17-like NILs with introgression of the reciprocal parental genome (B73 or Mo17) and were developed by three repeated backcrosses into recurrent parent followed by four to six generations of self-pollination (Eichten *et al.* 2011).

Genome-wide association (GWA) mapping: For this experiment, *Oy1*-*N1989*/*oy1*:B73 plants were crossed to 343 inbred lines that included 249 inbreds from maize association panel (Flint-Garcia *et al.* 2005), and 94 inbred lines that included 82 Expired Plant Variety Protections (ExPVP) lines from the Ames panel (Romay *et al.* 2013). Pollen from *Oy1*-*N1989*/*oy1*:B73 plants were used for these crosses except for the popcorn lines in the maize association panel, where *Oy1*-*N1989*/*oy1*:B73 plants were used as an ear-parent to avoid the crossing barrier affected by gametophytic factor GA1-S (first described by Correns in 1902; Mangelsdorf and Jones 1926; Lauter *et al.* 2017). This panel of 343 inbred lines is referred to as maize diversity lines (MDL). The full list of IBM, Syn10, BM-NILs, and MDL used to develop F_1_ hybrid populations are provided in **Tables S1-S4**.

### Field trials

All field experiments were performed at the Purdue Agronomy Center for Research and Education (ACRE) in West Lafayette, Indiana. All F_1_ populations described below were planted as a single plot of 12-16 plants that segregated for both mutant and wild-type siblings. Plots were sown in a 3.84 m long row with the inter-row spacing of 0.79 meters and an alley space of 0.79 meters. No irrigation was applied during the entire crop season as rainfalls were uniformly distributed for satisfactory plant growth. Conventional fertilizer, pest and weed control practices for growing field maize in Indiana were followed. Progenies of *Oy1*-*N1989*/*oy1*:B73 pollen-parents crossed with B73 and Mo17 were planted as parental checks in each block of every experiment. The testcross F_1_ populations with IBM were evaluated in a single replication in the summer of 2013 with each range treated as a block. In 2016, the testcross F_1_ populations with Syn10 lines were evaluated as two replications in a randomized complete block design (RCBD) with each range divided into two blocks. The testcross F_1_ progenies with BM-NILs and parents (B73 and Mo17) were planted in a RCBD with five replications in 2016. In the same year, F_1_ populations with MDL were also evaluated with three replications planted in a RCBD. Each replication of MDL F_1_ population was divided into ten blocks of the same size, and parental checks were randomized within each block.

### Phenotyping and data collection

Maize seedlings used for destructive chlorophyll quantification were grown under greenhouse conditions using mogul base high-pressure sodium lamps (1000 Watts) as the supplemental light source for L:D cycle of 16:8 and temperature around 28°C (day-time) and 20°C (night-time). Destructive chlorophyll measurements were performed on the fresh weight basis in 80% acetone solution using a UV-VIS spectroscopic method described in Lichtenthaler & Buschmann, 2001.

For the field-grown experiments, mutant siblings in the suppressing genetic backgrounds were tagged at knee height stage with plastic tags so that they can be easily distinguished from the wild-type siblings for trait measurements at later developmental stages in the season. All the F_1_ families segregated for the mutant (*Oy1*-*N1989*/*oy1*) and wild-type (*oy1*/*oy1*) siblings in approximately 1:1 fashion. For each F_1_ family, two to four plants of each phenotypic class were picked at random for trait measurements. Non-destructive chlorophyll content in the maize leaves was approximated using a chlorophyll content meter model CCM-200 plus (Opti-Sciences, Inc., Hudson, NH) and the measurements were expressed as chlorophyll content index (CCM). Measurements were taken on the leaf lamina of the top fully expanded leaf at two time points. First CCM measurements were taken at 25-30 days after sowing (expressed as CCMI) and the second at 45-50 days after sowing (expressed as CCMII). For each trait, measurements were performed on both mutant (reported with a prefix MT) and wild-type (reported with a prefix WT) siblings. Besides using primary trait measurements of CCMI and CCMII on mutant and wild-type siblings, indirect CCM measurements were also calculated and expressed as ratios (MT/WT) and differences (WT-MT) of CCMI and CCMII. Phenotypic data of all the CCM traits in the F_1_ populations with both bi-parental populations, BM-NILs and MDL are provided in **Tables S1-S4**.

### Public/open-access genotypic and gene expression data

Public marker data for the IBM was obtained from www.maizegdb.org (Sen *et al.* 2010). A total of 2,178 retrieved markers were reduced to 2,156 after the removal of markers with the duplicate assessment. Approximately 13.3% of the marker data were missing. The reads per kilobase of transcript per million mapped reads (RPKM) for the *oy1* locus were obtained from a public repository of the National Science Foundation grant (GEPR: Genomic Analyses of shoot meristem function in maize; NSF DBI-0820610; https://ftp.maizegdb.org/MaizeGDB/FTP/shoot_apical_meristem_data_scanlon_lab/). These data consist of normalized read counts (expressed as RPKM) of the maize genes from the transcriptome of shoot apex of 14 days old IBM seedlings.

Marker data for Syn10 lines was obtained from Liu *et al*. 2015. The Syn10 lines were genotyped at 6611 positions (B73 RefGenv2) with SNP markers covering all ten chromosomes of maize. The entire set of markers were used for linkage analysis as there was no missing data. All B73-Mo17 NILs in both the B73 and Mo17 recurrent backgrounds that had introgression of the critical region from the opposite parent were selected for QTL validation. Genotyping data of the BM-NILs to choose informative lines and perform QTL validation was obtained from Eichten *et al*. 2011.

Genotypes for the MDL used in this study to perform GWA were obtained from third generation maize haplotypes (HapMap3) described in Bukowski *et al.* 2018. We identified 305 common inbred lines that were part of HapMap3. Briefly, HapMap3 consists of over 83 million variant sites across ten chromosomes of maize that are anchored to B73 version 3 assembly. After obtaining the genotypic data of 305 common inbred lines, variant sites were filtered for ≤10 percent missing data and minor allele frequency of ≥0.05 using VCFtools (Danecek *et al.* 2011). The filtered SNP dataset was used for GWA analyses. A summary of variant sites before and after the filtering procedure are listed in **Table S5.** A reduced set of genotypes for the same set of 305 accessions were obtained from the maize inbred lines described in Romay *et al*. 2013. These genotypes consist of 681 257 SNPs (physical positions from B73 RefGenv2) obtained using a GBS protocol (Elshire *et al.* 2011) covering all ten chromosomes of maize. This marker dataset was filtered for ≤10 percent missing data and minor allele frequency of ≥0.05 using TASSEL (Bradbury *et al.* 2007), reducing the marker number to 150 920 SNPs. This genotypic dataset was solely used to compute principal components (PC) and a kinship matrix to control for population structure and familial relatedness in a unified mixed linear model, respectively (Yu *et al.* 2006). GWA analyses using the reduced marker dataset (Romay *et al*. 2013) on the full set of MDL found similar results to the one by the larger marker dataset (HapMap3) on the reduced set of MDL (data not shown). To test for cis-eQTL at the *oy1* locus in the maize diversity lines used for GWA mapping, normalized count of OY1 expression derived from the germinating seedling shoots was obtained from http://www.cyverse.org (Kremling *et al.* 2018).

### Statistical analyses

QTL mapping: Line-cross phenotypes and markers were used to detect and localize QTL using the R/QTL package version 1.40-8 (Broman *et al*. 2003). Trait means were used for the QTL analyses. Single interval mapping (SIM) was used for all traits, although composite interval mapping (CIM) was carried out with remarkably similar results (data not shown). Statistical thresholds were computed by 1000 permutations (Churchill and Doerge 1994) of each trait in both bi-parental F_1_ populations.

Genome-wide association study (GWAS): Preliminary data analysis was done using JMP 13.0 (SAS Institute Inc. 2016). Statistical corrections on the raw phenotypic data were performed by determining the most significant terms in the model using analysis of variance (Fisher 1921). Genotype and replication were used as a random effect in a linear mixed-effects model built in the lme4 package (Bates *et al*. 2015) implemented in R (R core team, 2014) to calculate the best linear unbiased predictor (BLUP) for each trait. Broad-sense heritability (line mean basis) were calculated using BLUP values, using the method described by Lin and Allaire 1977. BLUP estimates for each trait were used to perform GWAS. GWAS was done using a compressed mixed linear model implemented in R package GAPIT (Lipka *et al*. 2012; Zhang *et al*. 2010). HapMap3 SNPs were used to calculate genotype to phenotype associations. As explained before, kinship and population structure estimates were obtained for the same population using the second subset of 150 920 SNPs to correct for spurious associations. The Bonferroni correction and false discovery rate (FDR) adjustments were used to compute a statistical threshold for the positive association to further control for false positive assessment of associations (Holm 1979; Benjamini and Hochberg 1995).

### Molecular Analyses

Genotyping: The recombinants in selected Syn10 lines and the BC_1_F_1_ population were detected using three PCR-based markers. Two markers detecting insertion polymorphisms flanking the *oy1* locus and one dCAPS marker at *oy1* locus were designed for this purpose. Genotyping at insertion-deletion (indel) marker *ftcl1* (flanking an indel polymorphism in intron 4 of *ftcl1*; GRMZM2G001904) was performed with forward primer 5’- GCAGAGCTGGAATATGGAATGC-3’ and reverse primer 5’-GATGACCTGAGTAGGGGTGC -3’. Genotyping at indel marker *gfa2* (flanking an indel polymorphism in the intron of *gfa2*; GRMZM2G118316) was performed with forward primer 5’- ACGGCTCCAAAAGTCGTGTA -3’ and reverse primer 5’-ATGGATGGGGTCAGGAAAGC -3’. A polymorphic SNP in the second intron at *oy1* was used to design a dCAPS forward primer 5’- CGCCCCCGTTCTCCAATCCTGC -3’ and a gene-specific reverse primer 5’- GACCTCGGGGCCCATGACCT -3’ (http://helix.wustl.edu/dcaps/dcaps.html; Neff *et al*. 2002). The PCR products from polymerization reactions with the dCAPS oligonucleotide at *oy1* were digested by *Pst* I restriction endonuclease (New England Biolabs, MA, USA) and resolved on 3.5% agarose gel.

Allele-specific expression analyses: Allelic bias at transcriptional level was quantified using the third leaf of maize seedlings at the V3 developmental stage. Total RNA was extracted using a modified Phenol/Lithium chloride protocol (Eggermont *et al*. 1996). The modification involved grinding of plant tissue in pestle and mortar into a fine powder using liquid nitrogen, instead of quartz sand and glass beads in the original protocol. Total RNA was subjected to DNase I treatment using Invitrogen Turbo DNA-free kit (Catalog#AM1907, Life Technologies, Carlsbad, CA) and 1 µg of DNase treated RNA from each sample was converted to cDNA using oligo dT primers and a recombinant M-MLV reverse transcriptase provided in iScript^TM^ Select cDNA synthesis kit (catalog#170-8896, Bio-Rad, Hercules, CA) according to the manufacturer’s recommendations. Besides the cDNA samples, genomic DNA samples were also prepared as a control to test the sensitivity of the assay. Genomic DNA controls included a 1:1 (F_1_ hybrid), 1:2, and 2:1 mixture of B73 and Mo17. PCR was conducted using gene-specific forward primer 5’- CAACGTCATCGACCCCAAGA -3’ and reverse primer 5’- GGTTACCAGAGCCGATGAGG -3’ for 30 cycles (94° for 30s, 56° for 30s, 72 ° for 30s and final extension for 2 minutes) to amplify the OY1 gene product. These primers flank SNP252 (C->T), which is the causative mutation of *Oy1*-*N1989*, and SNP317 (C->T) which is polymorphic between B73 and Mo17 but monomorphic between the B73 and *Oy1*-*N1989* genetic backgrounds. Corresponding PCR products were used to generate sequencing libraries using transposon-mediated library construction with the Illumina Nextera^®^ DNA library preparation kit, and sequence data were generated on a MiSeq instrument (Illumina, San Diego, CA) at the Purdue Genomics Core Facility. The SNP variation and read counts were decoded from the sequenced PCR amplicons by alignment of the quality controlled reads to the *oy1* reference allele from B73 using the BBMap (Bushnell 2014) and the GATK packages (DePristo *et al.* 2011). Additional analysis was performed using IGV (Robinson *et al.* 2011) to manually quality-check the alignments and SNP calls. Read counts at polymorphic sites obtained from GATK was used to calculate allele-specific expression. Genomic DNA control samples showed bias in the read counts in a dosage-dependent manner. DNA from F_1_ hybrids between B73 and Mo17 resulted in 1:1 reads at *oy1* demonstrating no bias in the assay to quantify expression.

OY1 sequencing: The coding sequence of the *oy1* locus from B73, Mo17, W22, and *Oy1*-*N1989* homozygous seedlings was obtained from the genomic DNA using PCR amplification. For the rest of the maize inbred lines, the *oy1* locus was amplified from cDNA synthesized from total RNA derived from the shoot tissue of 14 days old maize seedlings. PCR amplification of *oy1* locus from genomic DNA was performed using four primer pairs: (a) forward primer Oy1-FP1 5’- GCAAGCATGTTGGGCACAGCG -3’ and reverse primer Oy1-RP12 5’- GGGCGGCGGGATTGGAGAAC -3’, (b) forward primer Oy1-FP5 5’- GGTGGAGAGGGAGGGTATCT -3’ and reverse primer Oy1-RP6 5’- GGACCGAGGAAATACTTCCG -3’, (c) forward primer Oy1-F8 5’- ATGCCCCTTCTTCCTCTCCT -3’ and reverse primer Oy1-R8 5’- CGCCTTCTCGATGTCAATGG -3’, (d) forward primer Oy1-F9 5’- GGCACCATTGACATCGAGAA -3’ and reverse primer Oy1-R9 5’- GCTGTCCCTTCCTTTCAACG -3’. PCR amplification of OY1 transcripts from cDNA was performed using all primer pairs except Oy1-FP1/RP12. The PCR products from these samples were sequenced either using Sanger or Illumina sequencing. For Sanger sequencing, amplified PCR products were cleaned using homemade filters with sterile cotton plug and Bio-Gel^®^ P-30 gel (Bio-Rad, Hercules, CA) to remove unused dNTPs and primers. Cleaned PCR products were used to perform a cycle reaction using Big Dye version 3.1 chemistry (Applied Biosystems, Waltham, MA) and run on ABI 3730XL sequencer by Purdue genomics core facility. Read with high-quality base pairs from Sanger sequencing were aligned using ClustalW (Thompson *et al*. 1994). Illumina sequencing was performed as described above except in this case paired-end reads were aligned to the B73 reference of *oy1* gene using bwa version 0.7.12 (Li and Durbin 2009) and variant calling was done using Samtools (Li *et al*. 2009).

## Results

### Mo17 encodes an enhancer of the semi-dominant mutant allele ***Oy1*-*N1989* of maize**

The *Oy1-N1989* allele was recovered from a nitrosoguanidine mutant population in mixed genetic background. The molecular nature of the mutation is a single non-synonymous base pair change (Sawers *et al.* 2006). Heterozygous *Oy1-N1989* plants have the eponymous oil-yellow color but are reasonably vigorous and produce both ears and tassels. During introgression of the semi-dominant *Oy1-N1989* allele into B73 and Mo17 inbred backgrounds, we observed a dramatic suppression of the mutant phenotype in F_1_ crosses of the mutant stock (obtained from Maize COOP) to the B73 background. In contrast, crosses to Mo17 enhanced the mutant phenotype. The difference in phenotype expression was stable and persisted in both genetic backgrounds through all six backcross generations observed to date. To further explore and quantify this suppression, B73, Mo17, as well as *Oy1-N1989/oy1*:B73, crossed with each of these inbred lines were grown to the V3 stage in the greenhouse. To improve upon our visual assessment of leaf color and provide quantitation, optical absorbance was measured using a Chlorophyll Content Meter-200 plus (CCM; Opti-Sciences, Inc), a hand-held LED-based instrument. CCM is predicted to strongly correlate with chlorophyll and carotenoid contents. To confirm that our rapid phenotyping with the CCM would accurately assess chlorophyll levels, we measured these pigments using a UV-VIS spectrophotometer following destructive sampling of the same leaves used for CCM measurements. The non-destructive CCM measurements and destructive pigment quantification using UV-VIS protocol displayed a strong positive correlation with an R^2^ value of 0.94 for chl*a*, chl*b*, and total chlorophyll (**Figure S1**). Given this high correlation of maize leaf greenness between the rapid measurement using CCM-200 plus instrument and absolute pigment contents quantified using UV-VIS spectrophotometer (destructive sampling), we performed all chlorophyll measurements of *Oy1-N1989* enhancement discussed in later results using CCM values.

In the greenhouse grown seedlings, we observed no visible chlorophyll accumulation in the *Oy1*-*N1989* homozygotes using CCM or spectrophotometric method. In wild-type plants, CCM measurements were slightly higher in B73 than Mo17, but the spectrophotometric method did not identify any significant difference in the amount of chlorophyll *a* (chl*a*), chlorophyll *b* (chl*b*), total chlorophyll, or carotenoids between these two genotypes (**Table S6**). We detected a mild parent-of-origin-effect for both CCM and absolute amounts of chl*a*, chl*b*, total chlorophyll, and carotenoids in the wild-type siblings of our F_1_ crosses. These plants had slightly higher chlorophyll accumulation when B73 was used as the pollen parent (**Table S6**). However, no such effect of parent-of-origin was observed for the mutant heterozygotes (*Oy1-N1989/oy1*) and reciprocal hybrid combinations of crosses between *Oy1-N1989/oy1*:B73 and Mo17 were indistinguishable. Further, both CCM and absolute chlorophyll contents were higher in the *Oy1-N1989/oy1*:B73 plants compared to the mutants in B73 x Mo17 hybrid background. Thus, there was a strong effect of genetics on chlorophyll pigment variation in mutants, that went opposite to predictions for hybrid vigor.

Heterozygous maize plants encoding the *Oy1-N1989* allele display reduced chlorophyll pigment abundance compared to the wild-type siblings, resulting in a yellow-green whole plant phenotype due to reduced MgChl and ATPase activity (Sawers *et al.* 2006). We tested the progenies from the crosses of *Oy1-N1989/oy1*:B73 with B73 and Mo17 inbred lines in the field. Consistent with our previous observation, B73 inbred background resulted in substantially greener mutant heterozygotes (*Oy1-N1989*/*oy1*) than mutant siblings in the Mo17 x B73 F_1_ hybrid background (**Figure 1, Table S7**). The increased severity of *Oy1-N1989* heterozygotes in the Mo17 genetic background was observed even after six backcrosses (**Figure S2**). No suppressed mutant plants were observed during any generation of backcrossing into Mo17 (data not shown). Thus, these results demonstrate the profound negative impact of the *Oy1-N1989* allele on chlorophyll pigment accumulation and the dramatic differential suppression response of this allele by B73 and Mo17 genetic backgrounds.

**Figure 1.**
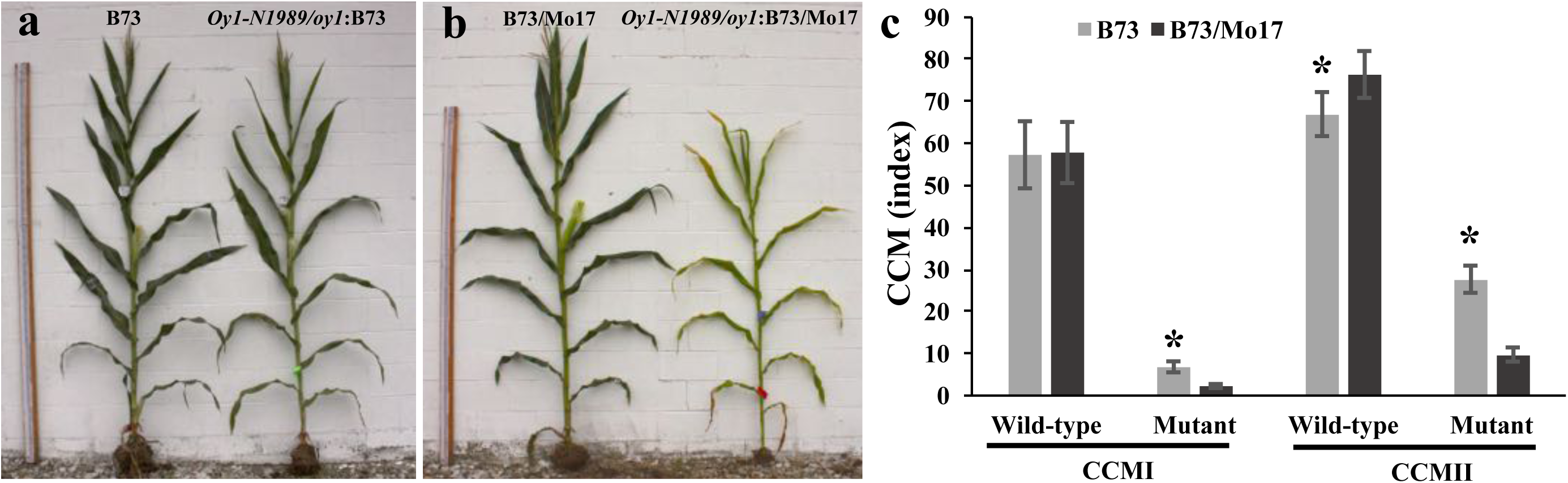
The chlorophyll pigment accumulation differs in severity for *Oy1-N1989/oy1* heterozygotes in the B73 and B73 x Mo17 hybrid backgrounds. The representative wild-type (*oy1/oy1*) and mutant (*Oy1-N1989/oy1*) sibling from (a) B73 x *Oy1-N1989/oy1*:B73 and (b) Mo17 x *Oy1-N1989/oy1*:B73 F1 crosses in the field-grown plants. The measuring stick in panel a and b is 243 cm. (c) Non-destructive chlorophyll approximation in mutant and wild-type siblings at an ~3 weeks (CCMI) and ~6 weeks (CCMII) after planting; data for each class of genotype is derived from 39 replications planted in RCBD. The asterisk indicates statistical significance between the means in each genotype category at p <0.01 determined using student’s t-test.

### *vey1* maps to a single locus that co-segregates with the *oy1* allele of Mo17 in DH, RIL, BC_1_F_1_ and NIL families derived from B73 and Mo17

To identify the genetic basis of the suppression of heterozygotes with the *Oy1-N1989* allele in B73, we performed a series of crosses to four mapping populations. In each case, we crossed a B73 line into which we have introgressed *Oy1-N1989* allele in heterozygous condition (*Oy1*-*N1989*/*oy1*:B73) as a pollen-parent to a population of recombinant lines as ear-parent (**Figure 2**). We chose two public populations to map all modifiers altering the severity of the *Oy1*-*N1989* phenotypes. The IBM is a publicly available RIL population that has been extensively used by the maize community for a variety of mapping experiments (Lee *et al*. 2002). A second intermated B73 x Mo17 population Syn10 is derived from ten rounds of intermating followed by fixing alleles using double haploid process (Hussain *et al*. 2007). IBM and Syn10 differ in the number of rounds of intermating, and therefore vary in the number of recombinants captured and genetic resolution of trait localization. Each F_1_ progeny of the testcross with *Oy1-N1989/oy1*:B73 pollen-parent segregated approximately 1:1 for wild-type (*oy1*/*oy1*) and mutant heterozygote (*Oy1*-*N1989*/*oy1*) in the hybrid genetic background with B73. In the mutant heterozygote siblings of both (IBM and Syn10) F_1_ populations, chlorophyll approximation using CCM measured at an early (CCMI) and late (CCMII) developmental stages displayed bimodal trait distributions (**Figures 3a and 3b; Figures S3 and S4**), and there was no overlap between the wild-type and mutant CCM distribution (**Figures S5a and S5b**). Moreover, mutant siblings alone in these F_1_ populations with IBM and Syn10 displayed bimodality for CCM measurements, suggesting segregation of a single, effectively Mendelian, suppressor locus (**Figures 3a and 3b**). We name this suppressor locus *very oil yellow1* (*vey1*). CCM values collected at two different times (CCMI and CCMII) in the wild-type F_1_ siblings show positive correlations in both F_1_ populations (**Tables S8 and S9; Figures S3 and S4**). Similar trend was also observed for the CCMI and CCMII in the mutant F_1_ siblings. However, CCM measurements in the wild-type and mutant displayed statistically insignificant correlations with each other. This suggests that higher chlorophyll accumulation in the mutant siblings is not predicted by the amount of chlorophyll accumulation in the wild-type siblings. To control for variation in CCM observed due to the genetic potential of each line that was independent of the *Oy1-N1989* modification, we divided the mutant CCM trait values by the congenic wild-type sibling CCM values to derive ratio for both time points. We also calculated differences between congenic wild-type and mutant CCM values. Each of the direct measurements, as well as the ratio-metric and difference values, were used as phenotypes to detect and localize QTL.

QTL mapping was carried out on all traits using single interval mapping by the EM algorithm as implemented in R/qtl (Broman *et al.* 2003). Summary of the peak positions of all QTLs passing a permutation computed threshold are presented in **Table S10** for the IBM F_1_ populations and **Table S11** for the Syn10 F_1_ populations. All mutant CCM traits, all mutant to wild-type CCM ratios, and all differences between mutant and wild-type CCM measurements identified *vey1* as a QTL of large effect on chromosome 10. A plot of the log_10_ of odds (LOD) score and permutation calculated threshold for CCMII from the mutant siblings in the Syn10 F_1_ population is plotted in **Figures 3c and 3d**. Other mutant-related traits in Syn10 and IBM F_1_ populations produced similar plots (data not shown). The magnitude of the effects of *vey1* genetic position on all of the traits in the mutant siblings, particularly the effects on CCMI and CCMII, were consistent with the prediction of a single Mendelian locus controlling the majority of the phenotypic variance in these traits based on trait distributions (**Figure 3a and 3b**). Only one additional minor effect QTL was identified that influenced the wild-type CCMI. This QTL was only found in the IBM F_1_ population (**Table S10**). No QTL affecting the chlorophyll accumulation of wild-type siblings were detected at the position of *vey1,* or anywhere else in the genome. Interestingly, *vey1* mapped to the same genetic position as *oy1* locus itself (**Figures 3e and 3f**). These analyses indicate that the *vey1* QTL encodes a single locus with an effect contingent upon the allelic status at *oy1* responsible for the suppression of *Oy1-N1989* in these populations.

**Figure 2.**
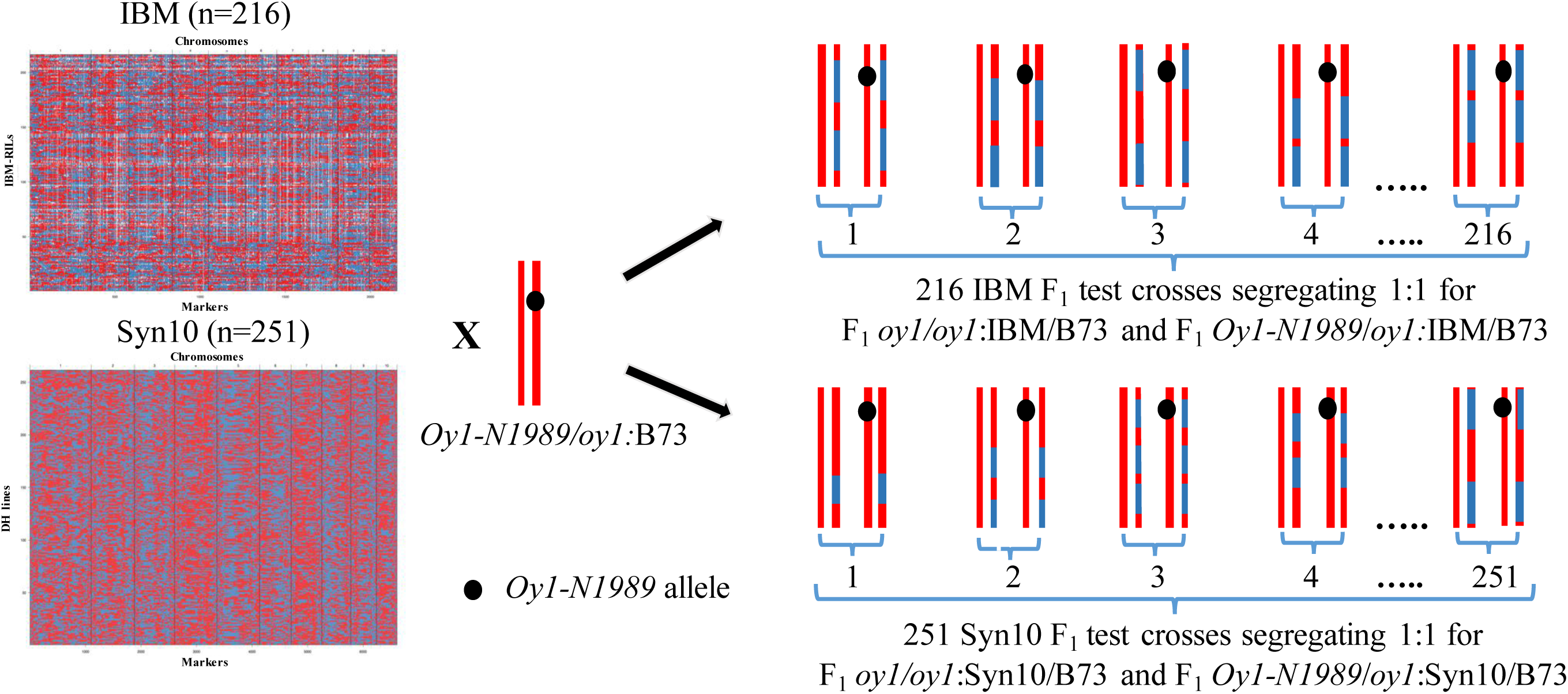
The crossing scheme used to map *Oy1-N1989* enhancer/suppressor loci in IBM and Syn10 populations. Red, blue and white colors indicate B73, Mo17, and missing genotypes. The heterozygous tester (*Oy1-N1989/oy1*:B73) shows chromosome 10 with a black dot indicating *Oy1-N1989* mutant allele. The F_1_ progenies depicted here shows a hypothetical state of chromosome 10 for each F_1_ progeny showing segregation of wild-type and mutant (with the black spot) siblings.

**Figure 3.**
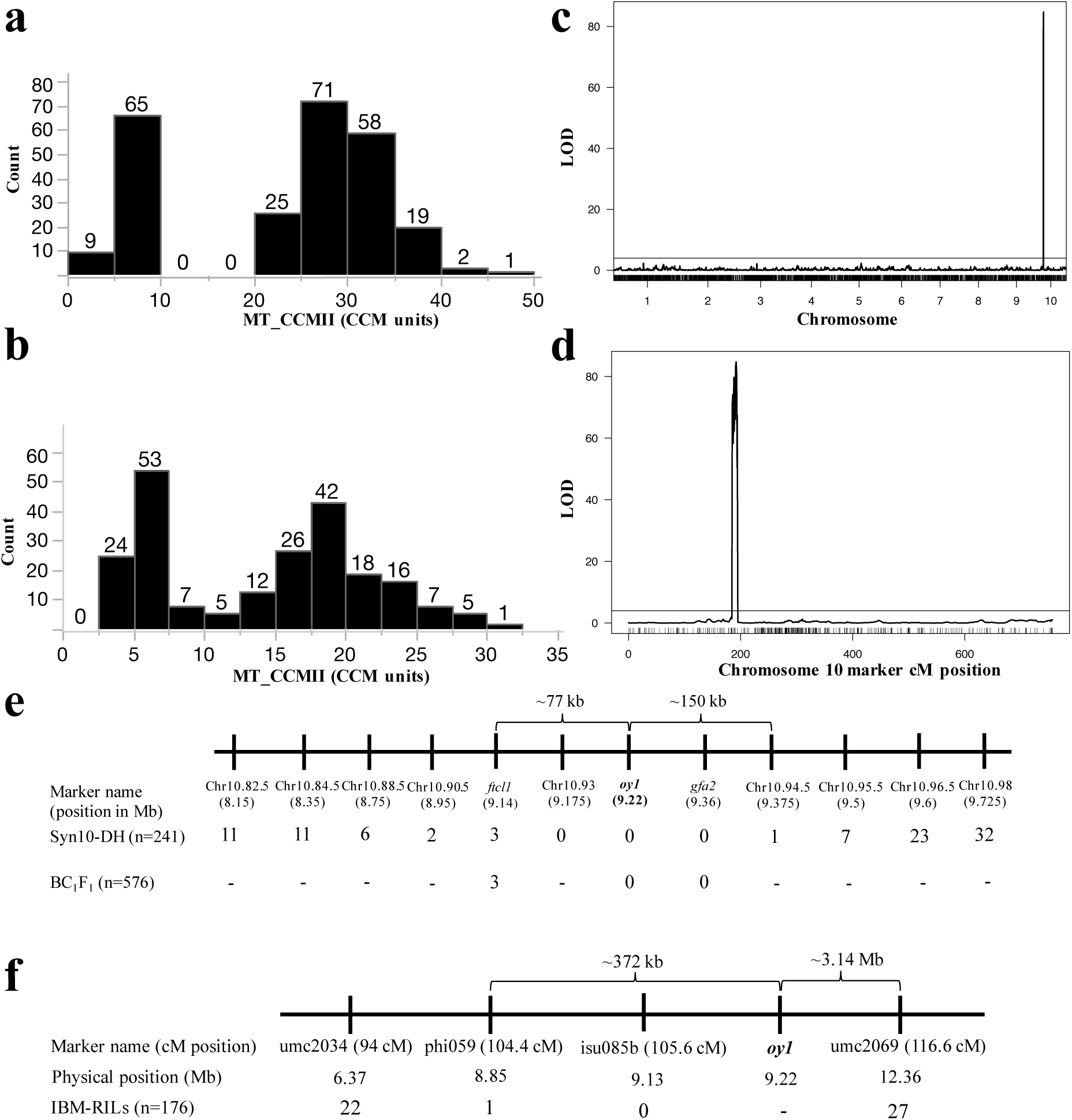
The phenotypic distribution, QTL analysis, and fine mapping results of MT_CCMII trait. The distribution of MT_CCMII in (a) Syn10 x *Oy1-N1989/oy1*:B73 F_1_ population and (b) IBM x *Oy1-N1989/oy1*:B73 F_1_ population. (c) Genomewide QTL plot of MT_CCMII in Syn10 x Oy1- N1989/oy1:B73 F_1_ population. The x-axis indicates the chromosome number and Y-axis indicates the logarithm to the base 10 of odds (LOD) of tested markers. Black horizontal bar indicates the permutation testing-based threshold to declare statistical significance of the QTL. (d) Close-up view of the vey1 locus on chromosome 10. The x-axis indicates the centiMorgan (cM) position of the molecular markers. Recombinants detected in F_1_ crosses of *Oy1-N1989/oy1*:B73 as pollen-parent with (e) Syn10 lines and B73 x Mo17 F_1_ (BC_1_F_1_), and (f) IBM. A number at a given marker and population intersection in (e) and (f) indicates the total number of recombinants between the respective marker genotype and phenotype; hyphen denotes no genotyping. dCAPS marker at *oy1* is highlighted in bold.

The identification of the *vey1* modifier as a single Mendelian locus of large effect in the presence of the *Oy1-N1989* allele suggested that we could take a similar approach to localization as for a previous metabolic QTL (Li *et al*. 2014). Thus, we classified each F_1_ mutant hybrid in Syn10 F_1_ population as high or low CCMII by using the bimodal distribution to assign lines into phenotypic categories. We then compared the marker genotypes at each marker under *vey1* with the phenotypic categories. **Table S12** shows the mutant trait values, marker genotypes, and phenotypic categories for the nine F_1_ Syn10 lines with recombinants within the *vey1* region (between the flanking markers 10.90.5 and 10.95.5). Because of the high penetrance of the CCM trait, we interpret a discordance between the marker genotype and F_1_ mutant phenotype for high and low CCMII categorization as recombination between *vey1* and that marker. A summary of the recombinant data across the *vey1* flanking markers and additional markers with the physical position of each marker annotated by the number of instances of discordance between the markers and the phenotypic class is presented in **Figures 3e and 3f**. Genotypes at marker 10.93 perfectly predicted CCMII trait expression in Syn10 F_1_ mutant siblings. Recombinants separated the trait outcome from the marker genotype in one Syn10 line at marker 10.94.5 and two Syn10 lines at 10.90.5, indicating that the QTL resides in ∼227kb interval between markers 10.94.5 and 10.90.5 (**Figure 3e**). This physical position includes the *oy1* locus itself, suggesting that *vey1* may be encoded by the Mo17 allele of *oy1* which enhances the impact of the *Oy1-N1989* allele. In the three Syn10 lines that contained recombination within this critical region, the genotype at *oy1* matched mutant CCM trait values perfectly. To improve the resolution of *vey1* localization we developed additional markers in the region. The genotypes at polymorphic indel markers were determined for the Syn10 recombinants. One marker was encoded by the *ftcl1* locus, two genes towards the telomere from *oy1* and the second, was encoded by *gfa2*, one gene towards the centromere. No recombinants were detected between an indel marker in the proximal end of the *gfa2* gene and *vey1*, although marker 10.94.5 from the Syn10 population is at the distal end of the *gfa2* gene and did show one recombinant (**Figure 3e**). Three recombinants were identified that separated the *ftcl1* marker from the *vey1* QTL. As a result, *vey1* could be encoded by the genomic region between *ftcl1* and *gfa2*. This region includes *oy1, ereb28* (GRMZM2G544539), the small regions of *gfa2* (from marker 10.94.5), and *ftcl1* proximate to *oy1*.

A similar fine mapping approach was adopted for the data from the F_1_ crosses to the IBM. Mutant CCMII readings were used to bin IBM F_1_ population into suppressed and enhanced categories. The number of cases of discordance between each marker genotype in the *vey1* region and the F_1_ mutant phenotypic class is summarized in **Figure 3f**. For all IBM with unambiguous genotypes, the *Oy1-N1989* suppression or enhancement phenotype was correctly predicted by marker genotypes at isu085b. A single recombinant between phenotype and genotype was identified for the flanking marker phi059 which is ∼372 kb from the *oy1* locus (towards the telomere). Twenty-seven recombinants between the phenotype and genotype were noted for the marker umc2069 which is ∼3.14 Mb from *oy1* (towards the centromere). This analysis identified an ∼3.51 Mbp window in IBM containing *vey1* on chromosome 10 providing a confirmation of the Syn10 results with no additional resolution. We attempted to generate additional recombinants by generating a population of ∼1100 BC_1_F_1_ plants from (B73 x Mo17) x *Oy1-N1989/oy1*:B73 crosses. All the mutant siblings from this population (n=576) were separated into suppressed and enhanced phenotype categories by CCM readings and genotyped at *ftcl1, oy1*, and *gfa2*. In a sample of 576 mutants, we identified three recombinants between *vey1* and *ftcl1* indel marker, whereas, no recombinant at *oy1* and *gfa2* were detected (**Figure 3e**). Thus, we could not further narrow down this QTL interval.

To validate the *vey1* locus, we made crosses between *Oy1-N1989/oy1* in the B73 background and a series of BM-NILs that contained the *vey1* QTL region introgressed into a homogeneous background of either B73 or Mo17. These BM-NILs also displayed the bimodal effect observed in the QTL experiment, but now without additional segregating B73 and Mo17 alleles. This bimodality was still visible in crosses of *Oy1-N1989* mutant to NILs that used B73 as the recurrent parent to test the *vey1* QTL in an otherwise inbred B73 background demonstrating that a hybrid background was not necessary for *vey1* expression (**Figure S6 and Table S3**). We also observed an increase in the mutant CCM in F_1_ hybrids of *Oy1-N1989/oy1*:B73 with NILs that used Mo17 as a recurrent parent but had B73 introgression at *vey1* locus. Thus, our NIL data confirm the expectation of the QTL mapping and validates the existence of the *vey1* modifier. The recombinants in the BM-NILs were sufficient to identify an ∼3.01 Mb region that must contain the *vey1* QTL, but this did not further narrow the region from the four gene window which included the *oy1* locus itself, identified in the Syn10 F_1_ population.

Thus, the formal list of candidate genes for the *vey1* QTL is the *oy1* gene itself and the three most closely linked loci. Locus *ftcl1* is annotated as a 5-formyl tetrahydrofolate cyclo-ligase1, which is involved in folate metabolism. The ortholog of *Zmftcl1* (∼62% protein identity) from *Arabidopsis thaliana* has been shown to be localized in the chloroplast and T-DNA insertion knockouts are embryo lethal (Pribat *et al.* 2011). The maize gene *gfa2* is uncharacterized, but mutation of the *Arabidopsis* ortholog caused defects in megagametogenesis including failures of polar nuclear fusion in the female gametophyte and synergid cell-death at fertilization (Christensen *et al*. 2002; Christensen, Subramanian, and Drews 1998). The third linked gene, *ereb28* (Apetela2-Ethylene Responsive Element Binding Protein-transcription factor 28) exhibits poor homology to other plant species but has a highly conserved AP2/EREB domain. This gene has a very low expression level and is localized only to the root tissue of maize (http://www.maizegdb.org/gene_center/gene?id=GRMZM2G544539#rnaseq).

### Controlling for the *vey1* QTL neither detected additional epistatic interactions with *vey1* nor *Oy1-N1989* phenotype expression

The non-normality of some of the trait distributions and apparent thresholds prompted us to explore additional QTL models. No additional QTL were recovered by implementing two-part threshold models (Broman *et al.* 2003) for any of the traits (data not shown). This observation is consistent with a single major QTL affecting the non-normality in the phenotypic data and normal distribution of residuals remaining after single marker regressions (data not shown). Similarly, two-way scans of the genome also failed to detect any statistically significant genetic interactions. It is worth noting that in both the IBM and the Syn10, the region encoding *vey1* exhibited substantial segregation distortion with the B73:Mo17 alleles present at 120:72 in the IBM and 175:76 in the Syn10 population. This uneven sample size will reduce the power to detect epistasis with *vey1* but would not limit the detection of additional unlinked epistatic modifiers of *Oy1-N1989*.

We used the top marker at *vey1* as a covariate to control for the contribution of this allele to phenotypic variation and performed a one-dimensional scan of the genome (Broman *et al.* 2003). In our previous naïve one-dimensional scans, the large effect of *vey1* partitioned into the error term and might reduce our power to detect additional unlinked QTL(s). By adding a marker linked to *vey1* as a covariate, this term will capture the variance explained by *vey1* and should improve detection of additional QTL(s) of presumably smaller effect. Use of *vey1* linked marker as a covariate, in both IBM and Syn10 F_1_ populations, did not identify any additional QTL for any trait (data not shown). Thus, modification of the *Oy1-N1989* phenotype by *vey1* was inherited as a single QTL, acting alone.

### GWAS for chlorophyll content in maize diversity lines and ***Oy1-N1989*/*oy1* F_1_ genotypes identifies *vey1***

We undertook GWA mapping of *Oy1-N1989* severity to search for additional loci and potentially identify recombinants at *vey1*. A population of 343 lines including members of the maize association panel (Flint-Garcia *et al.* 2005) and the Ames panel including ExPVPs (Romay *et al.* 2013) were crossed to *Oy1-N1989/oy1*:B73. This procedure generated MDL x *Oy1-N1989/oy1*:B73 F_1_ populations segregating 1:1 for mutant and wild-type siblings in hybrid genetic background. There was total separation between mutant and wild-type siblings in the MDL F_1_ populations for the CCMI and CCMII traits (**Figure S5c**). Additionally, dramatically enhanced mutant F_1_ families, similar to the F_1_ progeny of Mo17 x *Oy1*-*N1989*/*oy1*:B73, were present within the MDL F_1_ populations (**Figure S7**). Pairwise correlations of the CCM trait measurements at two-time points (CCMI and CCMII) in the wild-type siblings displayed statistically significant positive relationship (**Table S13**). CCM traits were much more strongly correlated in the mutant F_1_ siblings, similar to the B73 x Mo17 F_1_ populations. However, weak positive correlations were also observed between mutant and wild-type CCM measurements in the MDL F_1_ populations. Broad-sense heritability estimates were also calculated in the MDL F_1_ population. The chlorophyll estimates of the leaves (CCMI and CCMII) showed very high heritability for the mutant and ratio traits, whereas wild-type siblings had much lower repeatability (**Table S14**).

GWA was performed using the HapMap3 SNP data set (Bukowski *et al.* 2018). Although 343 inbred maize lines were crossed to *Oy1*-*N1989*/*oy1*:B73, only 305 inbred lines were genotyped as part of HapMap3 and were subsequently used for GWAS. The variation in mutant plant’s CCMI, CCMII, and their ratios identified a single locus that passed a multiple test correction (see Methods) on chromosome 10 at the site of the *oy1* gene. Just like the detection of the *vey1* QTL in the B73 x Mo17 RIL, no other loci modifying these traits were identified in the GWA analysis. No statistically significant peak was detected for the wild-type CCM traits. The Manhattan plot showing the negative log_10_ of the p-values from GWA tests for all the SNPs for MT_CCMII trait is graphed in **Figure 4a**. A closer view of the SNPs within the region encoding the *vey1* locus on chromosome 10 are plotted in **Figure 4c**. A summary of the GWAS results for mutant CCM and ratio traits is presented in **Table S15**. The top association for the mutant CCM traits was a SNP at position 9161643 on chromosome 10 located just 3’ of the *oy1* locus (a gene on the reverse strand). This SNP displays high allelic frequency in our population (f=0.49). Thus, it appears that the *oy1* locus can be responsible for the suppression of the *Oy1-N1989* mutant allele in the diverse panel of maize inbred lines analyzed in this experiment. Analysis of the LD between S10_9161643 and the other variants in this region identified no other variants with r^2^ greater than 0.5 despite the relatively strong associations between many SNPs and the CCM traits (**Figure 4e**). LD was substantially higher for SNPs encoded towards the telomere from *oy1* than towards the centromere, with a strong discontinuity of LD at the 3’-end of *oy1*. Given the relatively low p-values calculated for multiple SNPs in the area, this raises the possibility that multiple alleles could contribute to the suppression of the *Oy1*-*N1989* phenotype. To test for multiple genetic effects at this locus, the genotypes at SNP S10_9161643 were used as a covariate, and the genetic associations were recalculated. If SNPs segregate independently of S10_9161643 and contribute to *Oy1-N1989* suppression, the p-values of association test statistics for such variants should decrease in this analysis (become more significant). On the contrary, those SNPs that have relatively low p-values due to linkage with S10_9161643 should become less significant in the covariate model. When these analyses were done, low-frequency variants at the 5’ end of the *oy1* locus were identified as the most significant SNPs and passed a chromosome-wide multiple test correction (**Figure 4b and 4d**). This result suggests that there are multiple alleles capable of modifying the *Oy1-N1989* mutant phenotype in the MDL panel. The top SNP on chromosome 10 in the covariate model of MLM was detected at position 9179932 and the allele associated with *Oy1-N1989* suppression is a relatively rare variant (f=0.08). It remains formally possible that the SNP S10_9179932 is not causative and merely in LD with a causative polymorphism, and the second locus is fortuitously present in recombinant haplotypes. Analysis of LD of S10_9179932 with other SNPs in ∼500 kb window detected multiple SNPs of low allelic frequency that were in high LD (r^2^ ∼0.85) towards the 5’ end of *oy1* (**Figure 4f**). Consistent with the strong discontinuity of LD at 3’ end of *oy1* with S10_9161643, we observed discontinuity of LD with S10_9179932 at 5’ end of the *oy1* locus. The LD analyses suggest that S10_9161643 and S10_9179932 are not in LD with each other and can act independently. These two SNPs account for ∼27 percent of the mutant CCMII variation in the MDL x *Oy1-N1989/oy1*:B73 F_1_ population (**Table S15**).

**Figure 4.**
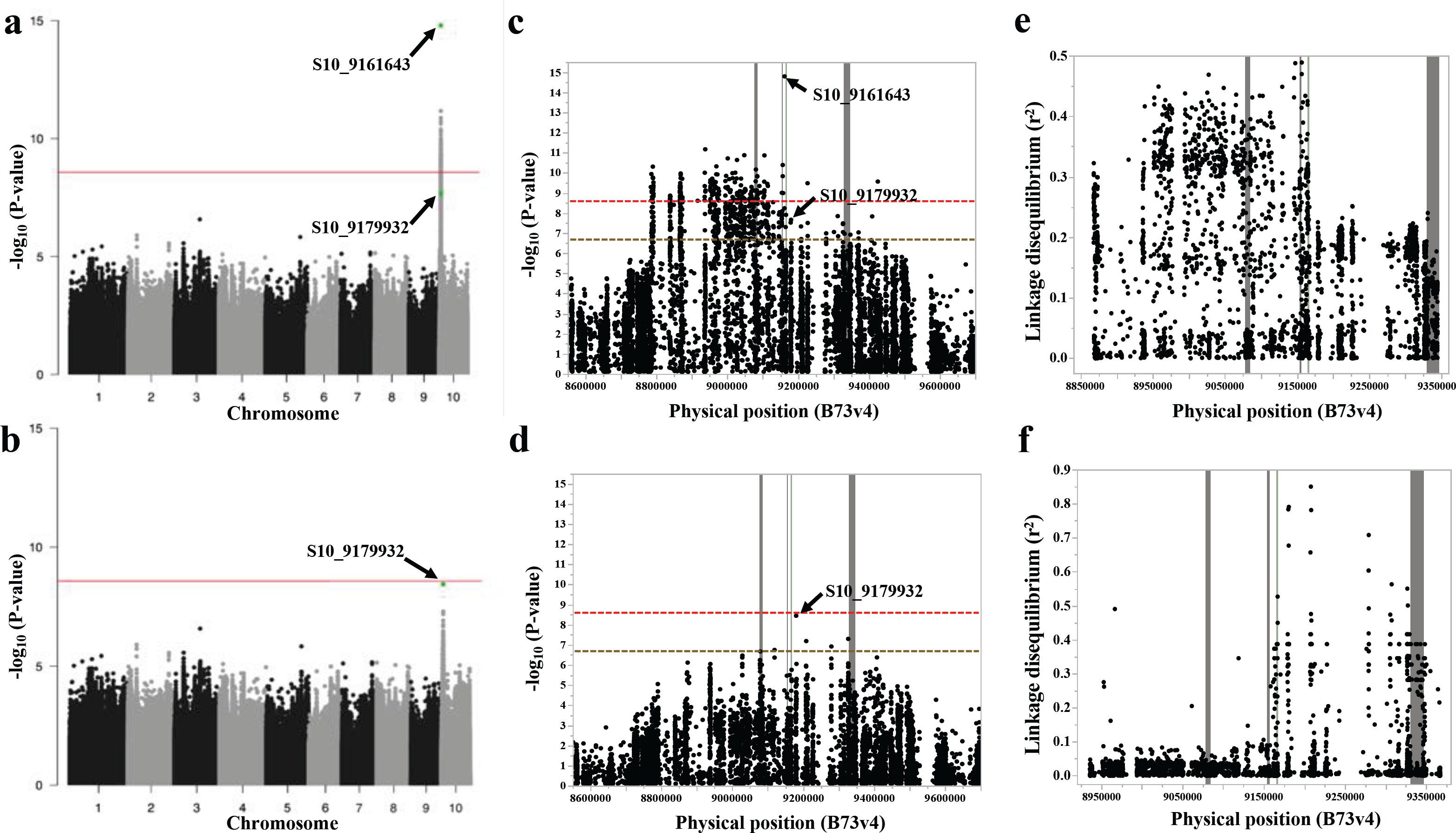
The Manhattan plots of SNPs associations with MT_CCMII trait in MDL x *Oy1-N1989/oy1*:B73 F_1_ populations. The genome-wide association of (a) MT_CCMII, (b) MT_CCMII using S10_9161643 as a covariate. The close-up view of the region on chromosome 10 for (c) MT_CCMII result shown in panel a, (d) MT_CCMII results shown in panel b. Arrows in panels a-d identify the data point corresponding to SNPs S10_9161643 and S10_9179932. The horizontal red and hashed red lines in panels a-d indicates the genomewide Bonferroni cut-off at p<0.05 and hashed golden line in panels c-d is the chromosome-wide FDR-adjusted threshold at p<0.05. The linkage disequilibrium of all SNPs in a ~450 kb region around oy1 with SNPs (e) S10_9161643, and (f) S10_9179932. Vertical lines in panels c-f from left to right represent the genomic position of *ftcl1, ereb28, oy1* (green), and *gfa2* loci.

Given that the MLM model using S10_9161643 as a covariate detected S10_9179932 as the most significant association, we tested the phenotypic outcome of the four possible haplotypes at these two SNPs in the MDL F_1_ population. We observed that the four haplotypes at these two SNPs varied only for mutant CCM traits, with haplotypes AG and CA being the most favorable (highest CCM mean) and least favorable (lowest CCM mean), respectively (**Table S16**). Alleles at these two SNPs affected CCMI and CCMII in the mutant plants, consistent with additive inheritance for two polymorphisms. The additive impacts of the SNPs make mechanistic predictions about the suppression of *Oy1-N1989*. This additivity is consistent with independent alleles acting in cis at *oy1* to modify the *Oy1*-*N1989* mutant phenotype. Consistent with the strong enhancement caused by crossing Mo17 to *Oy1-N1989/oy1*:B73, the Mo17 *oy1* locus encodes the most severe, and relatively rare, CA allele combination, and B73 encodes the most suppressing AG allele combination (**Table S16**). Thus, the line-cross mapping was performed with inbred lines that carry the most phenotypically extreme allele combinations of these two SNPs in the vicinity of the *oy1* locus.

### *Oy1-N1989* is a semi-dominant chlorophyll mutant and enhanced by reduced function at *oy1*

If alleles of *oy1* encode the suppression of *Oy1-N1989*, then the phenotype of heterozygous *Oy1-N1989* should be strongly responsive to other mutant alleles at *oy1*. Maize seedlings that are heterozygotes between *Oy1-N1989* and hypomorphic *oy1* alleles are more severe than isogenic *Oy1-N1989*/*oy1* siblings (Sawers *et al.* 2006). To confirm that the reduced *oy1* function could determine the differential sensitivity to *Oy1-N1989*, we crossed dominant and recessive mutant alleles of *oy1* to each other. The recessive weak hypomorphic allele *oy1-yg* was obtained from the Maize COOP in the unknown genetic background. The homozygous *oy1*-*yg* plants were crossed as a pollen-parent with both B73 and Mo17 to develop F_1_ material that would segregate the mutation. The F_1_ plants were then crossed to *Oy1-N1989/oy1:*B73 as well as backcrossed to the *oy1-yg* homozygotes in the original mixed background. These crosses allowed us to recover plants that had *Oy1-N1989* in combination with the wild-type *oy1*^B73^, wild-type *oy1*^Mo17^, and mutant *oy1-yg* alleles. Chlorophyll contents were determined using CCM at 21 and 40 days after sowing in the field. The *Oy1-N1989* allele was substantially enhanced when combined with the *oy1-yg* allele, demonstrating that reduced function of the *oy1* allele in Mo17 could be the genetic basis of *vey1* QTL (**Figures 5a and 5b**). A summary of these data is presented in **Table S17**. This result is similar to the one described by Sawers *et al.* 2006, where a reduction in chlorophyll content was observed when *Oy1*-*N1989* mutant allele was combined with a recessive allele of *oy1* (*chlI*-*MTM1*). We also noticed a similar drop in chlorophyll accumulation in the *oy1-yg* homozygotes as oppose to *oy1-yg* heterozygotes with wild-type *oy1* allele from both B73 and Mo17 in the BC_1_F_1_ progenies (**Figure 5c and Table S17**). However, we did not observe any significant difference in the *oy1-yg* heterozygotes with B73 and Mo17 wild-type *oy1* allele. Selfed progeny from *Oy1-N1989* heterozygotes segregated for yellow-seedling lethal *Oy1-N1989* homozygotes with no detectable chlorophyll by either CCM or spectrophotometer quantification (**Table S6**). Therefore, consistent with previous work (Hansson *et al*. 2002; Sawers *et al*. 2006), the *Oy1-N1989* is a dominant-negative neomorphic mutant allele with no evident MgChl activity under the tested conditions. Based on these genetic data, any QTL resulting in decreased expression of *oy1* or an increased proportion of mutant to wild-type gene product in the *Oy1*-*N1989*/*oy1* heterozygotes can increase the severity of the mutant phenotype.

**Figure 5.**
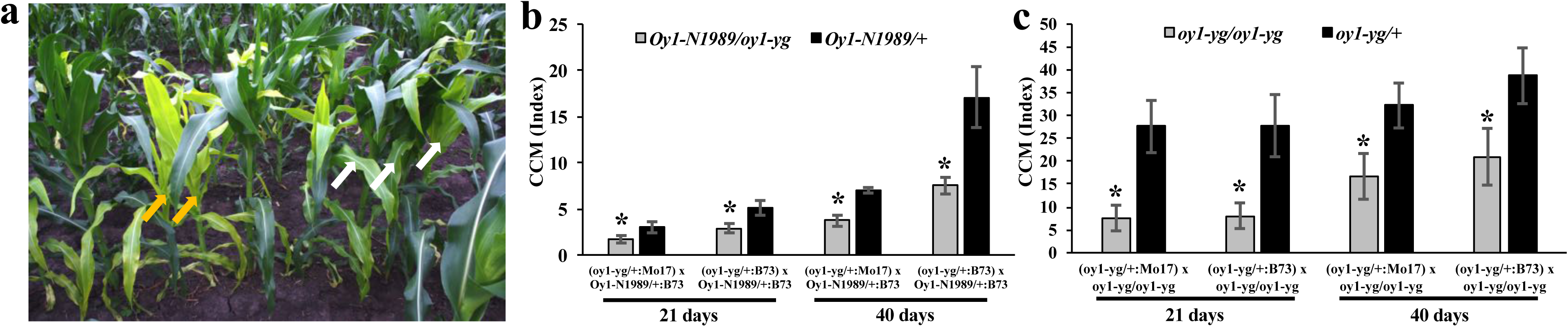
The single locus test of *oy1* showing the interaction between wild-type alleles of *oy1* from B73 and Mo17 with semi-dominant and recessive mutant alleles *Oy1-N1989* and *oy1-yg*, respectively. (a) Mutant (two severity groups) and wild-type individuals segregating in a cross (B73 x *oy1-yg/oy1-yg*) x *Oy1-N1989/oy1*:B73. White-fill arrows indicate *Oy1-N1989/oy1* plants (pale-green and suppressed), whereas yellow-fill arrows indicate *Oy1-N1989/oy1-yg* (yellow-green and severe) plants. The CCM measurements of testcrosses at 21 and 40 days after planting in the (b) mutant siblings (*Oy1-N1989/oy1-yg and Oy1-N1989/* +) of (Mo17 x *oy1-yg/oy1-yg*) x *Oy1-N1989/*+:B73, and (B73 x *oy1-yg/oy1-yg*) x *Oy1-N1989/*+:B73 crosses, (c) mutant (*oy1-yg/oy1-yg*) and wild-type (*oy1-yg/*+) siblings of (Mo17 x *oy1-yg/oy1-yg*) x *oy1-yg*/ *oy1-yg* and (B73 x *oy1-yg/oy1-yg*) x *oy1-yg/ oy1-yg* crosses. Asterisks in panel b-c indicate the significant difference of mean between the genotypes in a given cross at p<0.01 determined using student’s t-test. Check supplemental information for details.

### No coding sequence difference in OY1 accounts for *vey1* inheritance

Our reliance on SNP variation leaves us open to the problem that linked, but the unknown non-SNP variation can be responsible for *vey1*. Given that reduced *oy1* activity enhanced the phenotype of *Oy1-N1989*, we sequenced the *oy1* locus from Mo17 and B73 to determine if coding sequence differences could encode the *vey1* modifier. The only non-synonymous changes that distinguish these two alleles is at the site of the previously reported in-frame 6 bp insertion (Sawers *et al.* 2006), which adds alanine (A) and threonine (T) amino acid residues to the OY1 protein. PCR amplification of *oy1* locus in 18 maize inbred lines, as well as the *Oy1-N1989* allele, was performed. Sequencing of the amplification products confirmed the absence of the 6 bp insertion in *Oy1*-*N1989* allele reported by Sawers *et al.* 2006. In addition, multiple inbred lines including B73, CML103, and CML322 also carried this 6 bp in-frame deletion. A polymorphism within the 6 bp insertion was also found that resulted in an alternative in-frame insertion encoding an alanine and serine (S) codon in Mo17 and five other inbred lines. Thus, three alleles at this site were found to be a common variant in OY1 gene product. These allelic states of *oy1* did not explain the phenotypic severity of CCM trait value in the F_1_ mutant siblings (**Figure 6**). The allelic state of *oy1* at this polymorphic site in 18 maize inbred lines and the average CCM trait values in the wild-type and mutant siblings of their respective F_1_ progenies with *Oy1-N1989/oy1*:B73 are summarized in **Table S18.** Five inbred lines, including Mo17, resulted in dramatic enhancement of the CCMI and ratio of CCMI phenotypes of F_1_ plants crossed to *Oy1-N1989/oy1*:B73. These enhanced genotypes encoded all three possible alleles at *oy1*. In addition, the suppressing inbred lines also encoded all three possible alleles. Besides this 6 bp indel, three inbred lines had few more variants in OY1 protein. An enhancing inbred line CML322 had two missense mutations that lead to amino acid change at position 321 (D->E), and 374 (S->I). A suppressing inbred line NC358 had one amino acid change at position 336 (D->G) and the enhancing inbred Tzi8 had a 15 bp in-frame deletion leading to the removal of five amino acids (VMGPE) in the third exon of the coding sequence. Even considering the additional alleles at *oy1* found in few maize inbred lines, these results suggest that the only coding sequence polymorphism at *oy1* between B73 and Mo17 could not be genetic basis of *vey1*. This result leaves the two additive top SNPs in cis with *oy1* as the most likely cause of cis-acting regulatory variation.

**Figure 6.**
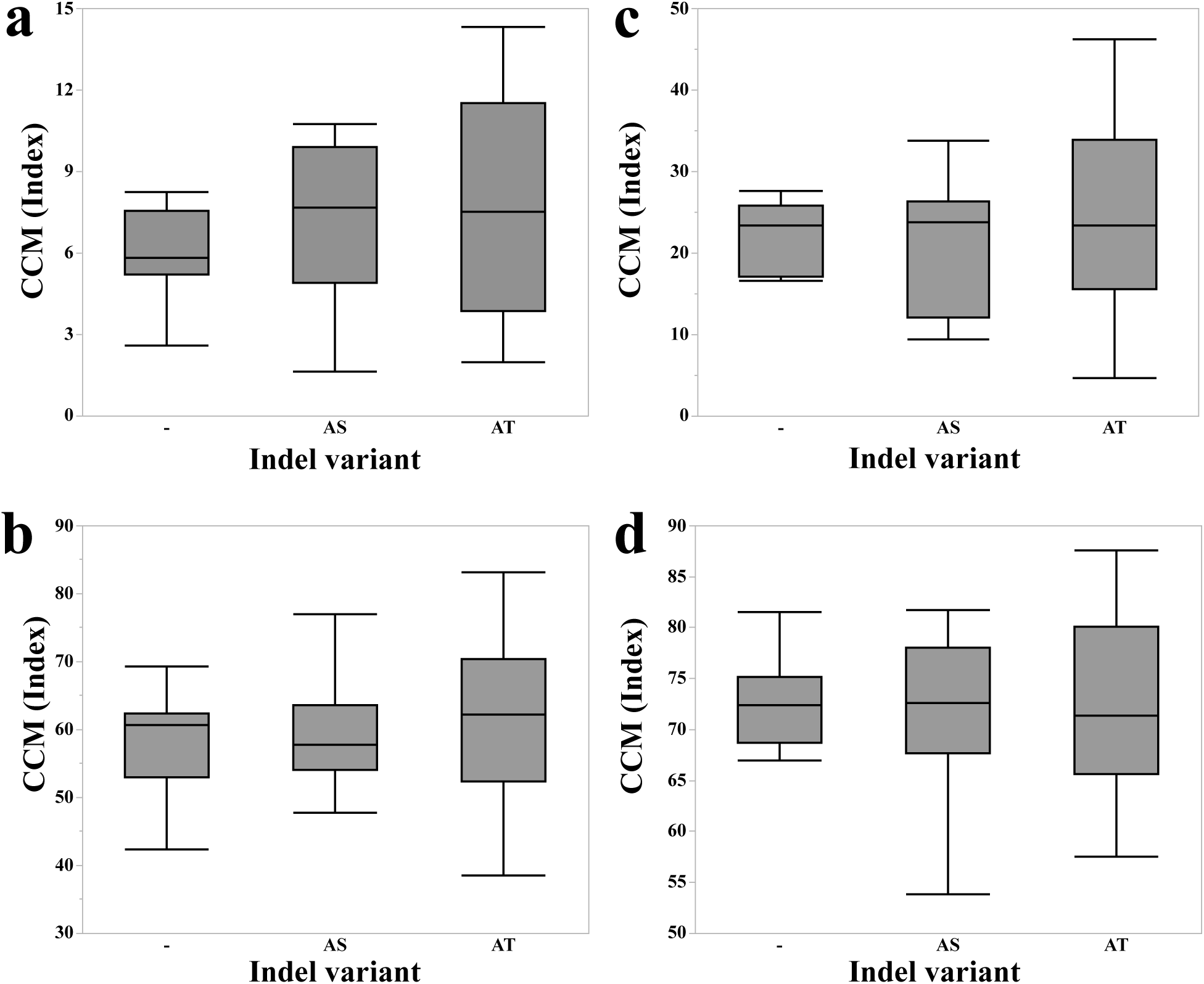
The distributions of CCM trait measurements in the F_1_ progenies of a sub-set of maize inbred lines crossed with *Oy1-N1989/oy1*:B73 at three allelic variants in the *oy1* coding sequence identified in respective inbred lines. The phenotypic distribution of (a) MT_CCMI, (b) WT_CCMI, (c) MT_CCMII, and (d) WT_CCMII. Symbols “-”, “AS”, and “AT” on the X-axis denote deletion of 6 base pairs (bp), insertion of amino acid residues Alanine-Serine (AS), and Alanine-Threonine (AT), respectively. Three inbred lines including B73 carried “-” allele, six inbred lines carried “AS” insertion, nine inbred lines carried “AT” insertion. No statistically significant difference was found among the three distributions in all panels using ANOVA. Check the supplemental information for more details.

### Expression level polymorphism at *oy1* co-segregates with suppression of *Oy1-N1989*

Measurements of mRNA accumulation from *oy1* in the IBM was available in a previously published study (Li *et al*. 2013, 2018). The normalized transcript abundance (expressed as RPKM) of OY1 from the shoot apex of 14 days old maize seedlings from IBM were obtained from MaizeGDB (Sen *et al.* 2010). Out of 105 IBM lines that were assessed for expression level, 74 were among those tested for chlorophyll accumulation in *Oy1*-*N1989* F_1_ hybrid populations. Using the genetic marker isu085b that is linked to *oy1* locus, we determined that a cis-acting eQTL controlled the accumulation of OY1 transcripts in IBM shoot apex **(Figure 7)**. A summary of these data is presented in **Table S19**. This cis-acting eQTL conditioned greater expression of the B73 allele and explained 19 percent of the variation (p < 0.0001) in OY1 transcript abundance in the IBM. Given the enhancement of *Oy1-N1989* by the *oy1-yg* allele, a lower expression of the wild-type *oy1* allele from Mo17 is expected to enhance the phenotype of *Oy1-N1989* (**Figure 5**). In addition, OY1 RPKM values obtained from the shoot apex of IBM were able to predict the CCM trait values in the mutant but not the wild-type siblings in IBM F_1_ population with *Oy1-N1989/oy1*:B73 (**Figure 7**). This result suggests that inbred lines with increased MgChl subunit I transcripts available for protein production and MgChl complex assembly could overcome chlorophyll accumulation defects caused by the *Oy1*-*N1989/oy1* genotype in IBM F_1_ hybrids. Consistent with this, full linear regression model that included both the isu085b marker (cis-eQTL) genotypes and the residual variation in RPKM at OY1 did a better job in predicting CCMI and CCMII in the IBM mutant F_1_ hybrids than the isu085b marker by itself. If the cis-eQTL at *oy1*, which results in differential accumulation of OY1 transcripts in the IBM inbred lines can affect allele-specific expression in the F_1_ hybrids, it could explain the better performance of the IBM mutant F_1_ hybrids with the B73 allele at *vey1*.

**Figure 7.**
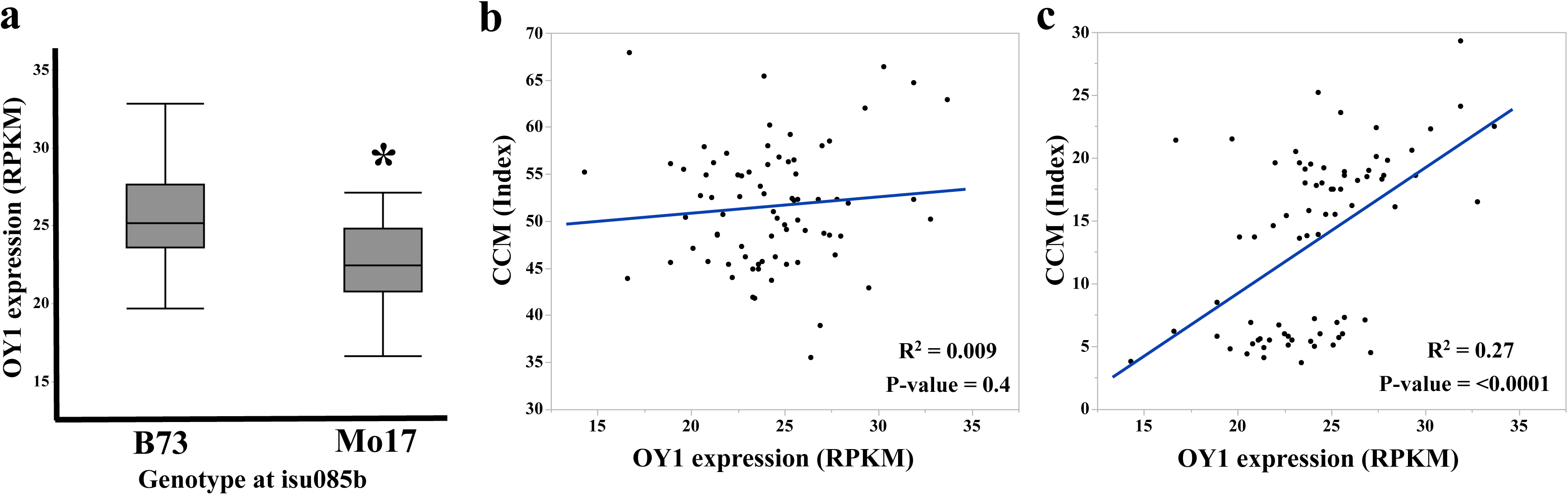
Expression of OY1 in the shoot apices of 14 days old IBM seedlings co-segregates with *vey1*. (a) Genotypic distribution of OY1 RPKM (X-axis) at the marker isu085b (linked to *vey1*). An asterisk indicates the significant difference in the mean between two groups using Student’s t-test at p<0.01. The linear regression of OY1 expression in IBM on CCMII in the (b) wild-type and (c) mutant siblings derived from IBM x *Oy1-N1989/oy1*:B73 crosses.

A previous study of allele-specific expression in the F_1_ hybrid maize seedlings identified expression bias at *oy1* towards B73 in the hybrid combinations of B73 inbred line with PH207 and Mo17 but not Oh43 (Waters *et al.* 2017). We used two SNP positions, SNP_252 and SNP_317, to explore the allele-specific expression of OY1 in our materials. SNP_252 is the causative polymorphism for the *Oy1-N1989* missense allele while SNP_317 is polymorphic between B73 and Mo17, but monomorphic between *Oy1*-*N1989* and B73. As the original allele of the *Oy1-N1989* mutation was isolated from a *r1 c1* colorless synthetic stock of mixed parentage (G. Neuffer, personal communication), this raises the possibility that the same cis-acting regulatory variation that lowered expression of OY1 from PH207 and Mo17 when combined with the B73 allele might also be present in the *oy1* allele that was the progenitor of *Oy1-N1989*. We tested this possibility by using the SNPs that distinguish B73, Mo17, and the *Oy1-N1989* alleles to measure allele-specific expression in each of the hybrids. Consistent with the previous data (Waters *et al.* 2017), we observed biased expression at *oy1* towards the B73 allele in the B73 x Mo17 F_1_ wild-type hybrids (**Table 1**). Extended data from this experiment is provided in **Table S20**. In the B73 isogenic crosses, transcripts from the *Oy1-N1989* and B73 wild-type alleles accumulated to equal levels in the heterozygotes, indicating that the suppressed phenotype of the mutants in B73 background was not due to a lowered expression of *Oy1-N1989* relative to the wild-type allele. Remarkably, mutant siblings from the reciprocal crosses between *Oy1-N1989/oy1*:B73 and Mo17 resulted in greater expression from the *Oy1-N1989* allele than the wild-type *oy1* allele of Mo17. Allele-specific bias at *oy1* was significantly higher towards the *Oy1*-*N1989* allele in the *Oy1-N1989* mutant heterozygotes in the B73 x Mo17 hybrid background compared to B73 isogenic material. Thus, in mutant hybrids, overexpression of *Oy1-N1989* relative to the wild-type *oy1* allele in Mo17 could account for increased phenotypic severity.

**Table 1.** The allele-specific expression at *oy1* in the top fully-expanded leaf at the V3 developmental stage of B73 x Mo17 F_1_ wild-type and *Oy1*-*N1989*/*oy1* mutant siblings, and inbred *Oy1*-*N1989*/*oy1:B73* mutants.

| Genotype<sup>1</sup> | SNP_252 | SNP_317 | Ratio_SNP252 | Ratio_SNP317 | Average |
| --- | --- | --- | --- | --- | --- |
| *oyl/oyl*:B/M<sup>&</sup> | . | . | . | . | 1.19±0.07 |
| *oyl/oyl*:B/M | C/C | C/T | . | 1.08±0.01<sup>a</sup> | 1.08±0.01<sup>a</sup> |
| *Oyl-N1989/oyl*:M/B | C/T | C/T | 1.12±0.01<sup>a</sup> | 1.10±0.02<sup>a</sup> | 1.11±0.01<sup>a</sup> |
| *Oyl-N1989/oyl*:B/M | C/T | C/T | 1.15±0.03<sup>a</sup> | 1.10±0.01<sup>a</sup> | 1.13±0.02<sup>a</sup> |
| *Oyl-N1989/oyl*:B | C/T | C/C | 1.01±0.02<sup>b</sup> | . | 1.01±0.02<sup>b</sup> |

<sup>1</sup>B73 is denoted as B, Mo17 is denoted as M, B/M denotes B73 x Mo17 cross direction and M/B is vice-versa. The mean ± standard deviation of the ratios of the read count from the reference/alternate allele at SNP\_252, SNP\_317, and the average of the ratios at SNP position 252 and 317. The connecting letter report for each trait indicates the statistical significance calculated using ANOVA with post-hoc analysis using Tukey's HSD with p<0.01.
<sup>&</sup>Data obtained from Waters et al. 2017.

If *vey1* is encoded by an eQTL, then PH207 should encode an enhancing allele and Oh43 should encode a suppressing allele of *vey1.* We tested this genetically by producing F_1_ progenies in crosses of PH207 and Oh43 by *Oy1-N1989/oy1*:B73 pollen. Oh43 was evaluated in our initial screening and also as part of the MDL panel used for GWAS. In both experiments, the F_1_ hybrids between Oh43 and *Oy1-N1989/oy1*:B73 suppressed the mutant phenotype, suggesting that Oh43 is a suppressing inbred line (CCM values in **Tables S4 and S18**). B73 x PH207 F_1_ hybrids were missing from our previous datasets. We crossed PH207 ears with pollen from *Oy1-N1989/oy1*:B73 plants. The F_1_ hybrids from this cross were analyzed in the greenhouse at seedlings stage along with F_1_ hybrids of B73 x *Oy1-N1989/oy1*:B73 and Mo17 x *Oy1-N1989/oy1*:B73 F_1_ progenies as controls. PH207 was an enhancing inbred genotype as mutant heterozygotes in a PH207 x B73 F_1_ genetic background accumulated less chlorophyll than mutants in the B73 isogenic background (**Figure S8 and Table S21**).

We further leveraged the normalized expression data of OY1 in the emerging shoot tissue of the maize diversity lines (Kremling *et al.* 2018) and used the top two additive SNPs (S10_9161643 and S10_9179932) at *vey1* from GWAS to test if these cis-variants of *oy1* affect its expression. When tested, plants carrying alleles A and G at marker S10_9161643 and S10_9179932, respectively, showed highest OY1 abundance in the emerging shoots of diverse maize inbred lines, whereas, plants with alleles C and A at S10_9161643 and S10_9179932, respectively, showed lowest OY1 count (**Table S22**). Alleles that suppress *Oy1-N1989* linked to either SNP were associated with the greater abundance of OY1 transcripts. This observation is consistent with the hypothesis that increased OY1 abundance can overcome the negative effect of the *Oy1-N1989* allele. Consistent with the additive suppression of leaf greenness in *Oy1-N1989* mutants by the alleles at S10_9161643 and S10_9179932 discussed previously (**Table S16**), these alleles were also additive for their impacts on OY1 transcript abundance (**Table S22**). Thus, it is likely that multiple phenotypically affective polymorphisms linked to these top unlinked SNPs underlie cis-acting regulatory variation at *oy1*.

The effect sizes of the gene expression changes observed in the IBM, diverse maize inbred lines, and allele-specific expression in hybrids are quite modest, resulting in ∼10% of differences in *oy1* accumulation. If these changes in wild-type OY1 transcript accumulation are responsible for suppression of *Oy1-N1989*, then the severity of the mutant phenotype (as indicated by CCM) in the MDL F_1_ population should correlate with OY1 expression level. As expected, we observed a statistically significant positive correlation between the mutant derived CCM traits and OY1 counts (**Table S23**). Wild-type CCM in the MDL F_1_ population did not show any significant correlation with OY1 abundance in the emerging shoot tissue of maize inbred lines. These correlations are in agreement with the lack of any QTL at this locus controlling wild-type chlorophyll levels, and the epistatic relationship between *vey1* and *Oy1-N1989*.

## Discussion

The semi-dominant mutant allele *Oy1-N1989* encodes a dominant-negative allele at the *oy1* locus, which compromises MgChl enzyme activity (Sawers *et al.* 2006). In a heterozygous condition, the strength of the negative effect of this allele on the MgChl enzyme complex depends on the wild-type *oy1* allele. B73 and Mo17 show differential suppression response in mutant heterozygotes resulting in suppressed and severe mutant phenotype, respectively. Thus, the *Oy1*-*N1989* mutant allele can sensitize maize plants to variation in MgChl and expose a phenotypic consequence for genetic variants that are otherwise invisible. Similar methodology has been adopted previously in maize to gain the genetic understanding of various traits (Chintamanani *et al.* 2010; Olukolu *et al.* 2013, 2014; Buescher *et al.* 2014). Employing a mutant allele as a reporter to screen for effects of natural variants is a simple and efficient technique to detect standing variation in a specific biological process. Although *vey1* polymorphisms are phenotypically consequential in the presence of *Oy1-N1989* allele, no QTL was detected in the absence of *Oy1-N1989* allele. These cryptic genetic variants, or contingent QTL, result from epistasis of *Oy1-N1989* and permits the discovery and re-classification of DNA sequence variants that might otherwise be hypothesized to be neutral or non-functional in plant adaptation. Thus, genetic screens based upon semi-dominant mutant alleles as reporters offer a cost-effective and robust approach to map QTL(s) for metabolic pathways of interest by leveraging the publicly available genetic resources such as bi-parental mapping populations and maize diversity lines.

Alleles with modest fitness consequences may not be visible to researchers working with population sizes even in the thousands, such as in GWAS. By contrast, evolutionarily-relevant segregating variation may have minimal phenotypic effects. One of the possible uses of MAGIC is in boosting the relative contribution of natural variants to phenotypic variance in the trait of interest. This can both uncover cryptic variation affecting a biochemical pathway of interest and assist in improving the mapping resolution of a detected QTL. Not all of the cryptic variation observed by these mutant-contingent QTL approaches need be fitness-affecting, and interpreting mutant-conditioned phenotypes as non-neutral variation would be a mistake. It is, of course, possible that neutral variants may result in increased severity of a mutant phenotype due to changes not physiologically relevant for all alleles of that reporter gene present in a species. Nevertheless, it can identify new pathway member and inform us about the allelic variation in the species and pathway topology via gene discovery.

In the current study, use of bi-parental mapping populations derived from the same inbred lines but developed using different intercross and inbreeding schemes provided the opportunity to compare the effect of additional rounds of random interbreeding in the development of mapping population on the genetic resolution. The comparative fine mapping of *vey1* in the Syn10 population, that employed ten rounds of random mating, provided far better localization of the *vey1* QTL than the IBM populations that was derived from four rounds of random mating. This observation demonstrates the benefits of increased recombination during random intermating of early generations in QTL localization. Based on these results, as future RILs are generated, we recommend increased intermating in the early generations followed by DH induction rather than relying on further recombination during the self-pollination cycles of RIL development.

As expected, GWAS provided a fine-scale genetic resolution and corroborated the mapping of *vey1.* The distance between the best marker and the *oy1* gene was substantially less in the GWA experiments. GWAS identified two SNPs, one in the 5’ and another in the 3’ intergenic DNA, proximate to *oy1* that represent candidate quantitative trait nucleotides (QTN). The architecture of this region included four haplotypes with every combination of alleles at these two SNPs. The signal detection by multiple unlinked SNPs in GWAS may indicate a complex set of phenotypically affective alleles at the *oy1* locus. Alternatively, it could very well be an artifact of missing the causative polymorphism within our genetic data resulting in strong associations with markers that are tightly linked to the causative variation but unlinked or in repulsion to each other. For example, indel variation is not captured by the approaches used in the GWA analysis. Consistent with two QTN, rather than fortuitous linkage of tag SNPs with a single causative polymorphism, alleles at the two SNPs additively influenced chlorophyll contents in the mutant siblings. Future work to identify the nucleotide changes responsible for the differences between B73, Mo17, and other inbred lines that encode *vey1* will be required to test these possibilities definitively. Together, this study illustrates the complementary nature of line-cross QTL mapping and GWAS to explore the genetics of the trait under investigation.

We used *Oy1-N1989* together with a non-destructive, inexpensive, and rapid phenotyping method to measure leaf chlorophyll. Rapid and robust phenotyping is critical and contributed to the strong correlation between the absolute trait measurement and the estimated phenotypes like CCM. Previous studies have highlighted the importance of benchmarking indirect measurements or proxies for traits of interest. For instance, near infrared reflectance spectroscopy (NIRS) that estimates major and total carotenoids in maize kernels could replace sensitive, accurate, cumbersome, expensive, slow, and destructive measurements by HPLC (Berardo *et al.* 2004). Comparisons of HPLC and NIRS measurements resulted in a correlation of 0.85 for total carotenoids (Berardo *et al.* 2004). A subjective visual score for yellow color in maize kernels was not adequate, yielding a correlation of 0.12 with HPLC measurements of total carotenoids (Harjes *et al.* 2008). Thus, using mutant alleles as reporters to develop methodologies that rely on non-invasive multispectral or hyperspectral data as proxies for specific biochemical compounds will accelerate studies on gene function and allele discovery. This approach can enable genetic studies that are currently deemed unfeasible due to the arduous task of phenotyping large populations for traits only visible in the laboratory.

### How could a 10% change in wild-type OY1 expression affect chlorophyll biosynthesis in the *Oy1*-*N1989*/*oy1* mutant heterozygotes?

Magnesium chelatase (MgChl) is formed by a trimer of dimers of MgChl subunit I interacting with the other subunits of MgChl complex. A previous study found that addition of mutant or wild-type BCHI protein to pre-assembled MgChl complexes resulted in altered reaction rates due to differences in subunit turnover, which occurred on a minutes time-scale (Lundqvist *et al.* 2013). This subunit turnover and reformation of the complex dynamically exchange mutant and wild-type BCHI subunits over time. Therefore, any net increase in the amount of wild-type OY1 in the reaction pool, for instance, due to higher transcription of wild-type *oy1* allele will allow a higher rate of magnesium chelatase activity and result in more chlorophyll biosynthesis. The observation of stronger affinity and greater dissociation rate of BCHI^L111F^ subunits (orthologous to the L176F change encoded by *Oy1-N1989*) for the wild-type subunits (Hansson *et al*. 2002) suggests that exchange of BCHI monomeric units in the magnesium chelatase complex might also differ based on the structure of BCHI protein (Lundqvist *et al.* 2013). In the AAA^+^ protein family, the ATP-binding site is located at the interface of two neighboring subunits in the oligomeric complex (Vale 2000). Since a dimer of functional MgChl subunit I proteins are required for the complex to carry out MgChl activity (Lundqvist *et al.* 2013), approximately 1 in 3 dimers of assembled MgChl subunit I will be active in a 1:1 mixture of wild-type and BCHI^L111F^. Indeed, complexes made from reaction mixtures with equal proportions of wild-type and BCHI^L111F^ subunits resulted in ∼26% of the enzyme activity of an equivalent all-wildtype mix (Lundqvist *et al.* 2013). Therefore, we expect that decreasing expression of the wild-type *oy1* subunit by 10% and creating a 0.9:1.1 mixture of wild-type OY1 and mutant OY1-N1989 protein, respectively, would result in ∼21% activity compared to the activity of all-wildtype mixture. Likewise, increasing the wild-type *oy1* expression by 10% would result in ∼30% activity of MgChl compared to the all-wildtype mixture. This dosage-sensitivity is a general feature of protein complexes (Birchler and Veitia 2012; Veitia 2003; Grossniklaus, Madhusudhan, and Nanjundiah 1996; Birchler and Newton 1981), and the semi-dominant nature of *Oy1-N1989* is dosage sensitive. Taking these observations and proposed models on the dynamics of molecular interaction between the wild-type and mutant BCHI protein subunits (especially BCHI^L111F^) into account, it is formally possible that a small change in the expression of wild-type OY1 can have a significant impact on the magnesium chelatase activity of heterozygous *Oy1*-*N1989*/*oy1* plants. The increase in magnesium chelatase activity due to the even small relative increase in the proportion of wild-type OY1 transcripts over the mutant OY1-N1989 transcripts will read out as a proportional, presumably non-linear, increase in chlorophyll accumulation.

### What is *vey1*?

Variation at the *vey1* locus appears to be the result of allelic diversity linked to the *oy1* locus. The only remaining possibilities are regulatory polymorphisms within the cis-acting control regions of *oy1*. Previous studies have utilized reciprocal test-crosses to loss-of-function alleles in multiple genetic backgrounds to provide single-locus tests of additive QTL alleles in an otherwise identical hybrid background (Dilkes *et al.* 2008). Protein-null alleles of *oy1* isolated directly from the B73 and Mo17 backgrounds could be used to carry out a similar test. The intergenic genomic region in maize is spanned by transposable elements and can be highly divergent between different inbred lines due to large insertions/deletions polymorphisms (SanMiguel and Bennetzen 1998). Consistent with this, inbreds B73 and Mo17 are polymorphic at the region between *oy1* and *gfa2.* These two maize inbred lines share ∼12 kb of sequence interspersed with numerous large insertions and deletions that add ∼139 kb of DNA sequence to Mo17 as compared to B73 (data not shown). However, we did not find any conserved non-coding sequence (CNS) in this region (data not shown). It is conceivable that recombinants at *oy1* between B73 and Mo17 themselves could be identified and used to test the effects of upstream or downstream regulatory sequences. The recombinant haplotypes encoding all four possible alleles at the top two SNPs identified in the GWAS indicate that Mo17 may be a strong enhancer due to more than one causative polymorphism.

In the absence of such recombinants, we can only consider what mechanisms might be consistent with the observed suppression of the chlorophyll biosynthesis defects such as cis-acting effects on OY1 transcript abundance, and allele-specific gene expression. Genotype at *vey1* in wild-type plants accounted for only 19 percent of the variation in OY1 abundance in the shoot apices of the IBM. This QTL accounted for more than 80 percent of the variation in chlorophyll content in mutants. If transcript accumulation is insensitive to the presence of *Oy1-N1989* allele, then the remaining ∼80 percent of the variation in OY1 abundance observed as not *vey1*-dependent in the wild-type IBM might be expected to account for a greater change in phenotype. However, trans-acting effects or environmental effects that would equally affect both alleles at the *oy1* locus would be expected to have a lesser impact on suppression of the deleterious *Oy1-N1989* allele. The effect of cis-acting eQTL on *oy1* expression would eventually result in a greater or lesser proportion of wild-type OY1 protein accumulation. These allele-specific differences will alter the mutant to wild-type subunit stoichiometry which should have a greater impact on chlorophyll content than changes in absolute RPKM, due to the competitive inhibition of MgChl complex activity by *Oy1-N1989* (Sawers *et al.* 2006; Lundqvist *et al.* 2013), as detailed above.

The reciprocal crosses between *Oy1*-*N1989*/*oy1*:B73 and Mo17 did not result in different phenotypes. Thus, *vey1* does not exhibit imprinting. Allele-specific expression at *oy1* locus in the *Oy1*-*N1989* heterozygous mutants demonstrated the existence of functional cis-acting regulatory polymorphism between *Oy1*-*N1989* and both wild-type *oy1* alleles in B73 and Mo17. In addition, *oy1* is affected by a cis-eQTL in the IBM and MDL. Together with our other data, the allele-specific expression at OY1 that was visible when we reanalyzed the data from Waters *et al.* 2017, we propose that *vey1* is encoded by cis-acting regulatory DNA sequence variation (**Figure S9**). Ultimately, transgenic testing of *oy1* cis-acting regulatory polymorphisms identified from assembled maize genomes is required to determine the causative variant(s) encoding *vey1*. Sequence comparisons outside the protein coding sequence of the gene can be quite challenging, especially in maize, as it exhibits limited to poor sequence conservation between different inbred lines (SanMiguel *et al.* 1996). Thus, distinguishing phenotypically affective polymorphisms from the neutral variants is not trivial. As a result, biochemical and *in vitro* studies are the best tool for functional validation of these polymorphisms (Wray *et al.* 2003). Similar experiments have been done to characterize the role of DNA sequence polymorphisms in cis on the expression of downstream genes in case of *flowering locus T* in *Arabidopsis* and *teosinte branched1* in maize (Adrian *et al.* 2010; Studer *et al.* 2011).

### Relevance to research on transcriptional regulation

Since their discovery, the role of regulatory elements in gene function has been recognized as vital to our understanding of biological systems (McClintock 1950, 1956a, 1956b, 1961; Peterson 1953; Jacob and Monod 1961). Gene regulation and gene product dosage are at the forefront of evolutionary theories about sources of novelty and diversification (Ohno 1972; King and Wilson 1975). Transcriptional regulation of a gene can be as important as the protein coding sequence (Wray *et al.* 2003). For instance, complete knock-down of expression of a gene by a regulatory polymorphism will have the same phenotypic consequence as the non-sense mediated decay of a transcript harboring an early stop codon (Willing *et al.* 1996). But we do not have a set of rules, analogous to codon tables, for functional polymorphisms outside the coding sequence of a gene. Detecting expression variation and tying it to phenotypic consequence, especially in the absence of CNS, remains a challenge to this day.

Several eQTL studies ranging from unicellular to multicellular eukaryotic organisms have found abundant cis and trans-acting genomic regions that affect gene expression (Brem *et al*. 2002; West *et al*. 2007; Li *et al*. 2013, 2018). The proportion of cis-acting eQTLs from the total eQTLs detected in various studies including yeast, mouse, rat, humans, eucalyptus, and maize range from 19-92% (Gibson and Weir 2005; Li *et al*. 2013, 2018). Expression polymorphisms are a potential source of variation in some phenotypic traits (Gibson and Weir 2005), and multiple studies detected expression polymorphisms co-segregating with phenotypic variation, including contributions to species domestication (Clark *et al.* 2006; Salvi *et al.* 2007; Schwartz *et al.* 2009; Lemmon *et al.* 2014). But a majority of the cis-eQTLs only exhibit a moderate difference in the gene expression. Detecting such variants and linking them to visible phenotypes may require detailed study using approaches focused on the specific biological process affected by the gene product. We do not yet have a standardized experimental tool for these purposes. As such, we cannot simultaneously identify and characterize the phenotypic impact of most cis-acting eQTLs.

The *vey1* polymorphism detected in the current study co-segregates with a cis-eQTL at *oy1* in IBM (Li *et al*. 2013, 2018) and diverse maize inbred lines (Kremling *et al.* 2018). Using *Oy1-N1989* allele to expose the consequences of these cis-eQTLs allowed chlorophyll approximation using CCM to substitute for expensive, cumbersome, highly sensitive gene expression assays or metabolite measurements. The high heritability of this alternative and direct phenotype, allowed us to scan large populations at a rapid rate to identify the genomic regions underlying the cis-acting regulatory elements and study allelic diversity in the natural population of maize. We propose that the approach we have taken is not likely to be unique to *oy1*. Therefore, we propose that all semi-dominant mutant alleles can be used as reporters to not only detect novel cis-acting gene regulatory elements but also functionally validate previously-detected cis-eQTL(s) from genome-wide eQTL studies.

## Acknowledgements

Authors acknowledge the help of the staff at Purdue ACRE farm for assistance with planting and crop management of all the field experiments described in this study. R.S.K was supported by USAID Feed the Future CGIAR-US Initiative grant “Heat stress resilient maize for South Asia through a public-private partnership” to G.S.J.. This work was supported by NSF grant 1444503 awarded to G.S.J. and B.P.D. We especially thank the Muehlbauer lab for making their data open to the public, and the USDA for the MaizeGDB website, which allowed us to reanalyze the eQTL data.

## Supplemental tables

**Table S1.**
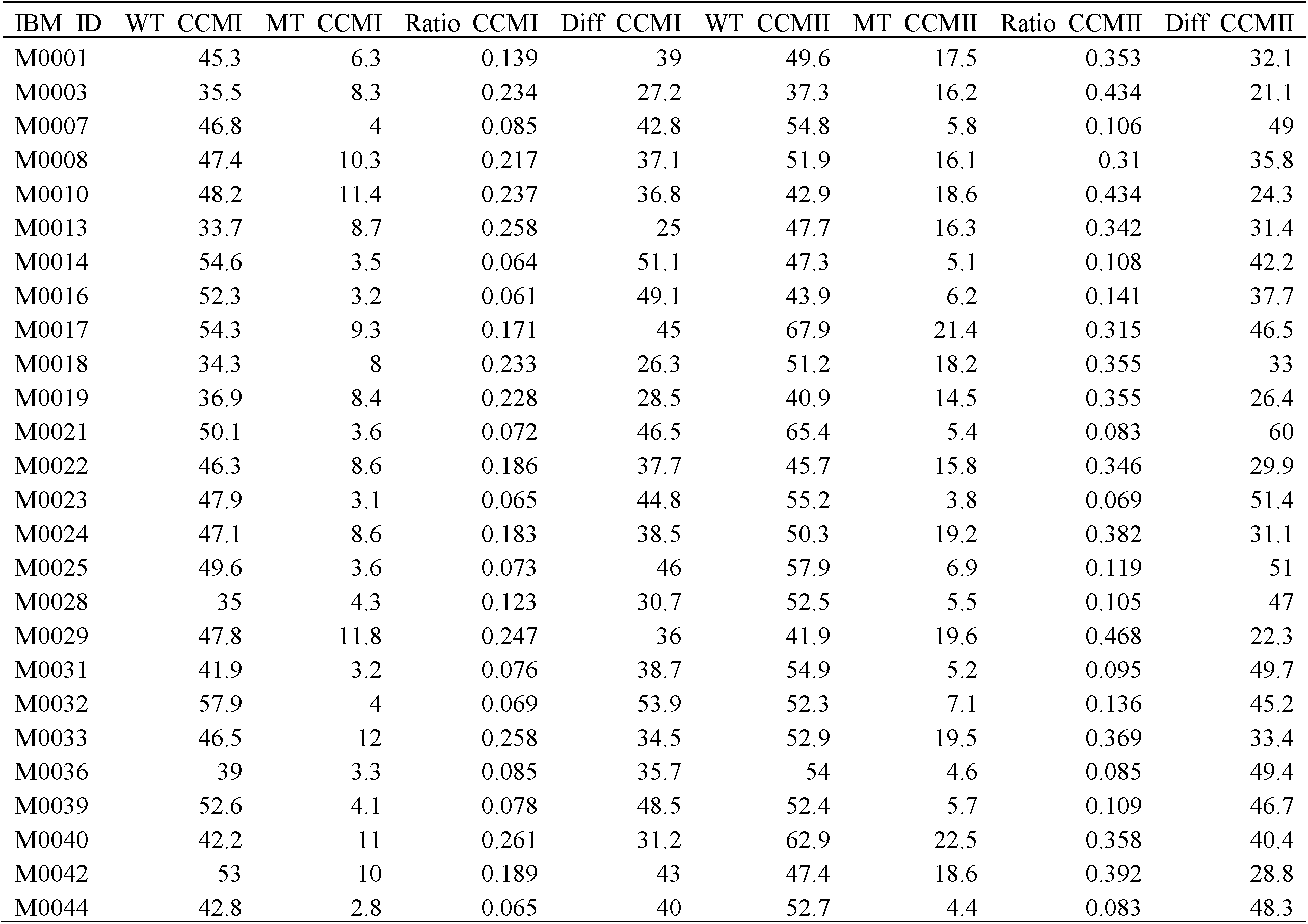
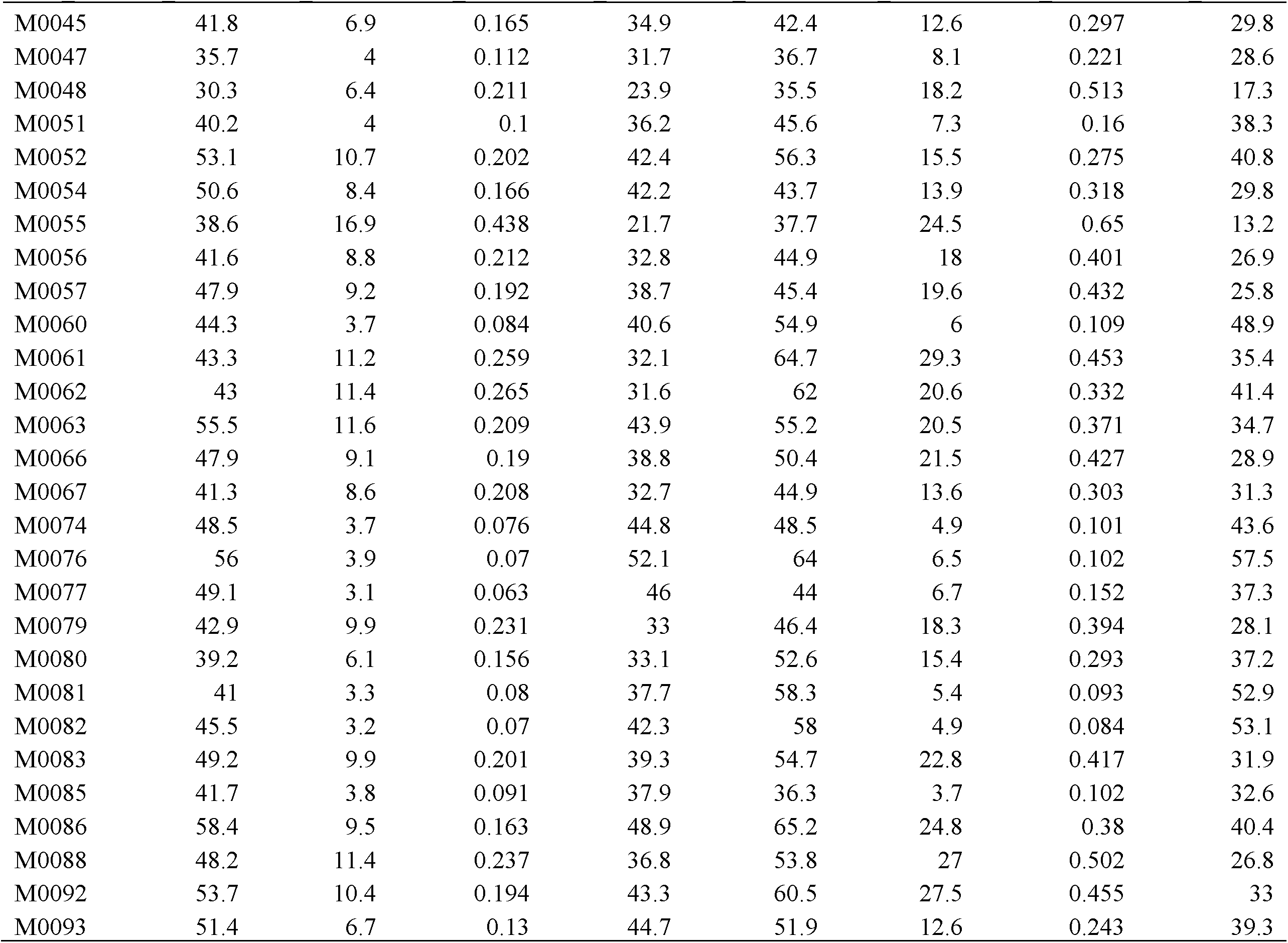
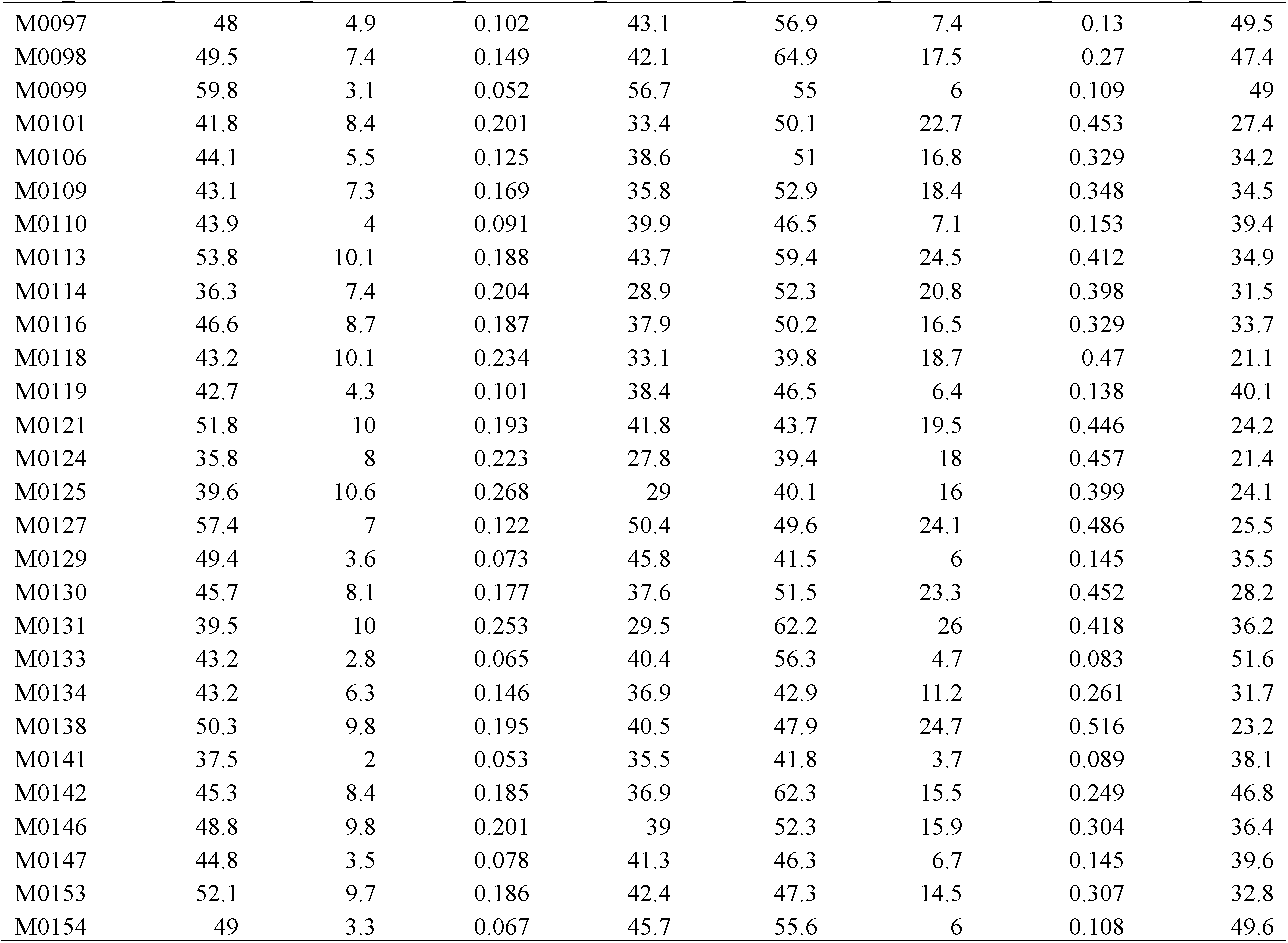
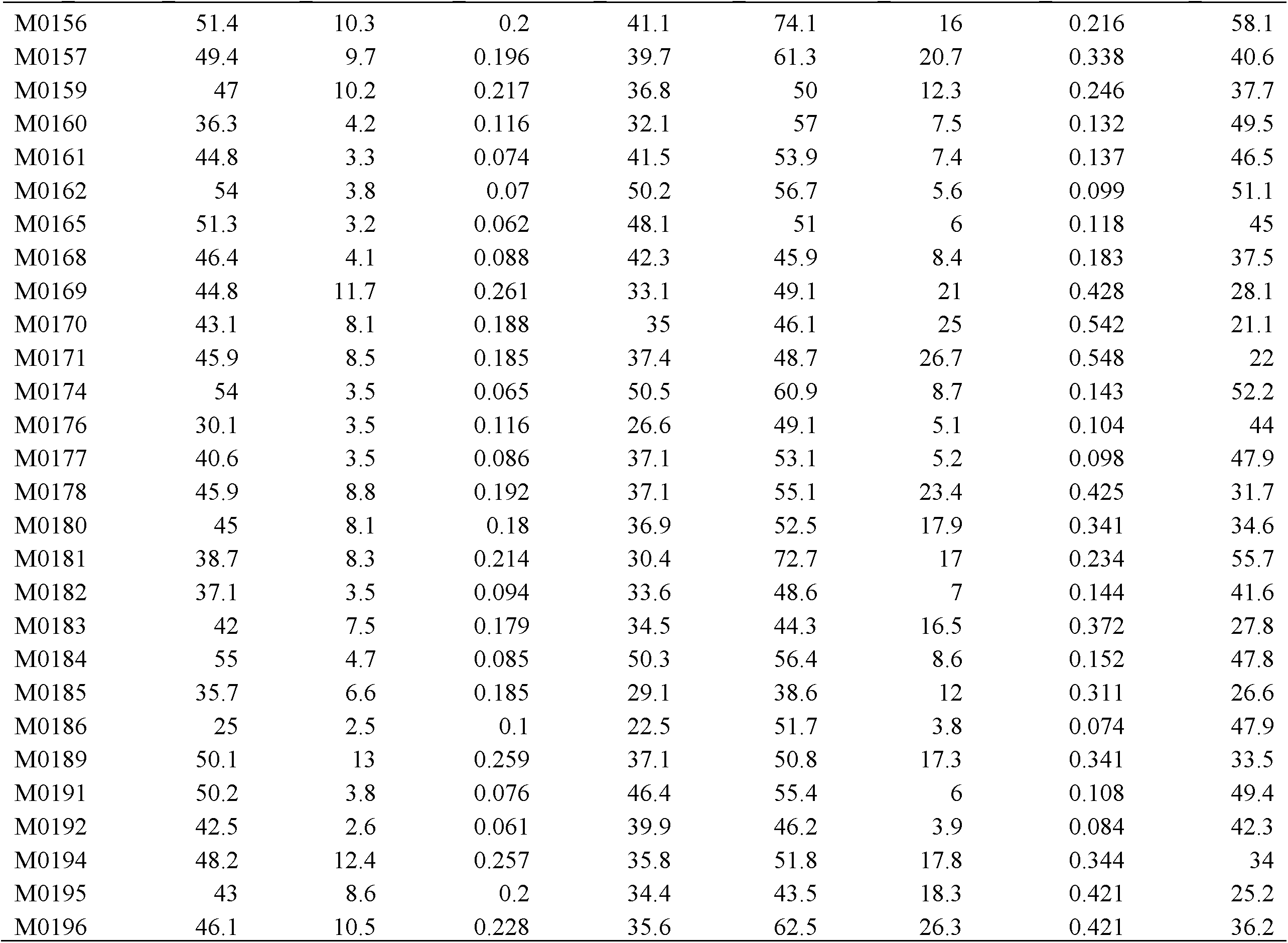
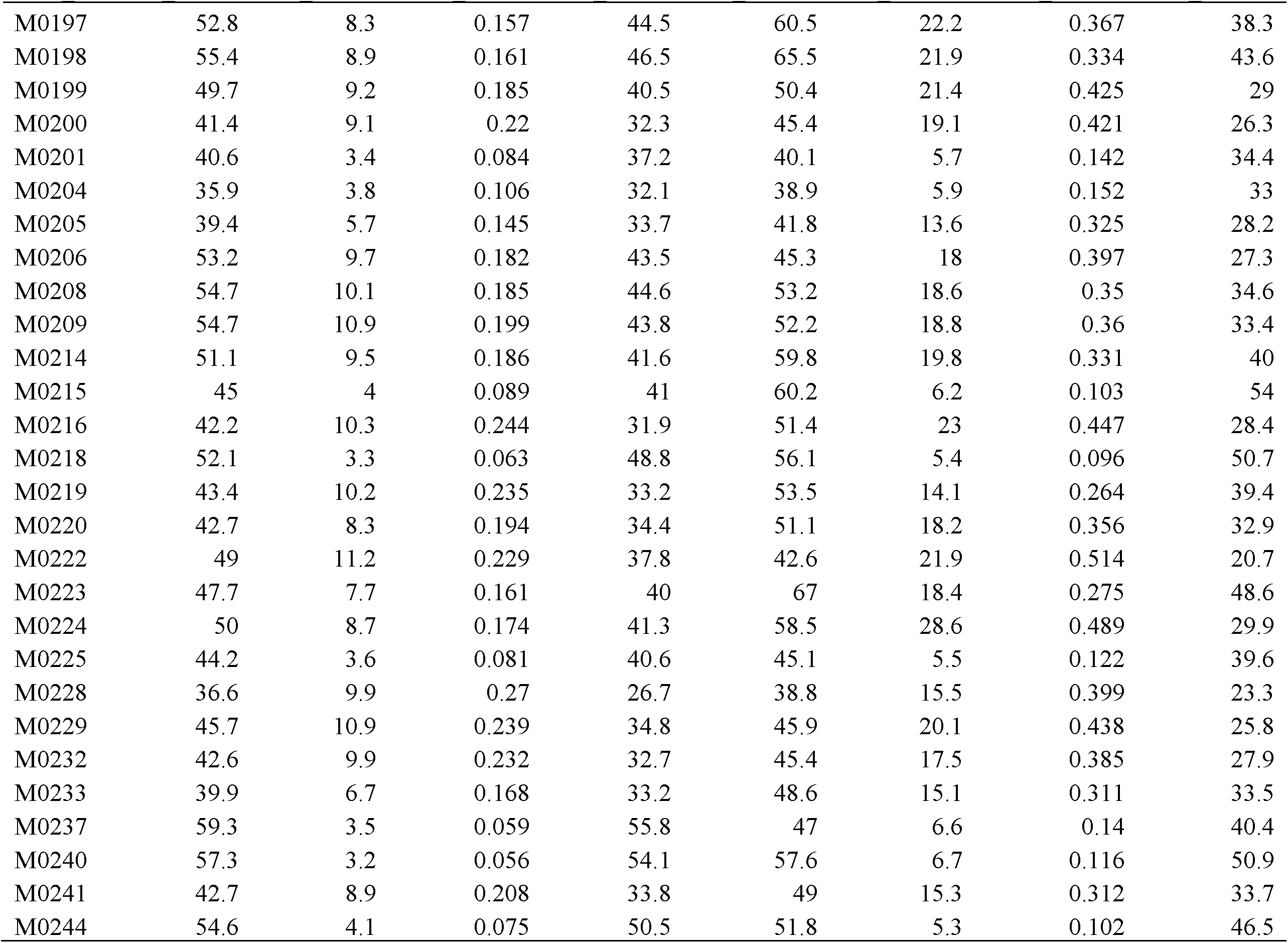
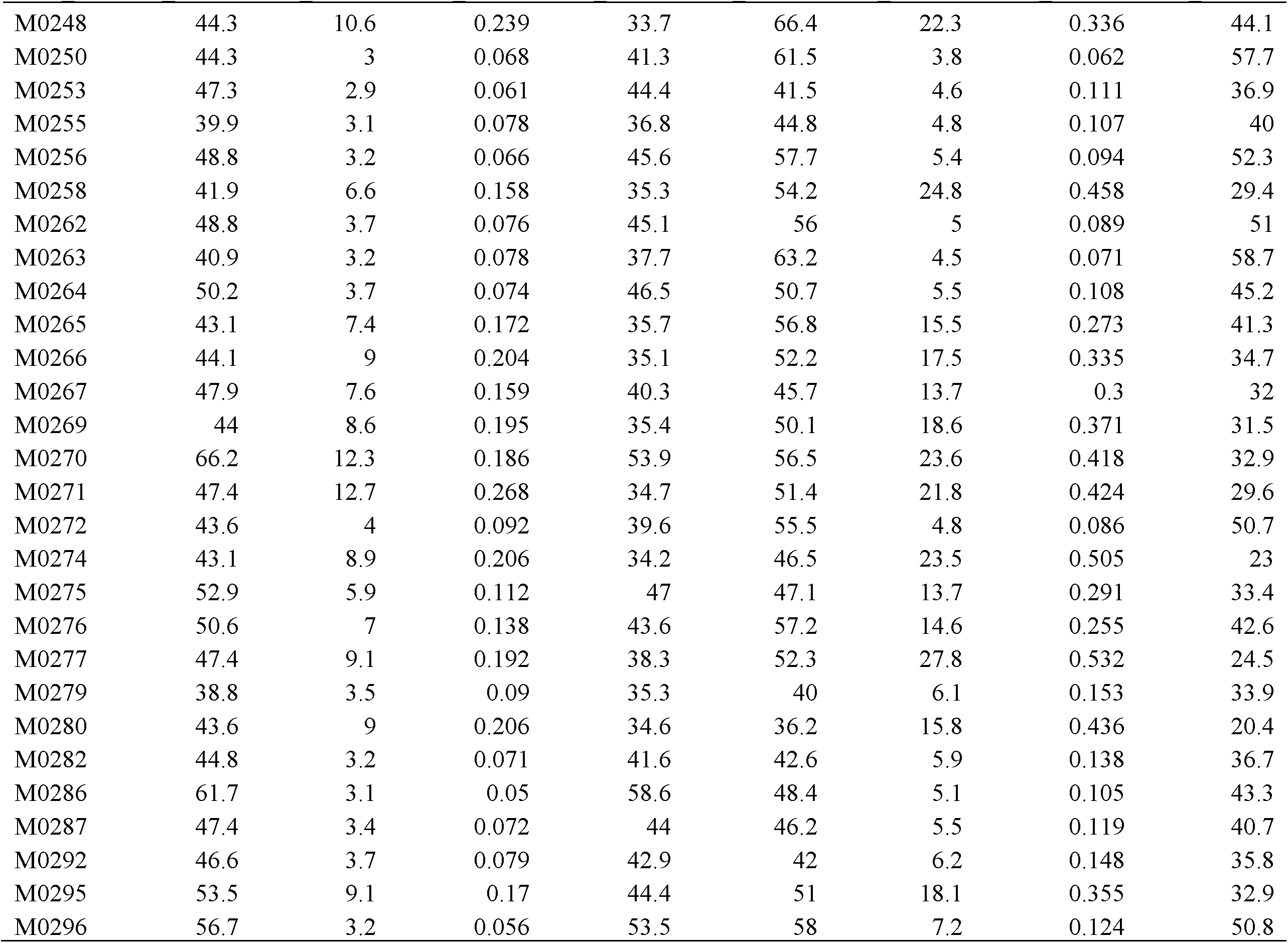
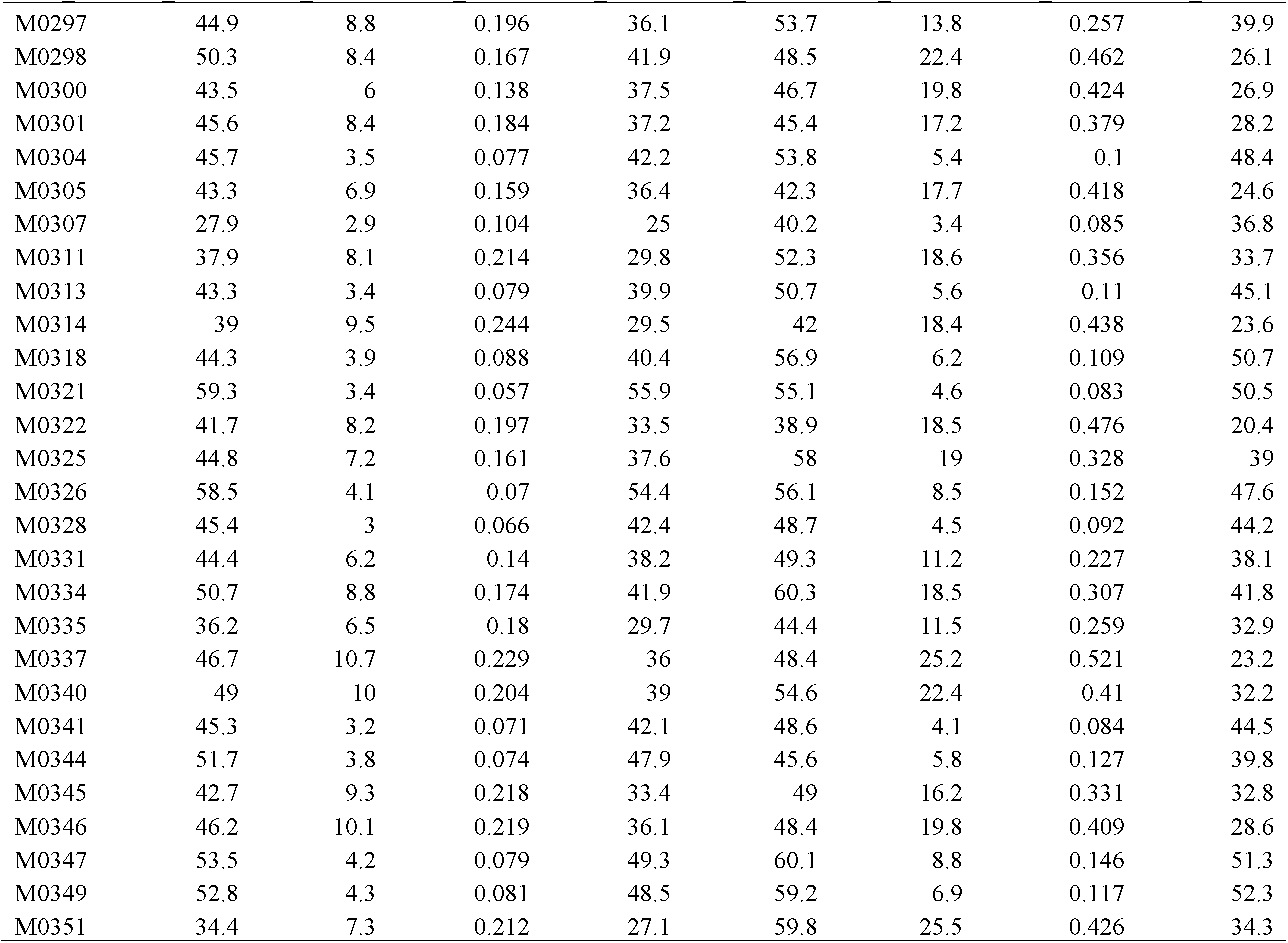
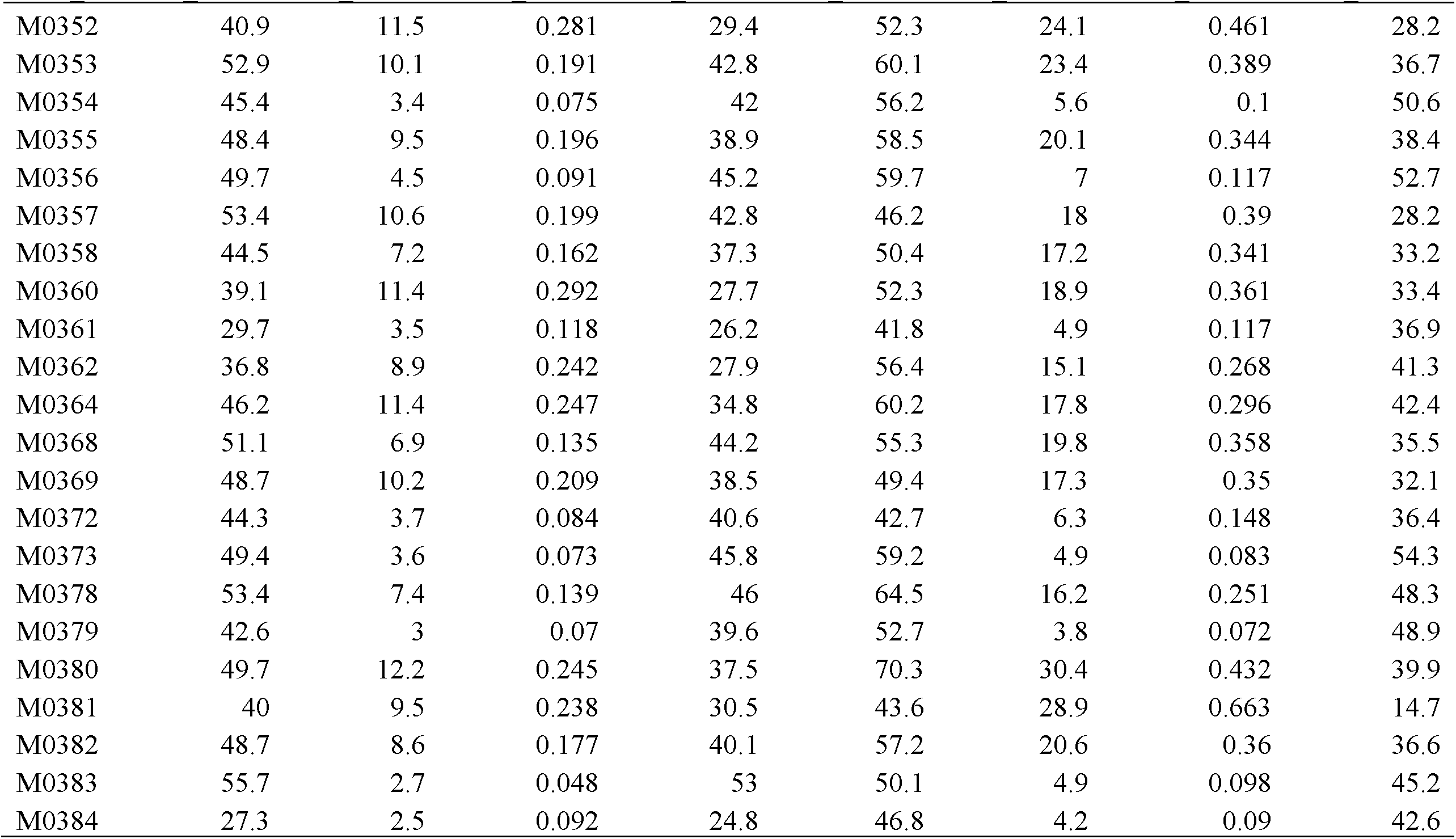
The trait mean values of the CCM traits for the wild-type (WT) and mutant (MT) siblings of the F_1_ hybrids of *Oy1*-*N1989*/*oy1*:B73 (pollen-parent) with respective IBM lines.

**Table S2.**
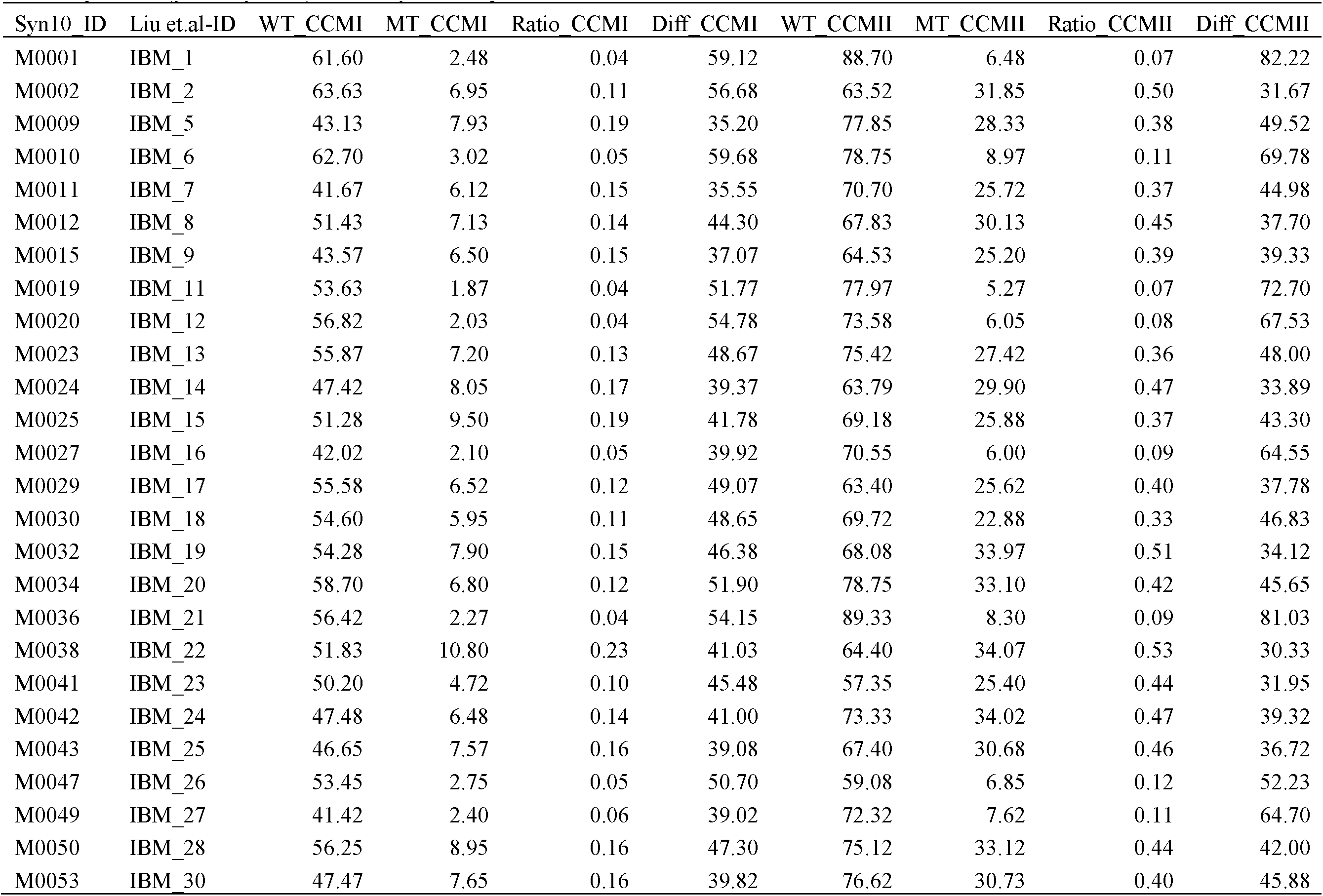
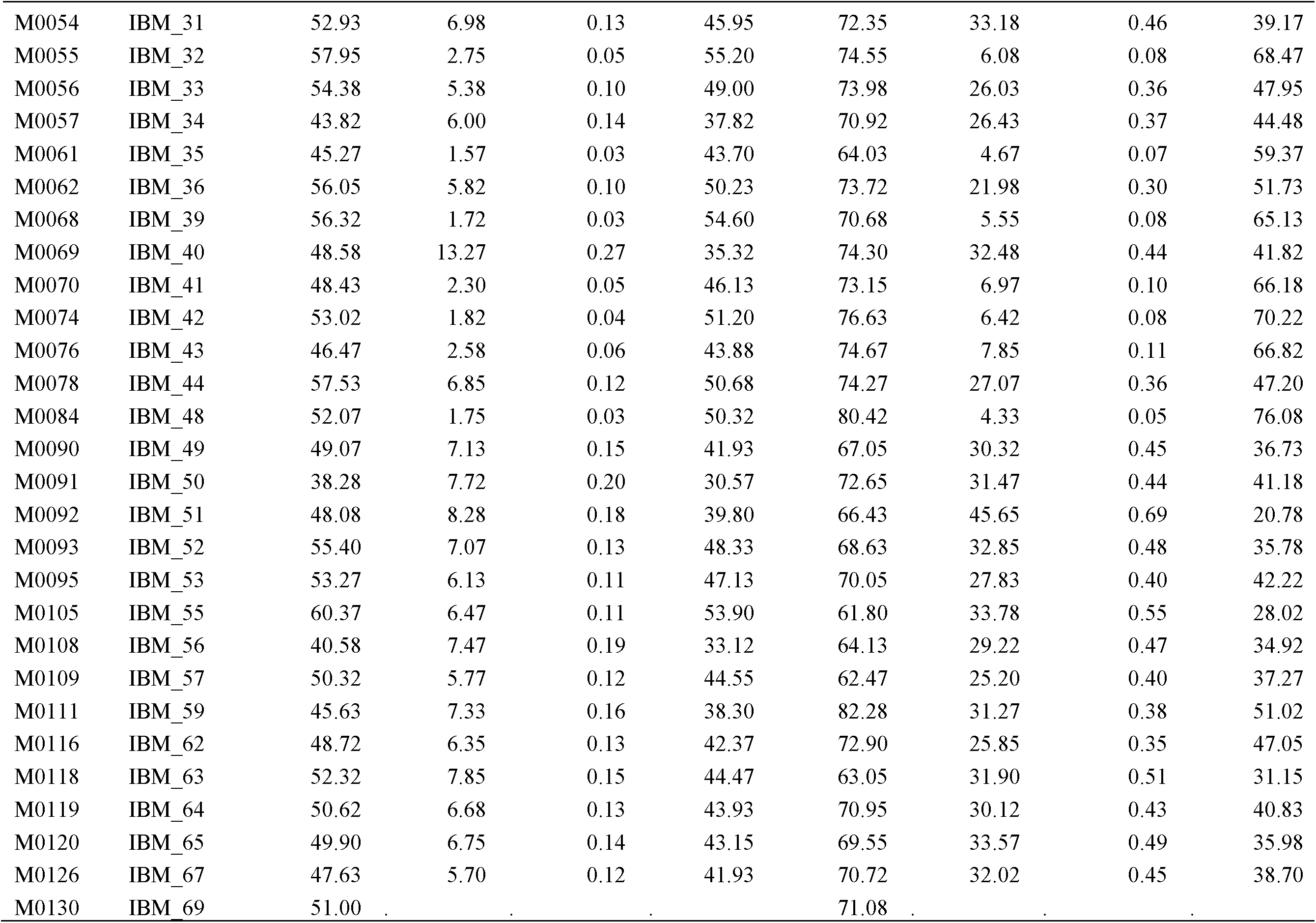
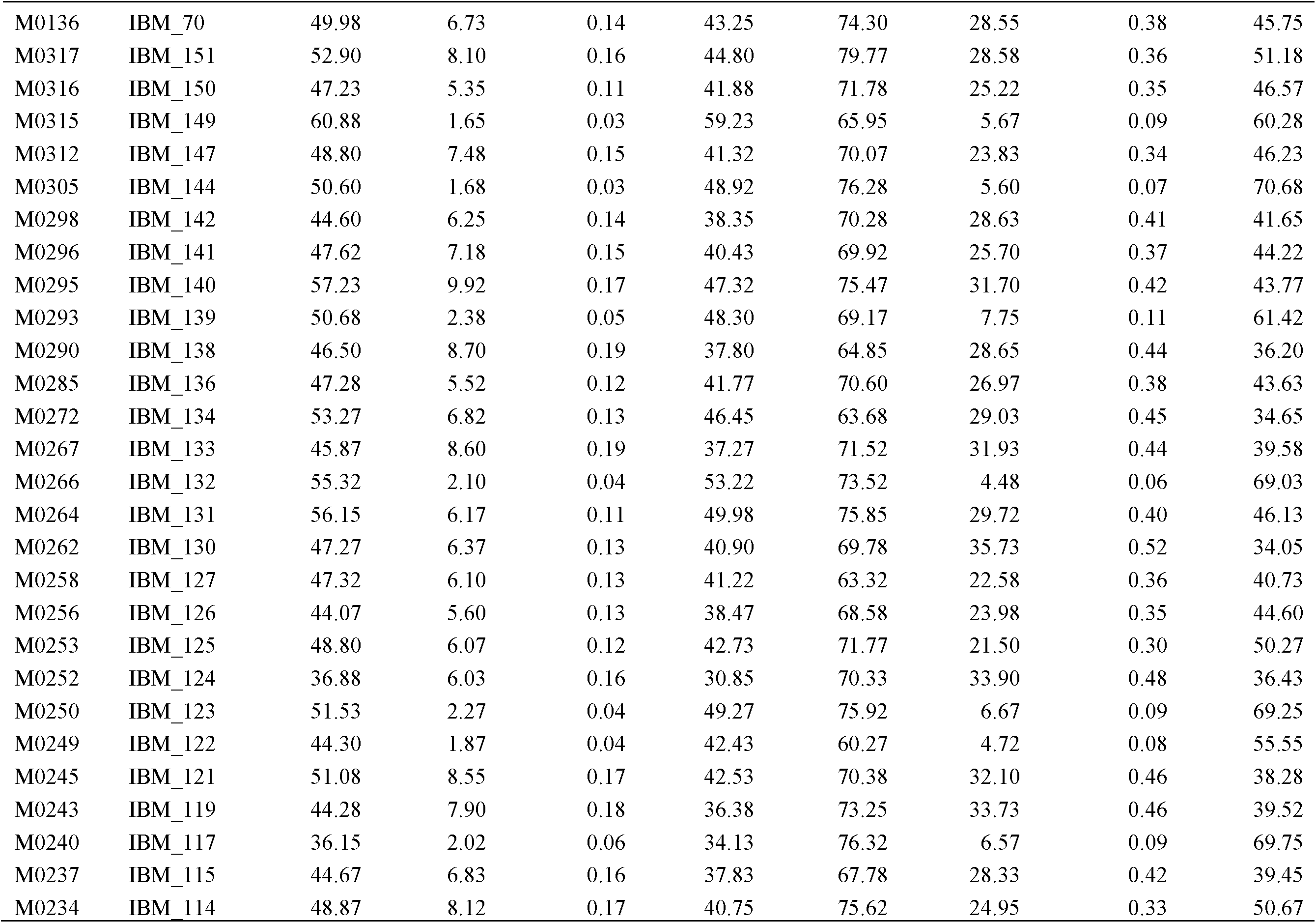
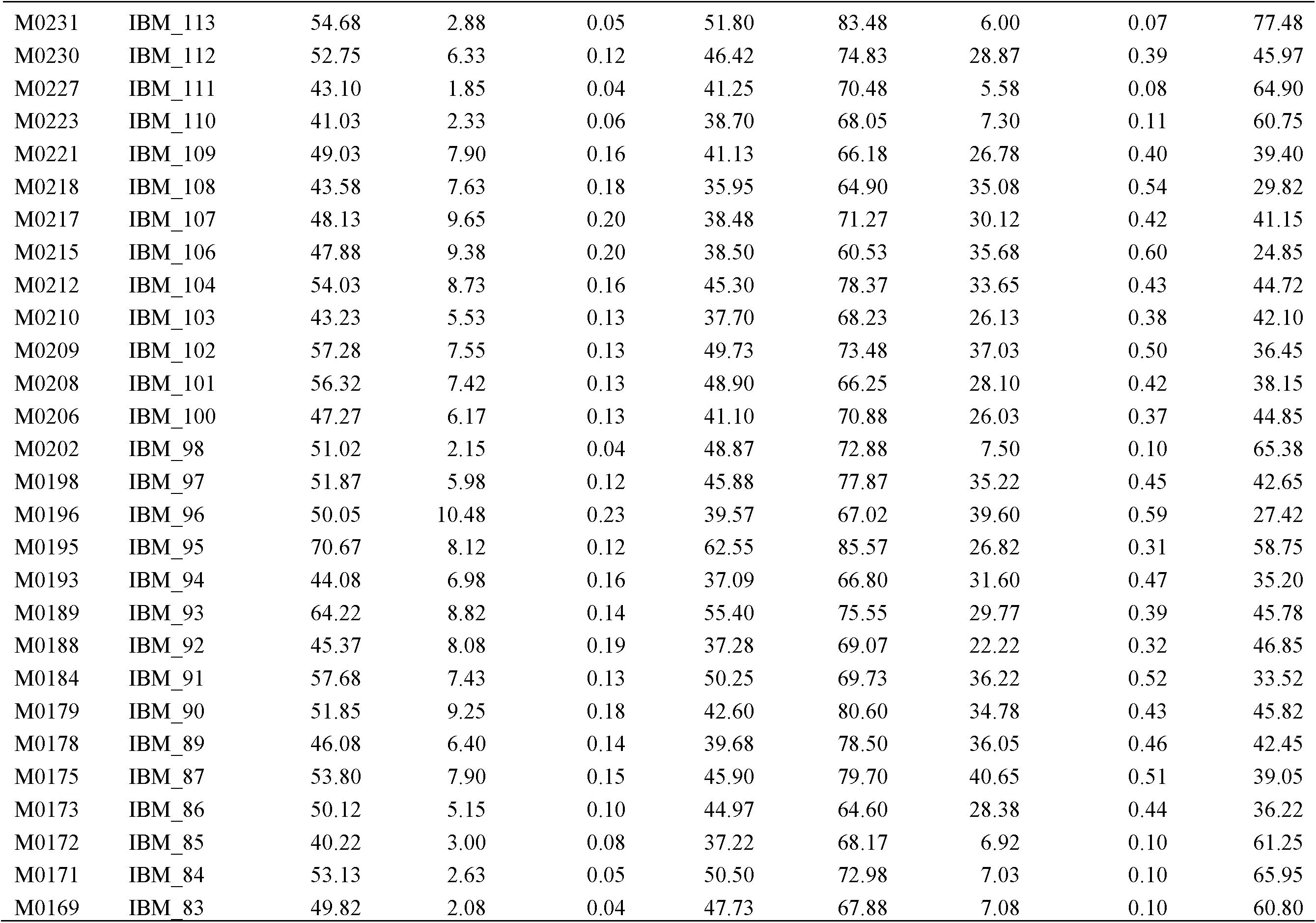
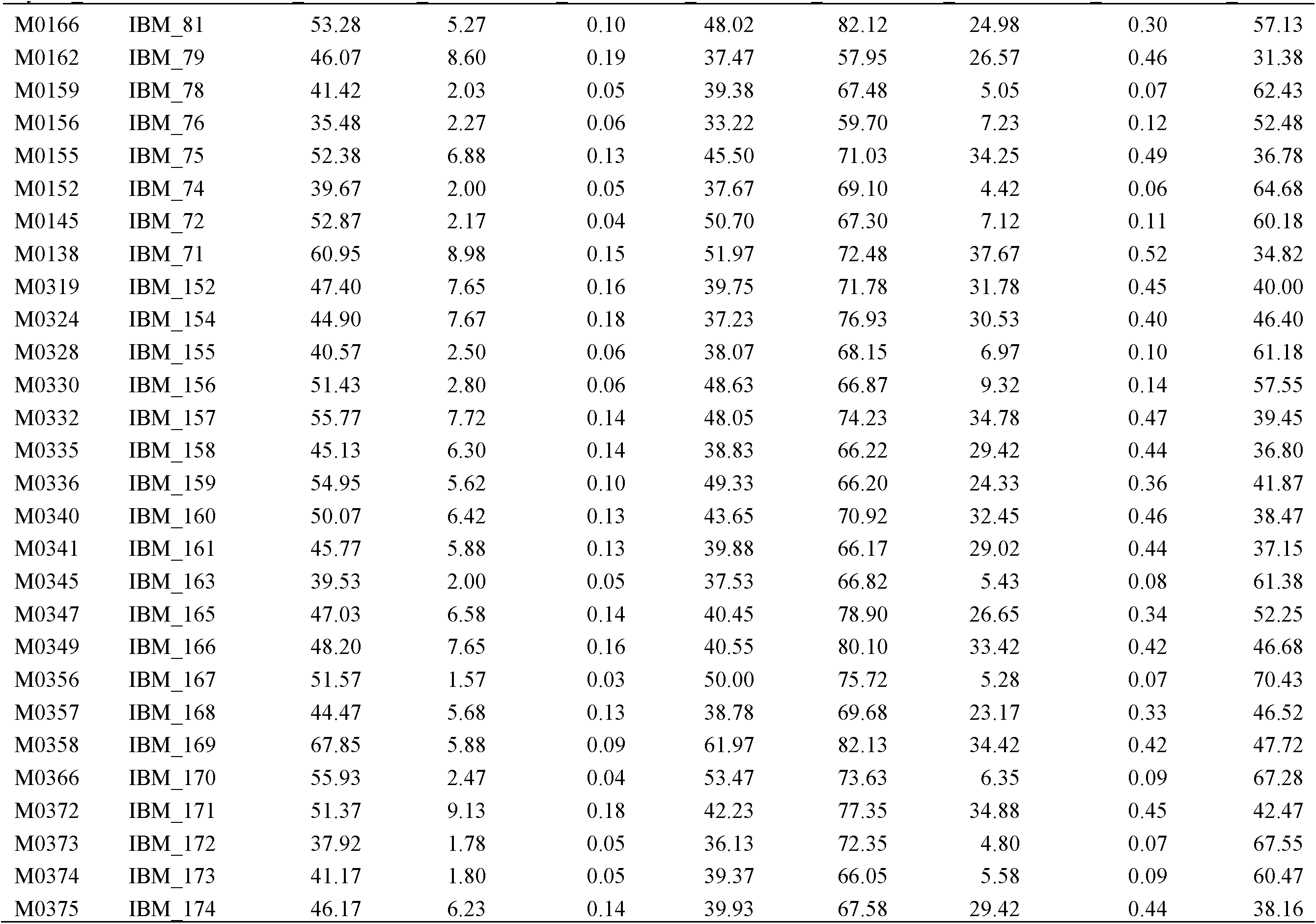
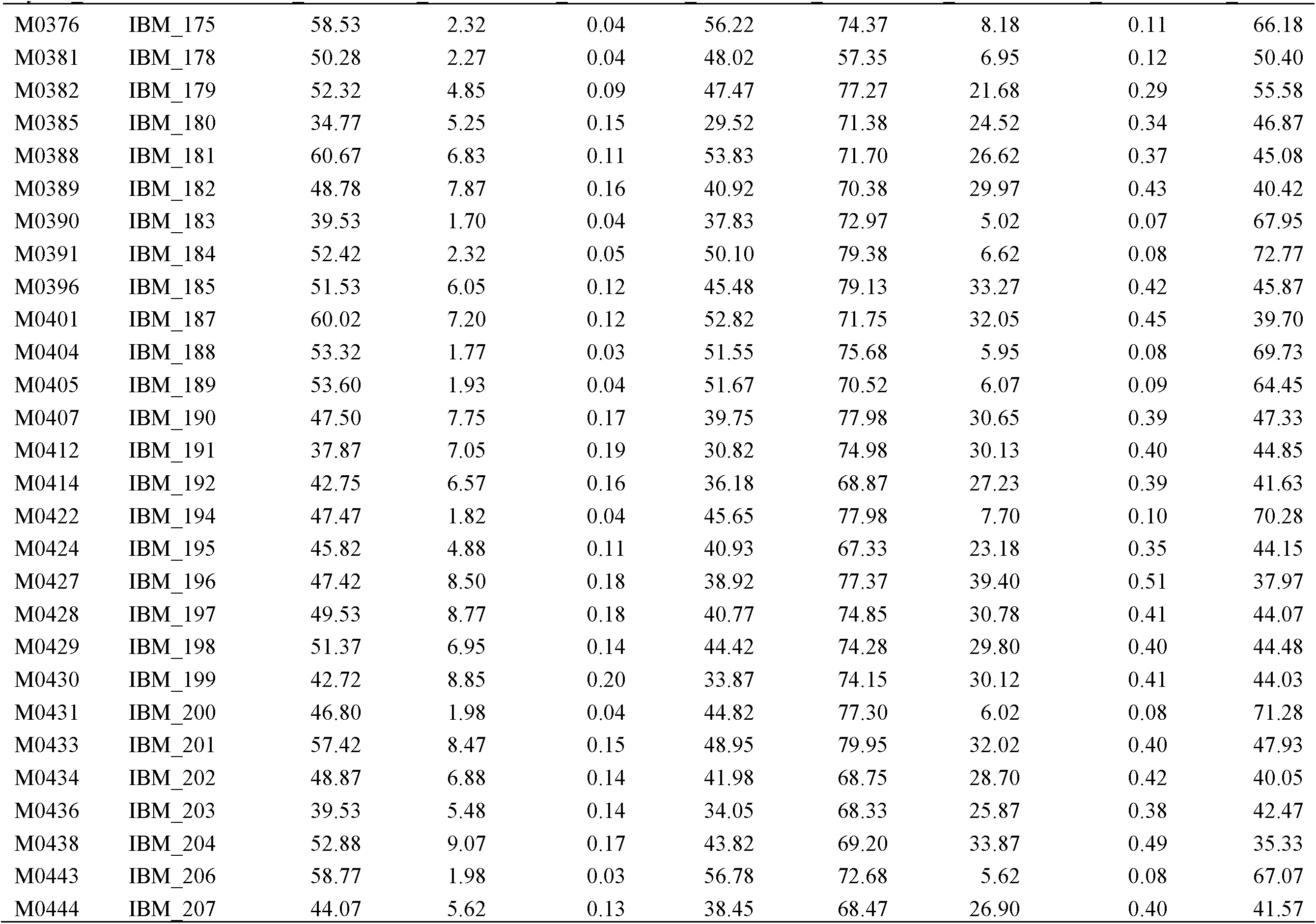
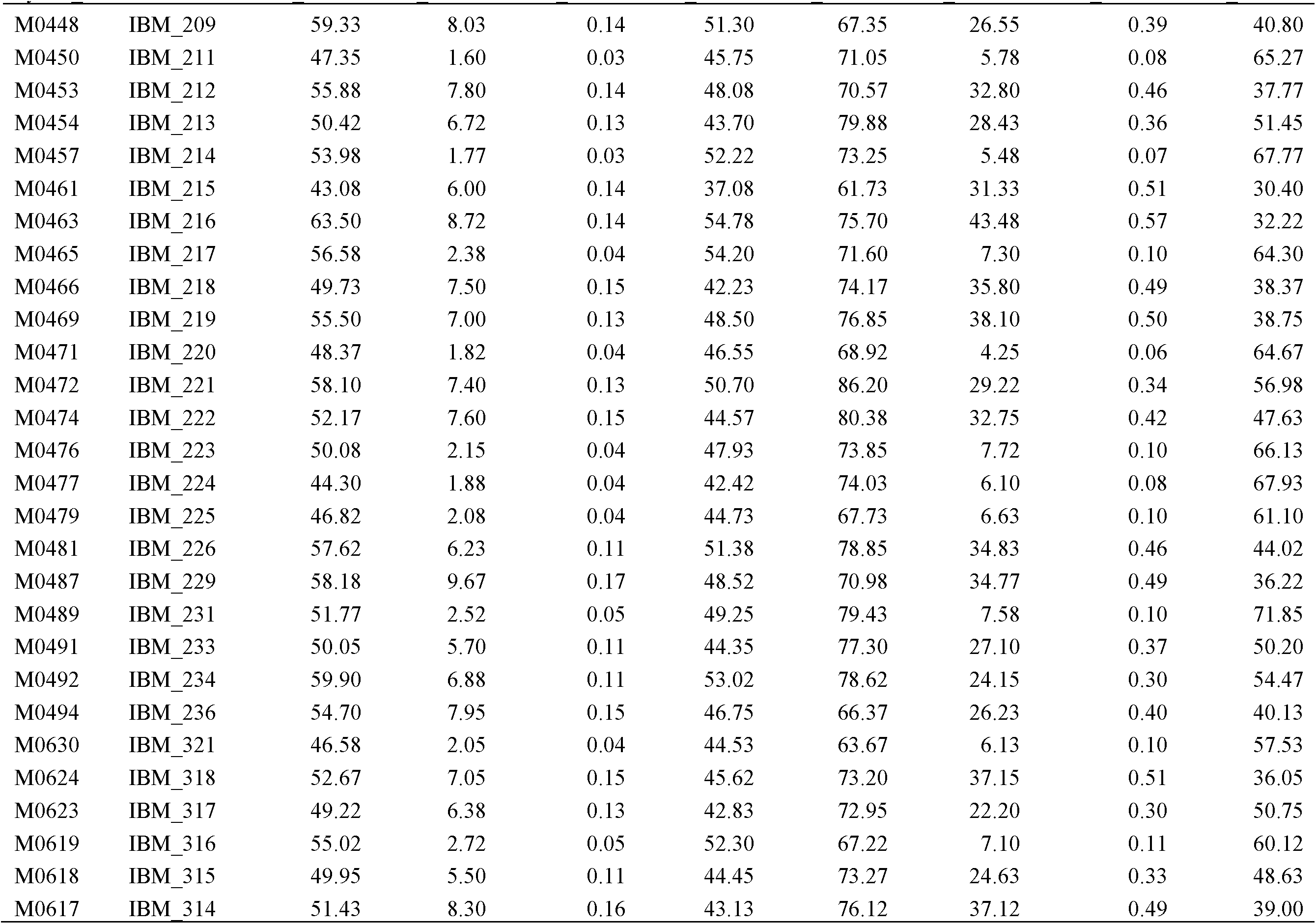
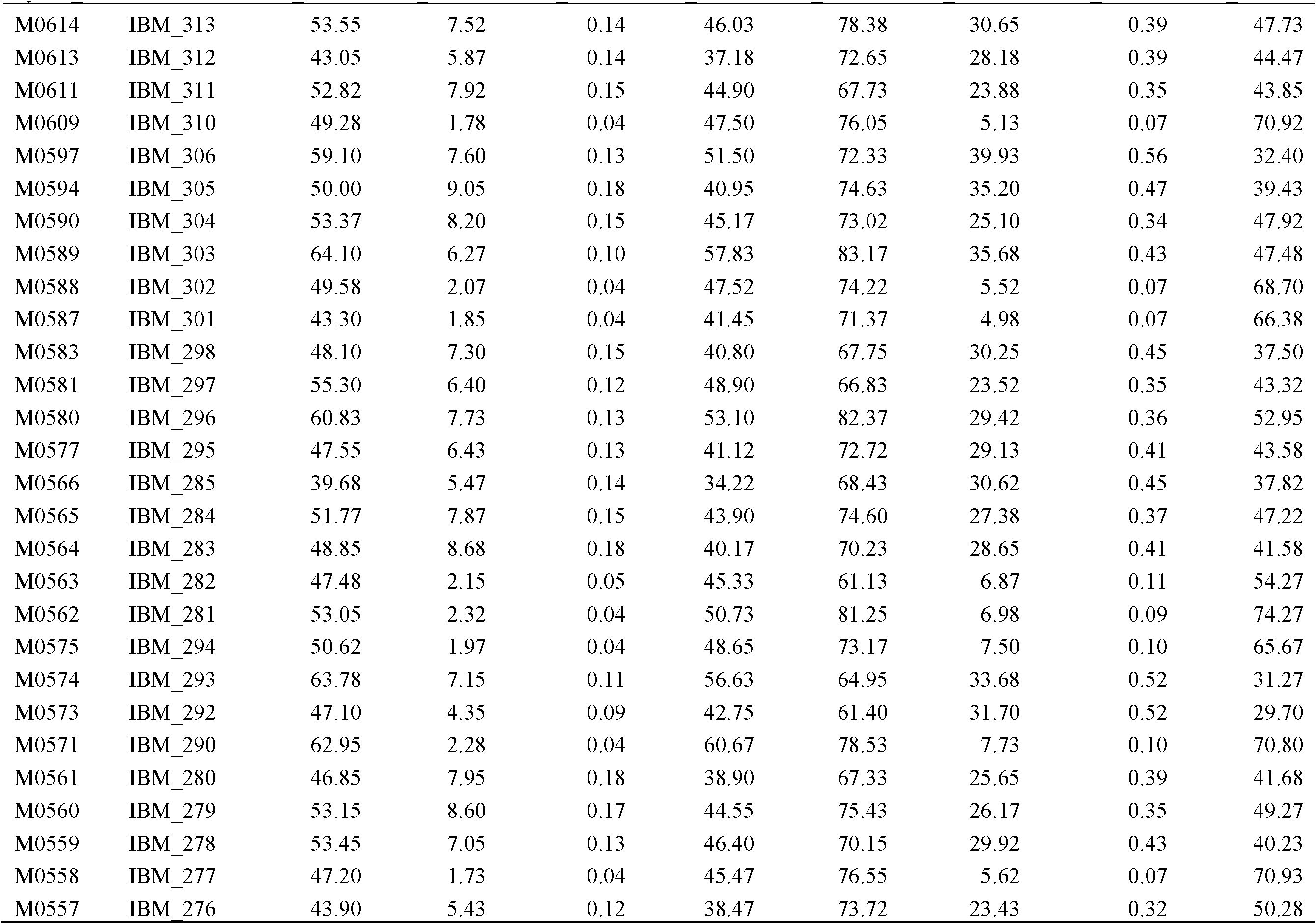
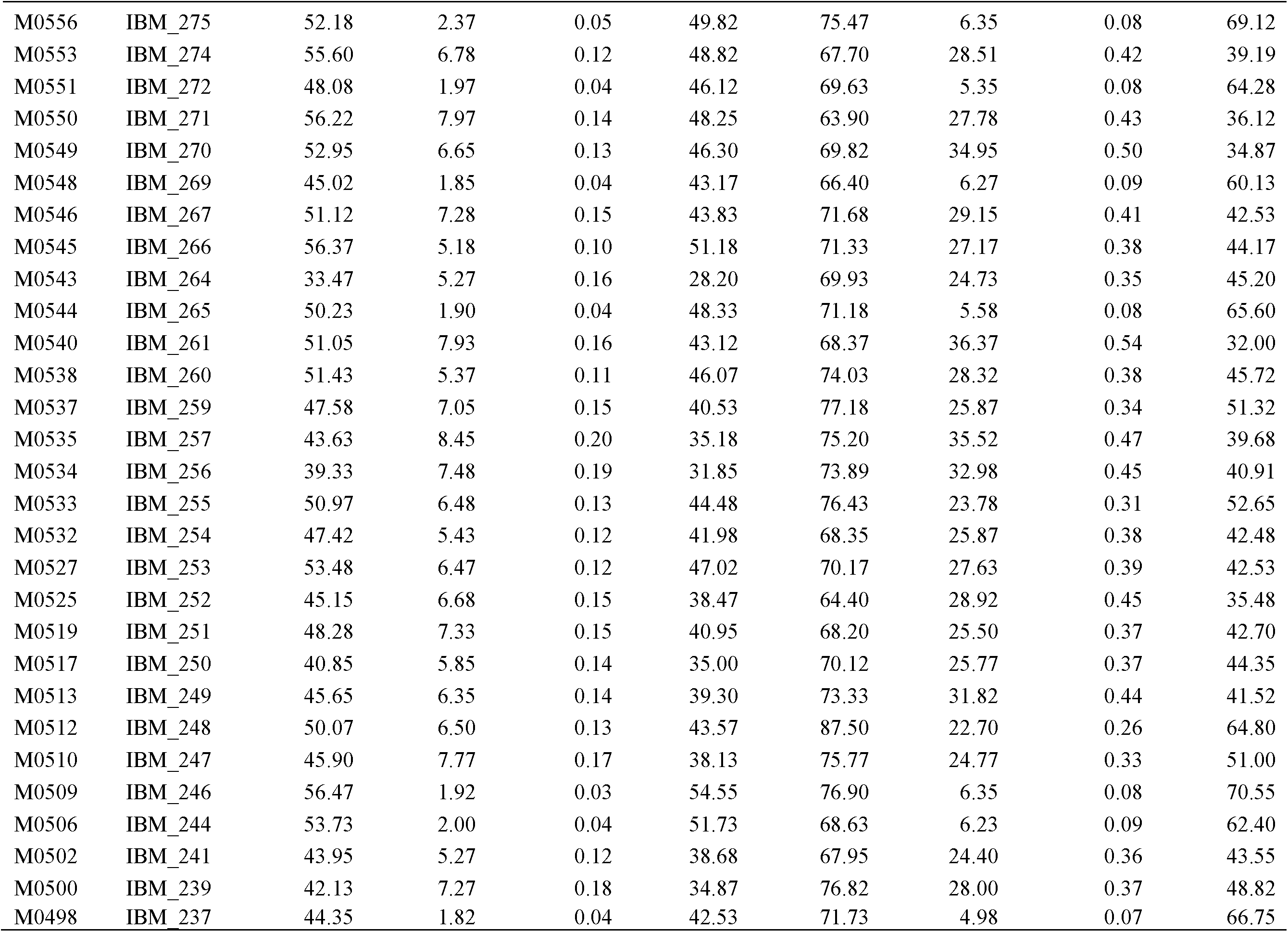
The trait mean values of the CCM traits for the wild-type (WT) and mutant (MT) siblings of F_1_ hybrids of *Oy1*-*N1989*/*oy1*:B73 (pollen-parent) with respective Syn10 line.

| Syn10_ID | Liu et.al-ID | WT_CCM I | MT_CCM I | Ratio_CCM I | Diff_CCM I | WT_CCM II | MT_CCM II | Ratio_CCM II | Diff_CCM II |
| --- | --- | --- | --- | --- | --- | --- | --- | --- | --- |
| M0001 | IBM_1 | 61.60 | 2.48 | 0.04 | 59.12 | 88.70 | 6.48 | 0.07 | 82.22 |
| M0002 | IBM_2 | 63.63 | 6.95 | 0.11 | 56.68 | 63.52 | 31.85 | 0.50 | 31.67 |
| M0009 | IBM_5 | 43.13 | 7.93 | 0.19 | 35.20 | 77.85 | 28.33 | 0.38 | 49.52 |
| M0010 | IBM_6 | 62.70 | 3.02 | 0.05 | 59.68 | 78.75 | 8.97 | 0.11 | 69.78 |
| M0011 | IBM_7 | 41.67 | 6.12 | 0.15 | 35.55 | 70.70 | 25.72 | 0.37 | 44.98 |
| M0012 | IBM_8 | 51.43 | 7.13 | 0.14 | 44.30 | 67.83 | 30.13 | 0.45 | 37.70 |
| M0015 | IBM_9 | 43.57 | 6.50 | 0.15 | 37.07 | 64.53 | 25.20 | 0.39 | 39.33 |
| M0019 | IBM_11 | 53.63 | 1.87 | 0.04 | 51.77 | 77.97 | 5.27 | 0.07 | 72.70 |
| M0020 | IBM_12 | 56.82 | 2.03 | 0.04 | 54.78 | 73.58 | 6.05 | 0.08 | 67.53 |
| M0023 | IBM_13 | 55.87 | 7.20 | 0.13 | 48.67 | 75.42 | 27.42 | 0.36 | 48.00 |
| M0024 | IBM_14 | 47.42 | 8.05 | 0.17 | 39.37 | 63.79 | 29.90 | 0.47 | 33.89 |
| M0025 | IBM_15 | 51.28 | 9.50 | 0.19 | 41.78 | 69.18 | 25.88 | 0.37 | 43.30 |
| M0027 | IBM_16 | 42.02 | 2.10 | 0.05 | 39.92 | 70.55 | 6.00 | 0.09 | 64.55 |
| M0029 | IBM_17 | 55.58 | 6.52 | 0.12 | 49.07 | 63.40 | 25.62 | 0.40 | 37.78 |
| M0030 | IBM_18 | 54.60 | 5.95 | 0.11 | 48.65 | 69.72 | 22.88 | 0.33 | 46.83 |
| M0032 | IBM_19 | 54.28 | 7.90 | 0.15 | 46.38 | 68.08 | 33.97 | 0.51 | 34.12 |
| M0034 | IBM_20 | 58.70 | 6.80 | 0.12 | 51.90 | 78.75 | 33.10 | 0.42 | 45.65 |
| M0036 | IBM_21 | 56.42 | 2.27 | 0.04 | 54.15 | 89.33 | 8.30 | 0.09 | 81.03 |
| M0038 | IBM_22 | 51.83 | 10.80 | 0.23 | 41.03 | 64.40 | 34.07 | 0.53 | 30.33 |
| M0041 | IBM_23 | 50.20 | 4.72 | 0.10 | 45.48 | 57.35 | 25.40 | 0.44 | 31.95 |
| M0042 | IBM_24 | 47.48 | 6.48 | 0.14 | 41.00 | 73.33 | 34.02 | 0.47 | 39.32 |
| M0043 | IBM_25 | 46.65 | 7.57 | 0.16 | 39.08 | 67.40 | 30.68 | 0.46 | 36.72 |
| M0047 | IBM_26 | 53.45 | 2.75 | 0.05 | 50.70 | 59.08 | 6.85 | 0.12 | 52.23 |
| M0049 | IBM_27 | 41.42 | 2.40 | 0.06 | 39.02 | 72.32 | 7.62 | 0.11 | 64.70 |
| M0050 | IBM_28 | 56.25 | 8.95 | 0.16 | 47.30 | 75.12 | 33.12 | 0.44 | 42.00 |
| M0053 | IBM_30 | 47.47 | 7.65 | 0.16 | 39.82 | 76.62 | 30.73 | 0.40 | 45.88 |

| Syn10_ID | Liu et.al-ID | WT_CCM1 | MT_CCM1 | Ratio_CCM1 | Diff_CCM1 | WT_CCM2 | MT_CCM2 | Ratio_CCM2 | Diff_CCM2 |
| --- | --- | --- | --- | --- | --- | --- | --- | --- | --- |
| M0054 | IBM_31 | 52.93 | 6.98 | 0.13 | 45.95 | 72.35 | 33.18 | 0.46 | 39.17 |
| M0055 | IBM_32 | 57.95 | 2.75 | 0.05 | 55.20 | 74.55 | 6.08 | 0.08 | 68.47 |
| M0056 | IBM_33 | 54.38 | 5.38 | 0.10 | 49.00 | 73.98 | 26.03 | 0.36 | 47.95 |
| M0057 | IBM_34 | 43.82 | 6.00 | 0.14 | 37.82 | 70.92 | 26.43 | 0.37 | 44.48 |
| M0061 | IBM_35 | 45.27 | 1.57 | 0.03 | 43.70 | 64.03 | 4.67 | 0.07 | 59.37 |
| M0062 | IBM_36 | 56.05 | 5.82 | 0.10 | 50.23 | 73.72 | 21.98 | 0.30 | 51.73 |
| M0068 | IBM_39 | 56.32 | 1.72 | 0.03 | 54.60 | 70.68 | 5.55 | 0.08 | 65.13 |
| M0069 | IBM_40 | 48.58 | 13.27 | 0.27 | 35.32 | 74.30 | 32.48 | 0.44 | 41.82 |
| M0070 | IBM_41 | 48.43 | 2.30 | 0.05 | 46.13 | 73.15 | 6.97 | 0.10 | 66.18 |
| M0074 | IBM_42 | 53.02 | 1.82 | 0.04 | 51.20 | 76.63 | 6.42 | 0.08 | 70.22 |
| M0076 | IBM_43 | 46.47 | 2.58 | 0.06 | 43.88 | 74.67 | 7.85 | 0.11 | 66.82 |
| M0078 | IBM_44 | 57.53 | 6.85 | 0.12 | 50.68 | 74.27 | 27.07 | 0.36 | 47.20 |
| M0084 | IBM_48 | 52.07 | 1.75 | 0.03 | 50.32 | 80.42 | 4.33 | 0.05 | 76.08 |
| M0090 | IBM_49 | 49.07 | 7.13 | 0.15 | 41.93 | 67.05 | 30.32 | 0.45 | 36.73 |
| M0091 | IBM_50 | 38.28 | 7.72 | 0.20 | 30.57 | 72.65 | 31.47 | 0.44 | 41.18 |
| M0092 | IBM_51 | 48.08 | 8.28 | 0.18 | 39.80 | 66.43 | 45.65 | 0.69 | 20.78 |
| M0093 | IBM_52 | 55.40 | 7.07 | 0.13 | 48.33 | 68.63 | 32.85 | 0.48 | 35.78 |
| M0095 | IBM_53 | 53.27 | 6.13 | 0.11 | 47.13 | 70.05 | 27.83 | 0.40 | 42.22 |
| M0105 | IBM_55 | 60.37 | 6.47 | 0.11 | 53.90 | 61.80 | 33.78 | 0.55 | 28.02 |
| M0108 | IBM_56 | 40.58 | 7.47 | 0.19 | 33.12 | 64.13 | 29.22 | 0.47 | 34.92 |
| M0109 | IBM_57 | 50.32 | 5.77 | 0.12 | 44.55 | 62.47 | 25.20 | 0.40 | 37.27 |
| M0111 | IBM_59 | 45.63 | 7.33 | 0.16 | 38.30 | 82.28 | 31.27 | 0.38 | 51.02 |
| M0116 | IBM_62 | 48.72 | 6.35 | 0.13 | 42.37 | 72.90 | 25.85 | 0.35 | 47.05 |
| M0118 | IBM_63 | 52.32 | 7.85 | 0.15 | 44.47 | 63.05 | 31.90 | 0.51 | 31.15 |
| M0119 | IBM_64 | 50.62 | 6.68 | 0.13 | 43.93 | 70.95 | 30.12 | 0.43 | 40.83 |
| M0120 | IBM_65 | 49.90 | 6.75 | 0.14 | 43.15 | 69.55 | 33.57 | 0.49 | 35.98 |
| M0126 | IBM_67 | 47.63 | 5.70 | 0.12 | 41.93 | 70.72 | 32.02 | 0.45 | 38.70 |
| M0130 | IBM_69 | 51.00 |  |  |  | 71.08 |  |  |  |
| M0136 | IBM_70 | 49.98 | 6.73 | 0.14 | 43.25 | 74.30 | 28.55 | 0.38 | 45.75 |
| M0317 | IBM_151 | 52.90 | 8.10 | 0.16 | 44.80 | 79.77 | 28.58 | 0.36 | 51.18 |
| M0316 | IBM_150 | 47.23 | 5.35 | 0.11 | 41.88 | 71.78 | 25.22 | 0.35 | 46.57 |
| M0315 | IBM_149 | 60.88 | 1.65 | 0.03 | 59.23 | 65.95 | 5.67 | 0.09 | 60.28 |
| M0312 | IBM_147 | 48.80 | 7.48 | 0.15 | 41.32 | 70.07 | 23.83 | 0.34 | 46.23 |
| M0305 | IBM_144 | 50.60 | 1.68 | 0.03 | 48.92 | 76.28 | 5.60 | 0.07 | 70.68 |
| M0298 | IBM_142 | 44.60 | 6.25 | 0.14 | 38.35 | 70.28 | 28.63 | 0.41 | 41.65 |
| M0296 | IBM_141 | 47.62 | 7.18 | 0.15 | 40.43 | 69.92 | 25.70 | 0.37 | 44.22 |
| M0295 | IBM_140 | 57.23 | 9.92 | 0.17 | 47.32 | 75.47 | 31.70 | 0.42 | 43.77 |
| M0293 | IBM_139 | 50.68 | 2.38 | 0.05 | 48.30 | 69.17 | 7.75 | 0.11 | 61.42 |
| M0290 | IBM_138 | 46.50 | 8.70 | 0.19 | 37.80 | 64.85 | 28.65 | 0.44 | 36.20 |
| M0285 | IBM_136 | 47.28 | 5.52 | 0.12 | 41.77 | 70.60 | 26.97 | 0.38 | 43.63 |
| M0272 | IBM_134 | 53.27 | 6.82 | 0.13 | 46.45 | 63.68 | 29.03 | 0.45 | 34.65 |
| M0267 | IBM_133 | 45.87 | 8.60 | 0.19 | 37.27 | 71.52 | 31.93 | 0.44 | 39.58 |
| M0266 | IBM_132 | 55.32 | 2.10 | 0.04 | 53.22 | 73.52 | 4.48 | 0.06 | 69.03 |
| M0264 | IBM_131 | 56.15 | 6.17 | 0.11 | 49.98 | 75.85 | 29.72 | 0.40 | 46.13 |
| M0262 | IBM_130 | 47.27 | 6.37 | 0.13 | 40.90 | 69.78 | 35.73 | 0.52 | 34.05 |
| M0258 | IBM_127 | 47.32 | 6.10 | 0.13 | 41.22 | 63.32 | 22.58 | 0.36 | 40.73 |
| M0256 | IBM_126 | 44.07 | 5.60 | 0.13 | 38.47 | 68.58 | 23.98 | 0.35 | 44.60 |
| M0253 | IBM_125 | 48.80 | 6.07 | 0.12 | 42.73 | 71.77 | 21.50 | 0.30 | 50.27 |
| M0252 | IBM_124 | 36.88 | 6.03 | 0.16 | 30.85 | 70.33 | 33.90 | 0.48 | 36.43 |
| M0250 | IBM_123 | 51.53 | 2.27 | 0.04 | 49.27 | 75.92 | 6.67 | 0.09 | 69.25 |
| M0249 | IBM_122 | 44.30 | 1.87 | 0.04 | 42.43 | 60.27 | 4.72 | 0.08 | 55.55 |
| M0245 | IBM_121 | 51.08 | 8.55 | 0.17 | 42.53 | 70.38 | 32.10 | 0.46 | 38.28 |
| M0243 | IBM_119 | 44.28 | 7.90 | 0.18 | 36.38 | 73.25 | 33.73 | 0.46 | 39.52 |
| M0240 | IBM_117 | 36.15 | 2.02 | 0.06 | 34.13 | 76.32 | 6.57 | 0.09 | 69.75 |
| M0237 | IBM_115 | 44.67 | 6.83 | 0.16 | 37.83 | 67.78 | 28.33 | 0.42 | 39.45 |
| M0234 | IBM_114 | 48.87 | 8.12 | 0.17 | 40.75 | 75.62 | 24.95 | 0.33 | 50.67 |
| M0231 | IBM_113 | 54.68 | 2.88 | 0.05 | 51.80 | 83.48 | 6.00 | 0.07 | 77.48 |
| M0230 | IBM_112 | 52.75 | 6.33 | 0.12 | 46.42 | 74.83 | 28.87 | 0.39 | 45.97 |
| M0227 | IBM_111 | 43.10 | 1.85 | 0.04 | 41.25 | 70.48 | 5.58 | 0.08 | 64.90 |
| M0223 | IBM_110 | 41.03 | 2.33 | 0.06 | 38.70 | 68.05 | 7.30 | 0.11 | 60.75 |
| M0221 | IBM_109 | 49.03 | 7.90 | 0.16 | 41.13 | 66.18 | 26.78 | 0.40 | 39.40 |
| M0218 | IBM_108 | 43.58 | 7.63 | 0.18 | 35.95 | 64.90 | 35.08 | 0.54 | 29.82 |
| M0217 | IBM_107 | 48.13 | 9.65 | 0.20 | 38.48 | 71.27 | 30.12 | 0.42 | 41.15 |
| M0215 | IBM_106 | 47.88 | 9.38 | 0.20 | 38.50 | 60.53 | 35.68 | 0.60 | 24.85 |
| M0212 | IBM_104 | 54.03 | 8.73 | 0.16 | 45.30 | 78.37 | 33.65 | 0.43 | 44.72 |
| M0210 | IBM_103 | 43.23 | 5.53 | 0.13 | 37.70 | 68.23 | 26.13 | 0.38 | 42.10 |
| M0209 | IBM_102 | 57.28 | 7.55 | 0.13 | 49.73 | 73.48 | 37.03 | 0.50 | 36.45 |
| M0208 | IBM_101 | 56.32 | 7.42 | 0.13 | 48.90 | 66.25 | 28.10 | 0.42 | 38.15 |
| M0206 | IBM_100 | 47.27 | 6.17 | 0.13 | 41.10 | 70.88 | 26.03 | 0.37 | 44.85 |
| M0202 | IBM_98 | 51.02 | 2.15 | 0.04 | 48.87 | 72.88 | 7.50 | 0.10 | 65.38 |
| M0198 | IBM_97 | 51.87 | 5.98 | 0.12 | 45.88 | 77.87 | 35.22 | 0.45 | 42.65 |
| M0196 | IBM_96 | 50.05 | 10.48 | 0.23 | 39.57 | 67.02 | 39.60 | 0.59 | 27.42 |
| M0195 | IBM_95 | 70.67 | 8.12 | 0.12 | 62.55 | 85.57 | 26.82 | 0.31 | 58.75 |
| M0193 | IBM_94 | 44.08 | 6.98 | 0.16 | 37.09 | 66.80 | 31.60 | 0.47 | 35.20 |
| M0189 | IBM_93 | 64.22 | 8.82 | 0.14 | 55.40 | 75.55 | 29.77 | 0.39 | 45.78 |
| M0188 | IBM_92 | 45.37 | 8.08 | 0.19 | 37.28 | 69.07 | 22.22 | 0.32 | 46.85 |
| M0184 | IBM_91 | 57.68 | 7.43 | 0.13 | 50.25 | 69.73 | 36.22 | 0.52 | 33.52 |
| M0179 | IBM_90 | 51.85 | 9.25 | 0.18 | 42.60 | 80.60 | 34.78 | 0.43 | 45.82 |
| M0178 | IBM_89 | 46.08 | 6.40 | 0.14 | 39.68 | 78.50 | 36.05 | 0.46 | 42.45 |
| M0175 | IBM_87 | 53.80 | 7.90 | 0.15 | 45.90 | 79.70 | 40.65 | 0.51 | 39.05 |
| M0173 | IBM_86 | 50.12 | 5.15 | 0.10 | 44.97 | 64.60 | 28.38 | 0.44 | 36.22 |
| M0172 | IBM_85 | 40.22 | 3.00 | 0.08 | 37.22 | 68.17 | 6.92 | 0.10 | 61.25 |
| M0171 | IBM_84 | 53.13 | 2.63 | 0.05 | 50.50 | 72.98 | 7.03 | 0.10 | 65.95 |
| M0169 | IBM_83 | 49.82 | 2.08 | 0.04 | 47.73 | 67.88 | 7.08 | 0.10 | 60.80 |

| Syn10_ID | Liu et.al-ID | WT_CCM1 | MT_CCM1 | Ratio_CCM1 | Diff_CCM1 | WT_CCM2 | MT_CCM2 | Ratio_CCM2 | Diff_CCM2 |
| --- | --- | --- | --- | --- | --- | --- | --- | --- | --- |
| M0166 | IBM_81 | 53.28 | 5.27 | 0.10 | 48.02 | 82.12 | 24.98 | 0.30 | 57.13 |
| M0162 | IBM_79 | 46.07 | 8.60 | 0.19 | 37.47 | 57.95 | 26.57 | 0.46 | 31.38 |
| M0159 | IBM_78 | 41.42 | 2.03 | 0.05 | 39.38 | 67.48 | 5.05 | 0.07 | 62.43 |
| M0156 | IBM_76 | 35.48 | 2.27 | 0.06 | 33.22 | 59.70 | 7.23 | 0.12 | 52.48 |
| M0155 | IBM_75 | 52.38 | 6.88 | 0.13 | 45.50 | 71.03 | 34.25 | 0.49 | 36.78 |
| M0152 | IBM_74 | 39.67 | 2.00 | 0.05 | 37.67 | 69.10 | 4.42 | 0.06 | 64.68 |
| M0145 | IBM_72 | 52.87 | 2.17 | 0.04 | 50.70 | 67.30 | 7.12 | 0.11 | 60.18 |
| M0138 | IBM_71 | 60.95 | 8.98 | 0.15 | 51.97 | 72.48 | 37.67 | 0.52 | 34.82 |
| M0319 | IBM_152 | 47.40 | 7.65 | 0.16 | 39.75 | 71.78 | 31.78 | 0.45 | 40.00 |
| M0324 | IBM_154 | 44.90 | 7.67 | 0.18 | 37.23 | 76.93 | 30.53 | 0.40 | 46.40 |
| M0328 | IBM_155 | 40.57 | 2.50 | 0.06 | 38.07 | 68.15 | 6.97 | 0.10 | 61.18 |
| M0330 | IBM_156 | 51.43 | 2.80 | 0.06 | 48.63 | 66.87 | 9.32 | 0.14 | 57.55 |
| M0332 | IBM_157 | 55.77 | 7.72 | 0.14 | 48.05 | 74.23 | 34.78 | 0.47 | 39.45 |
| M0335 | IBM_158 | 45.13 | 6.30 | 0.14 | 38.83 | 66.22 | 29.42 | 0.44 | 36.80 |
| M0336 | IBM_159 | 54.95 | 5.62 | 0.10 | 49.33 | 66.20 | 24.33 | 0.36 | 41.87 |
| M0340 | IBM_160 | 50.07 | 6.42 | 0.13 | 43.65 | 70.92 | 32.45 | 0.46 | 38.47 |
| M0341 | IBM_161 | 45.77 | 5.88 | 0.13 | 39.88 | 66.17 | 29.02 | 0.44 | 37.15 |
| M0345 | IBM_163 | 39.53 | 2.00 | 0.05 | 37.53 | 66.82 | 5.43 | 0.08 | 61.38 |
| M0347 | IBM_165 | 47.03 | 6.58 | 0.14 | 40.45 | 78.90 | 26.65 | 0.34 | 52.25 |
| M0349 | IBM_166 | 48.20 | 7.65 | 0.16 | 40.55 | 80.10 | 33.42 | 0.42 | 46.68 |
| M0356 | IBM_167 | 51.57 | 1.57 | 0.03 | 50.00 | 75.72 | 5.28 | 0.07 | 70.43 |
| M0357 | IBM_168 | 44.47 | 5.68 | 0.13 | 38.78 | 69.68 | 23.17 | 0.33 | 46.52 |
| M0358 | IBM_169 | 67.85 | 5.88 | 0.09 | 61.97 | 82.13 | 34.42 | 0.42 | 47.72 |
| M0366 | IBM_170 | 55.93 | 2.47 | 0.04 | 53.47 | 73.63 | 6.35 | 0.09 | 67.28 |
| M0372 | IBM_171 | 51.37 | 9.13 | 0.18 | 42.23 | 77.35 | 34.88 | 0.45 | 42.47 |
| M0373 | IBM_172 | 37.92 | 1.78 | 0.05 | 36.13 | 72.35 | 4.80 | 0.07 | 67.55 |
| M0374 | IBM_173 | 41.17 | 1.80 | 0.05 | 39.37 | 66.05 | 5.58 | 0.09 | 60.47 |
| M0375 | IBM_174 | 46.17 | 6.23 | 0.14 | 39.93 | 67.58 | 29.42 | 0.44 | 38.16 |
| M0376 | IBM_175 | 58.53 | 2.32 | 0.04 | 56.22 | 74.37 | 8.18 | 0.11 | 66.18 |
| M0381 | IBM_178 | 50.28 | 2.27 | 0.04 | 48.02 | 57.35 | 6.95 | 0.12 | 50.40 |
| M0382 | IBM_179 | 52.32 | 4.85 | 0.09 | 47.47 | 77.27 | 21.68 | 0.29 | 55.58 |
| M0385 | IBM_180 | 34.77 | 5.25 | 0.15 | 29.52 | 71.38 | 24.52 | 0.34 | 46.87 |
| M0388 | IBM_181 | 60.67 | 6.83 | 0.11 | 53.83 | 71.70 | 26.62 | 0.37 | 45.08 |
| M0389 | IBM_182 | 48.78 | 7.87 | 0.16 | 40.92 | 70.38 | 29.97 | 0.43 | 40.42 |
| M0390 | IBM_183 | 39.53 | 1.70 | 0.04 | 37.83 | 72.97 | 5.02 | 0.07 | 67.95 |
| M0391 | IBM_184 | 52.42 | 2.32 | 0.05 | 50.10 | 79.38 | 6.62 | 0.08 | 72.77 |
| M0396 | IBM_185 | 51.53 | 6.05 | 0.12 | 45.48 | 79.13 | 33.27 | 0.42 | 45.87 |
| M0401 | IBM_187 | 60.02 | 7.20 | 0.12 | 52.82 | 71.75 | 32.05 | 0.45 | 39.70 |
| M0404 | IBM_188 | 53.32 | 1.77 | 0.03 | 51.55 | 75.68 | 5.95 | 0.08 | 69.73 |
| M0405 | IBM_189 | 53.60 | 1.93 | 0.04 | 51.67 | 70.52 | 6.07 | 0.09 | 64.45 |
| M0407 | IBM_190 | 47.50 | 7.75 | 0.17 | 39.75 | 77.98 | 30.65 | 0.39 | 47.33 |
| M0412 | IBM_191 | 37.87 | 7.05 | 0.19 | 30.82 | 74.98 | 30.13 | 0.40 | 44.85 |
| M0414 | IBM_192 | 42.75 | 6.57 | 0.16 | 36.18 | 68.87 | 27.23 | 0.39 | 41.63 |
| M0422 | IBM_194 | 47.47 | 1.82 | 0.04 | 45.65 | 77.98 | 7.70 | 0.10 | 70.28 |
| M0424 | IBM_195 | 45.82 | 4.88 | 0.11 | 40.93 | 67.33 | 23.18 | 0.35 | 44.15 |
| M0427 | IBM_196 | 47.42 | 8.50 | 0.18 | 38.92 | 77.37 | 39.40 | 0.51 | 37.97 |
| M0428 | IBM_197 | 49.53 | 8.77 | 0.18 | 40.77 | 74.85 | 30.78 | 0.41 | 44.07 |
| M0429 | IBM_198 | 51.37 | 6.95 | 0.14 | 44.42 | 74.28 | 29.80 | 0.40 | 44.48 |
| M0430 | IBM_199 | 42.72 | 8.85 | 0.20 | 33.87 | 74.15 | 30.12 | 0.41 | 44.03 |
| M0431 | IBM_200 | 46.80 | 1.98 | 0.04 | 44.82 | 77.30 | 6.02 | 0.08 | 71.28 |
| M0433 | IBM_201 | 57.42 | 8.47 | 0.15 | 48.95 | 79.95 | 32.02 | 0.40 | 47.93 |
| M0434 | IBM_202 | 48.87 | 6.88 | 0.14 | 41.98 | 68.75 | 28.70 | 0.42 | 40.05 |
| M0436 | IBM_203 | 39.53 | 5.48 | 0.14 | 34.05 | 68.33 | 25.87 | 0.38 | 42.47 |
| M0438 | IBM_204 | 52.88 | 9.07 | 0.17 | 43.82 | 69.20 | 33.87 | 0.49 | 35.33 |
| M0443 | IBM_206 | 58.77 | 1.98 | 0.03 | 56.78 | 72.68 | 5.62 | 0.08 | 67.07 |
| M0444 | IBM_207 | 44.07 | 5.62 | 0.13 | 38.45 | 68.47 | 26.90 | 0.40 | 41.57 |
| M0448 | IBM_209 | 59.33 | 8.03 | 0.14 | 51.30 | 67.35 | 26.55 | 0.39 | 40.80 |
| M0450 | IBM_211 | 47.35 | 1.60 | 0.03 | 45.75 | 71.05 | 5.78 | 0.08 | 65.27 |
| M0453 | IBM_212 | 55.88 | 7.80 | 0.14 | 48.08 | 70.57 | 32.80 | 0.46 | 37.77 |
| M0454 | IBM_213 | 50.42 | 6.72 | 0.13 | 43.70 | 79.88 | 28.43 | 0.36 | 51.45 |
| M0457 | IBM_214 | 53.98 | 1.77 | 0.03 | 52.22 | 73.25 | 5.48 | 0.07 | 67.77 |
| M0461 | IBM_215 | 43.08 | 6.00 | 0.14 | 37.08 | 61.73 | 31.33 | 0.51 | 30.40 |
| M0463 | IBM_216 | 63.50 | 8.72 | 0.14 | 54.78 | 75.70 | 43.48 | 0.57 | 32.22 |
| M0465 | IBM_217 | 56.58 | 2.38 | 0.04 | 54.20 | 71.60 | 7.30 | 0.10 | 64.30 |
| M0466 | IBM_218 | 49.73 | 7.50 | 0.15 | 42.23 | 74.17 | 35.80 | 0.49 | 38.37 |
| M0469 | IBM_219 | 55.50 | 7.00 | 0.13 | 48.50 | 76.85 | 38.10 | 0.50 | 38.75 |
| M0471 | IBM_220 | 48.37 | 1.82 | 0.04 | 46.55 | 68.92 | 4.25 | 0.06 | 64.67 |
| M0472 | IBM_221 | 58.10 | 7.40 | 0.13 | 50.70 | 86.20 | 29.22 | 0.34 | 56.98 |
| M0474 | IBM_222 | 52.17 | 7.60 | 0.15 | 44.57 | 80.38 | 32.75 | 0.42 | 47.63 |
| M0476 | IBM_223 | 50.08 | 2.15 | 0.04 | 47.93 | 73.85 | 7.72 | 0.10 | 66.13 |
| M0477 | IBM_224 | 44.30 | 1.88 | 0.04 | 42.42 | 74.03 | 6.10 | 0.08 | 67.93 |
| M0479 | IBM_225 | 46.82 | 2.08 | 0.04 | 44.73 | 67.73 | 6.63 | 0.10 | 61.10 |
| M0481 | IBM_226 | 57.62 | 6.23 | 0.11 | 51.38 | 78.85 | 34.83 | 0.46 | 44.02 |
| M0487 | IBM_229 | 58.18 | 9.67 | 0.17 | 48.52 | 70.98 | 34.77 | 0.49 | 36.22 |
| M0489 | IBM_231 | 51.77 | 2.52 | 0.05 | 49.25 | 79.43 | 7.58 | 0.10 | 71.85 |
| M0491 | IBM_233 | 50.05 | 5.70 | 0.11 | 44.35 | 77.30 | 27.10 | 0.37 | 50.20 |
| M0492 | IBM_234 | 59.90 | 6.88 | 0.11 | 53.02 | 78.62 | 24.15 | 0.30 | 54.47 |
| M0494 | IBM_236 | 54.70 | 7.95 | 0.15 | 46.75 | 66.37 | 26.23 | 0.40 | 40.13 |
| M0630 | IBM_321 | 46.58 | 2.05 | 0.04 | 44.53 | 63.67 | 6.13 | 0.10 | 57.53 |
| M0624 | IBM_318 | 52.67 | 7.05 | 0.15 | 45.62 | 73.20 | 37.15 | 0.51 | 36.05 |
| M0623 | IBM_317 | 49.22 | 6.38 | 0.13 | 42.83 | 72.95 | 22.20 | 0.30 | 50.75 |
| M0619 | IBM_316 | 55.02 | 2.72 | 0.05 | 52.30 | 67.22 | 7.10 | 0.11 | 60.12 |
| M0618 | IBM_315 | 49.95 | 5.50 | 0.11 | 44.45 | 73.27 | 24.63 | 0.33 | 48.63 |
| M0617 | IBM_314 | 51.43 | 8.30 | 0.16 | 43.13 | 76.12 | 37.12 | 0.49 | 39.00 |
| M0614 | IBM_313 | 53.55 | 7.52 | 0.14 | 46.03 | 78.38 | 30.65 | 0.39 | 47.73 |
| M0613 | IBM_312 | 43.05 | 5.87 | 0.14 | 37.18 | 72.65 | 28.18 | 0.39 | 44.47 |
| M0611 | IBM_311 | 52.82 | 7.92 | 0.15 | 44.90 | 67.73 | 23.88 | 0.35 | 43.85 |
| M0609 | IBM_310 | 49.28 | 1.78 | 0.04 | 47.50 | 76.05 | 5.13 | 0.07 | 70.92 |
| M0597 | IBM_306 | 59.10 | 7.60 | 0.13 | 51.50 | 72.33 | 39.93 | 0.56 | 32.40 |
| M0594 | IBM_305 | 50.00 | 9.05 | 0.18 | 40.95 | 74.63 | 35.20 | 0.47 | 39.43 |
| M0590 | IBM_304 | 53.37 | 8.20 | 0.15 | 45.17 | 73.02 | 25.10 | 0.34 | 47.92 |
| M0589 | IBM_303 | 64.10 | 6.27 | 0.10 | 57.83 | 83.17 | 35.68 | 0.43 | 47.48 |
| M0588 | IBM_302 | 49.58 | 2.07 | 0.04 | 47.52 | 74.22 | 5.52 | 0.07 | 68.70 |
| M0587 | IBM_301 | 43.30 | 1.85 | 0.04 | 41.45 | 71.37 | 4.98 | 0.07 | 66.38 |
| M0583 | IBM_298 | 48.10 | 7.30 | 0.15 | 40.80 | 67.75 | 30.25 | 0.45 | 37.50 |
| M0581 | IBM_297 | 55.30 | 6.40 | 0.12 | 48.90 | 66.83 | 23.52 | 0.35 | 43.32 |
| M0580 | IBM_296 | 60.83 | 7.73 | 0.13 | 53.10 | 82.37 | 29.42 | 0.36 | 52.95 |
| M0577 | IBM_295 | 47.55 | 6.43 | 0.13 | 41.12 | 72.72 | 29.13 | 0.41 | 43.58 |
| M0566 | IBM_285 | 39.68 | 5.47 | 0.14 | 34.22 | 68.43 | 30.62 | 0.45 | 37.82 |
| M0565 | IBM_284 | 51.77 | 7.87 | 0.15 | 43.90 | 74.60 | 27.38 | 0.37 | 47.22 |
| M0564 | IBM_283 | 48.85 | 8.68 | 0.18 | 40.17 | 70.23 | 28.65 | 0.41 | 41.58 |
| M0563 | IBM_282 | 47.48 | 2.15 | 0.05 | 45.33 | 61.13 | 6.87 | 0.11 | 54.27 |
| M0562 | IBM_281 | 53.05 | 2.32 | 0.04 | 50.73 | 81.25 | 6.98 | 0.09 | 74.27 |
| M0575 | IBM_294 | 50.62 | 1.97 | 0.04 | 48.65 | 73.17 | 7.50 | 0.10 | 65.67 |
| M0574 | IBM_293 | 63.78 | 7.15 | 0.11 | 56.63 | 64.95 | 33.68 | 0.52 | 31.27 |
| M0573 | IBM_292 | 47.10 | 4.35 | 0.09 | 42.75 | 61.40 | 31.70 | 0.52 | 29.70 |
| M0571 | IBM_290 | 62.95 | 2.28 | 0.04 | 60.67 | 78.53 | 7.73 | 0.10 | 70.80 |
| M0561 | IBM_280 | 46.85 | 7.95 | 0.18 | 38.90 | 67.33 | 25.65 | 0.39 | 41.68 |
| M0560 | IBM_279 | 53.15 | 8.60 | 0.17 | 44.55 | 75.43 | 26.17 | 0.35 | 49.27 |
| M0559 | IBM_278 | 53.45 | 7.05 | 0.13 | 46.40 | 70.15 | 29.92 | 0.43 | 40.23 |
| M0558 | IBM_277 | 47.20 | 1.73 | 0.04 | 45.47 | 76.55 | 5.62 | 0.07 | 70.93 |
| M0557 | IBM_276 | 43.90 | 5.43 | 0.12 | 38.47 | 73.72 | 23.43 | 0.32 | 50.28 |
| M0556 | IBM_275 | 52.18 | 2.37 | 0.05 | 49.82 | 75.47 | 6.35 | 0.08 | 69.12 |
| M0553 | IBM_274 | 55.60 | 6.78 | 0.12 | 48.82 | 67.70 | 28.51 | 0.42 | 39.19 |
| M0551 | IBM_272 | 48.08 | 1.97 | 0.04 | 46.12 | 69.63 | 5.35 | 0.08 | 64.28 |
| M0550 | IBM_271 | 56.22 | 7.97 | 0.14 | 48.25 | 63.90 | 27.78 | 0.43 | 36.12 |
| M0549 | IBM_270 | 52.95 | 6.65 | 0.13 | 46.30 | 69.82 | 34.95 | 0.50 | 34.87 |
| M0548 | IBM_269 | 45.02 | 1.85 | 0.04 | 43.17 | 66.40 | 6.27 | 0.09 | 60.13 |
| M0546 | IBM_267 | 51.12 | 7.28 | 0.15 | 43.83 | 71.68 | 29.15 | 0.41 | 42.53 |
| M0545 | IBM_266 | 56.37 | 5.18 | 0.10 | 51.18 | 71.33 | 27.17 | 0.38 | 44.17 |
| M0543 | IBM_264 | 33.47 | 5.27 | 0.16 | 28.20 | 69.93 | 24.73 | 0.35 | 45.20 |
| M0544 | IBM_265 | 50.23 | 1.90 | 0.04 | 48.33 | 71.18 | 5.58 | 0.08 | 65.60 |
| M0540 | IBM_261 | 51.05 | 7.93 | 0.16 | 43.12 | 68.37 | 36.37 | 0.54 | 32.00 |
| M0538 | IBM_260 | 51.43 | 5.37 | 0.11 | 46.07 | 74.03 | 28.32 | 0.38 | 45.72 |
| M0537 | IBM_259 | 47.58 | 7.05 | 0.15 | 40.53 | 77.18 | 25.87 | 0.34 | 51.32 |
| M0535 | IBM_257 | 43.63 | 8.45 | 0.20 | 35.18 | 75.20 | 35.52 | 0.47 | 39.68 |
| M0534 | IBM_256 | 39.33 | 7.48 | 0.19 | 31.85 | 73.89 | 32.98 | 0.45 | 40.91 |
| M0533 | IBM_255 | 50.97 | 6.48 | 0.13 | 44.48 | 76.43 | 23.78 | 0.31 | 52.65 |
| M0532 | IBM_254 | 47.42 | 5.43 | 0.12 | 41.98 | 68.35 | 25.87 | 0.38 | 42.48 |
| M0527 | IBM_253 | 53.48 | 6.47 | 0.12 | 47.02 | 70.17 | 27.63 | 0.39 | 42.53 |
| M0525 | IBM_252 | 45.15 | 6.68 | 0.15 | 38.47 | 64.40 | 28.92 | 0.45 | 35.48 |
| M0519 | IBM_251 | 48.28 | 7.33 | 0.15 | 40.95 | 68.20 | 25.50 | 0.37 | 42.70 |
| M0517 | IBM_250 | 40.85 | 5.85 | 0.14 | 35.00 | 70.12 | 25.77 | 0.37 | 44.35 |
| M0513 | IBM_249 | 45.65 | 6.35 | 0.14 | 39.30 | 73.33 | 31.82 | 0.44 | 41.52 |
| M0512 | IBM_248 | 50.07 | 6.50 | 0.13 | 43.57 | 87.50 | 22.70 | 0.26 | 64.80 |
| M0510 | IBM_247 | 45.90 | 7.77 | 0.17 | 38.13 | 75.77 | 24.77 | 0.33 | 51.00 |
| M0509 | IBM_246 | 56.47 | 1.92 | 0.03 | 54.55 | 76.90 | 6.35 | 0.08 | 70.55 |
| M0506 | IBM_244 | 53.73 | 2.00 | 0.04 | 51.73 | 68.63 | 6.23 | 0.09 | 62.40 |
| M0502 | IBM_241 | 43.95 | 5.27 | 0.12 | 38.68 | 67.95 | 24.40 | 0.36 | 43.55 |
| M0500 | IBM_239 | 42.13 | 7.27 | 0.18 | 34.87 | 76.82 | 28.00 | 0.37 | 48.82 |

| Syn10_ID | Liu et.al-ID | WT_CCMI | MT_CCMI | Ratio_CCMI | Diff_CCMI | WT_CCMII | MT_CCMII | Ratio_CCMII | Diff_CCMII |
| --- | --- | --- | --- | --- | --- | --- | --- | --- | --- |
| M0498 | IBM_237 | 44.35 | 1.82 | 0.04 | 42.53 | 71.73 | 4.98 | 0.07 | 66.75 |

**Table S3.** The average values of the CCM traits in wild-type (WT) and mutant (MT) siblings of the F_1_ hybrids between *Oy1-N1989/oy1*:B73 (as a pollen-parent) with respective BM-NILs. Data is derived from the field-grown plants with five replications planted in a RCBD. Parental (B73 and Mo17) F_1_ crosses were planted as checks in each replication. Multiple plants (2-3) were measured for each genotype (wild-type or mutant) in each replication.

| Ear-parent | NIL background | *vey1* status | MT_CCMI | WT_CCMI | Ratio_CCMI | MT_CCMII | WT_CCMII | Ratio_CCMII |
| --- | --- | --- | --- | --- | --- | --- | --- | --- |
| **B73** | . | **B73** | **4.75** | **27.49** | **0.17** | **22.15** | **64.55** | **0.34** |
| **Mo17** | . | **Mo17** | **2.05** | **31.17** | **0.07** | **5.69** | **78.18** | **0.07** |
| b142 | B73 | B73 | 6.07 | 30.21 | 0.20 | 25.26 | 63.34 | 0.40 |
| b139 | B73 | B73 | 6.26 | 34.61 | 0.18 | 25.25 | 67.75 | 0.37 |
| b135 | B73 | B73 | 5.73 | 30.79 | 0.19 | 27.88 | 70.03 | 0.40 |
| b132 | B73 | B73 | 4.95 | 31.06 | 0.17 | 24.91 | 63.42 | 0.40 |
| b125 | B73 | B73 | 7.17 | 37.01 | 0.19 | 27.15 | 65.83 | 0.41 |
| b185 | B73 | B73 | 5.89 | 33.60 | 0.18 | 26.88 | 66.63 | 0.41 |
| b155 | B73 | B73 | 6.17 | 30.34 | 0.21 | 25.64 | 58.59 | 0.45 |
| b121 | B73 | B73 | 5.51 | 32.37 | 0.18 | 27.26 | 70.14 | 0.39 |
| b120 | B73 | B73 | 6.75 | 34.16 | 0.20 | 30.36 | 65.58 | 0.48 |
| b092 | B73 | B73 | 5.03 | 30.71 | 0.17 | 24.97 | 63.72 | 0.40 |
| b087 | B73 | B73 | 5.03 | 28.70 | 0.18 | 24.13 | 66.23 | 0.37 |
| b055 | B73 | B73 | 6.79 | 29.30 | 0.24 | 29.99 | 65.63 | 0.46 |
| b035 | B73 | B73 | 5.55 | 34.25 | 0.17 | 26.87 | 71.57 | 0.38 |
| b030 | B73 | B73 | 6.15 | 32.85 | 0.19 | 32.26 | 73.97 | 0.44 |
| b123 | B73 | Mo17 | 2.18 | 31.38 | 0.07 | 5.63 | 71.50 | 0.08 |
| b189 | B73 | Mo17 | 2.67 | 29.97 | 0.09 | 6.01 | 68.55 | 0.09 |
| b182 | B73 | Mo17 | 2.24 | 28.76 | 0.08 | 5.39 | 63.71 | 0.09 |
| b107 | B73 | Mo17 | 1.97 | 33.01 | 0.06 | 5.91 | 75.01 | 0.08 |
| b094 | B73 | Mo17 | 1.71 | 32.26 | 0.05 | 5.34 | 73.49 | 0.07 |
| b049 | B73 | Mo17 | 2.32 | 30.75 | 0.08 | 6.41 | 68.29 | 0.09 |
| b047 | B73 | Mo17 | 1.85 | 31.44 | 0.06 | 6.29 | 71.44 | 0.09 |
| b001 | B73 | Mo17 | 2.03 | 33.29 | 0.06 | 5.32 | 71.58 | 0.07 |

| Ear-parent | NIL background | *veyI* status | MT_CCMI | WT_CCMI | Ratio_CCMI | MT_CCMII | WT_CCMII | Ratio_CCMII |
| --- | --- | --- | --- | --- | --- | --- | --- | --- |
| mo24 | Mo17 | B73 | 5.73 | 29.76 | 0.20 | 30.65 | 68.89 | 0.45 |
| m097 | Mo17 | B73 | 5.53 | 35.29 | 0.16 | 29.03 | 72.02 | 0.41 |
| mo27 | Mo17 | Mo17 | 2.31 | 28.50 | 0.08 | 7.04 | 71.96 | 0.10 |
| m022 | Mo17 | Mo17 | 2.13 | 30.65 | 0.07 | 6.57 | 71.69 | 0.09 |
| m008 | Mo17 | Mo17 | 2.37 | 29.75 | 0.08 | 7.20 | 76.88 | 0.09 |
| m002 | Mo17 | Mo17 | 2.39 | 31.98 | 0.08 | 6.49 | 66.28 | 0.10 |
| m093 | Mo17 | Mo17 | 1.91 | 30.85 | 0.06 | 7.26 | 68.09 | 0.11 |
| m079 | Mo17 | Mo17 | 2.31 | 29.71 | 0.08 | 8.50 | 68.21 | 0.13 |
| m051 | Mo17 | Mo17 | 2.23 | 30.47 | 0.07 | 6.04 | 69.15 | 0.09 |
| m048 | Mo17 | Mo17 | 2.53 | 34.06 | 0.08 | 7.22 | 74.83 | 0.10 |
| m043 | Mo17 | Mo17 | 1.85 | 26.98 | 0.07 | 6.76 | 81.51 | 0.08 |
| m038 | Mo17 | Mo17 | 2.25 | 29.24 | 0.08 | 6.98 | 75.18 | 0.09 |
| m035 | Mo17 | Mo17 | 2.03 | 31.19 | 0.07 | 7.15 | 73.66 | 0.10 |

**Table S4.**
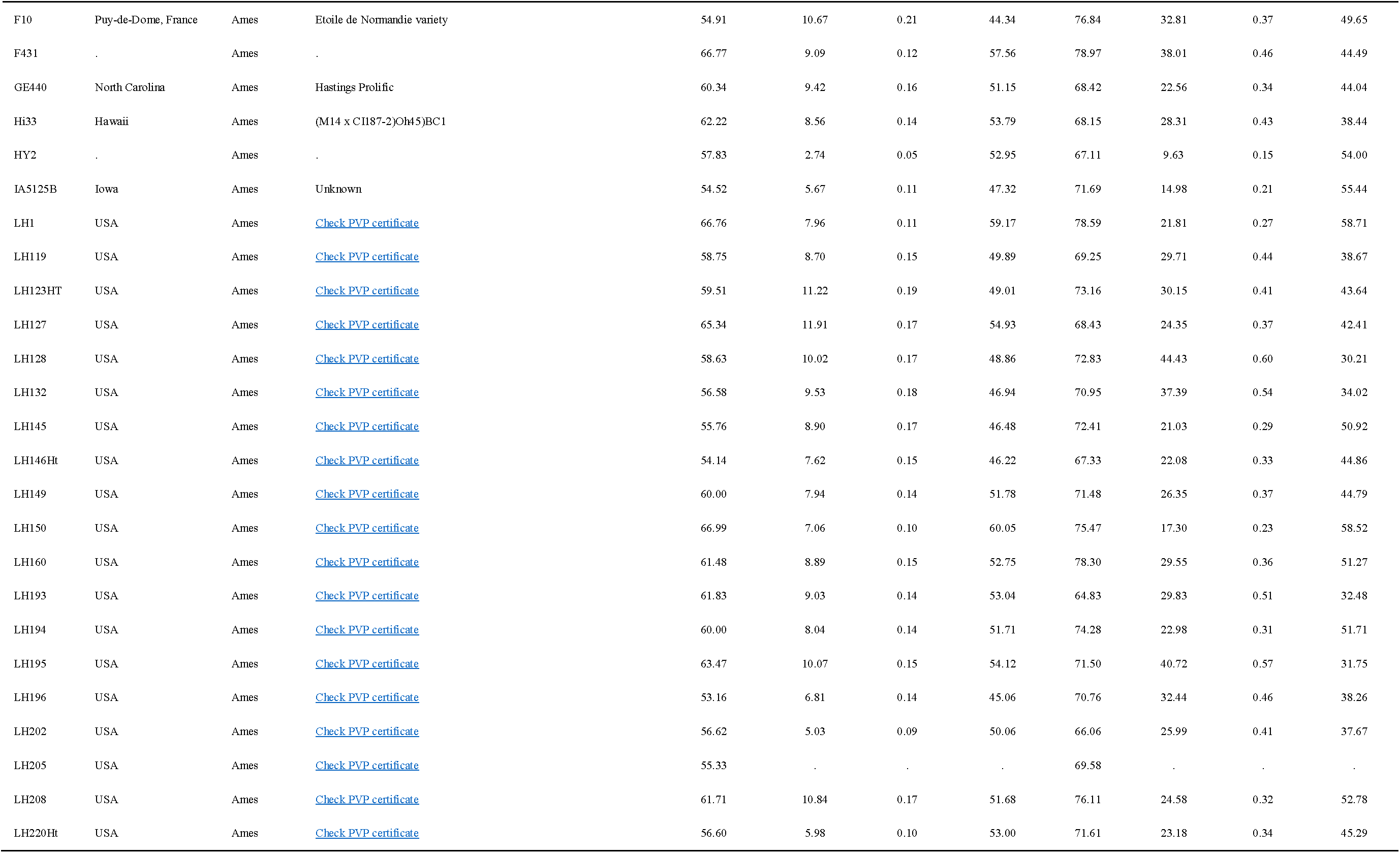
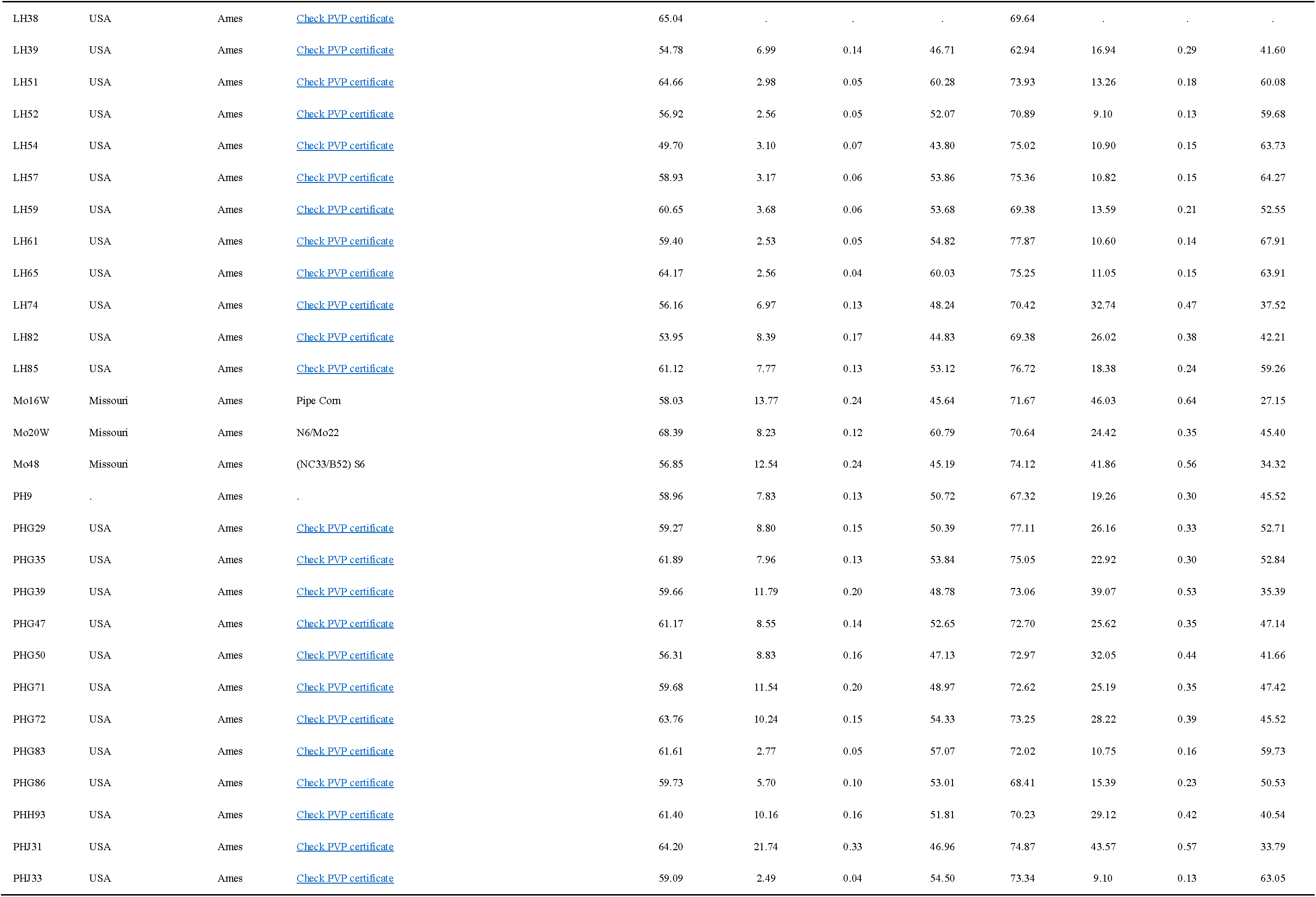
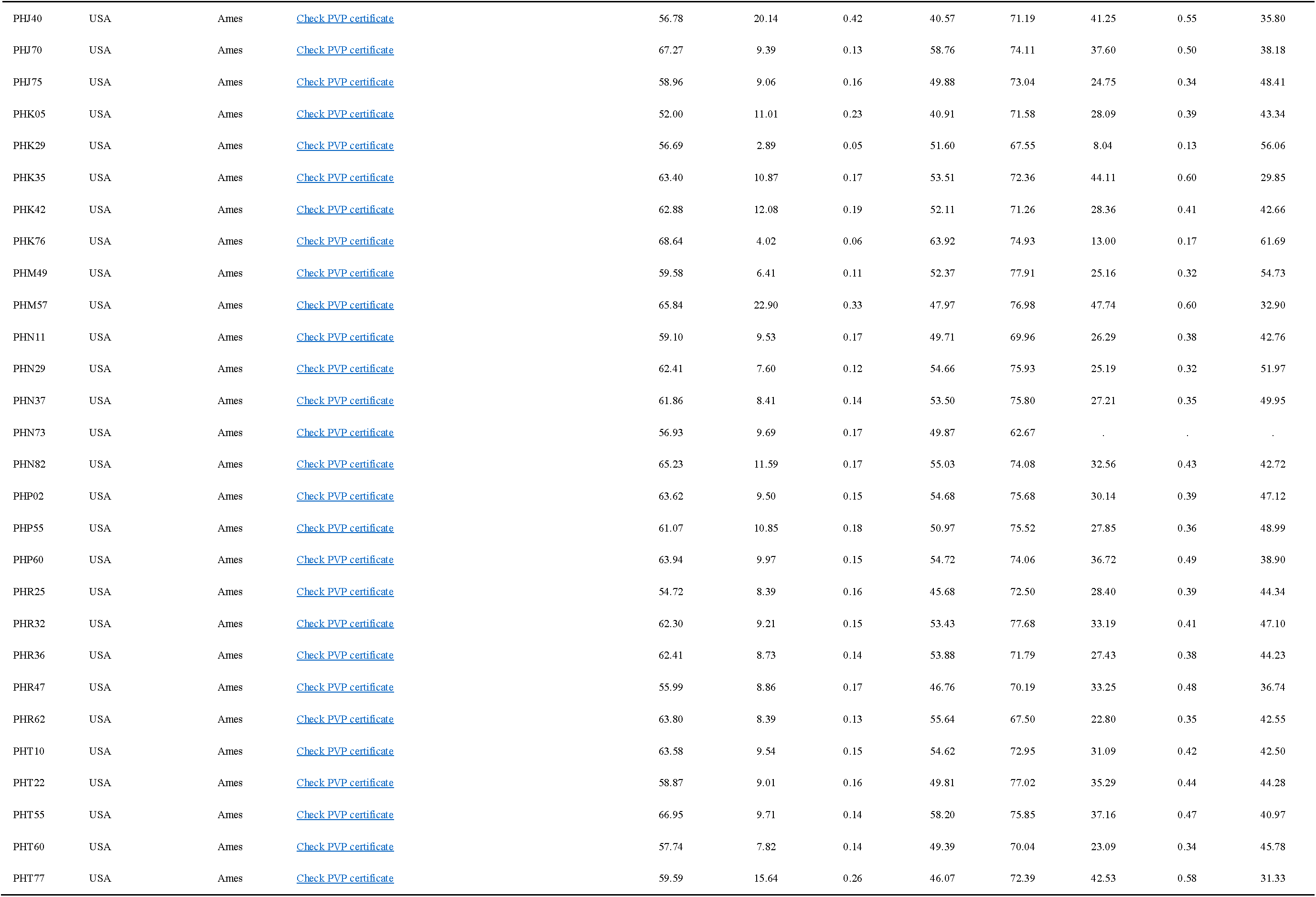
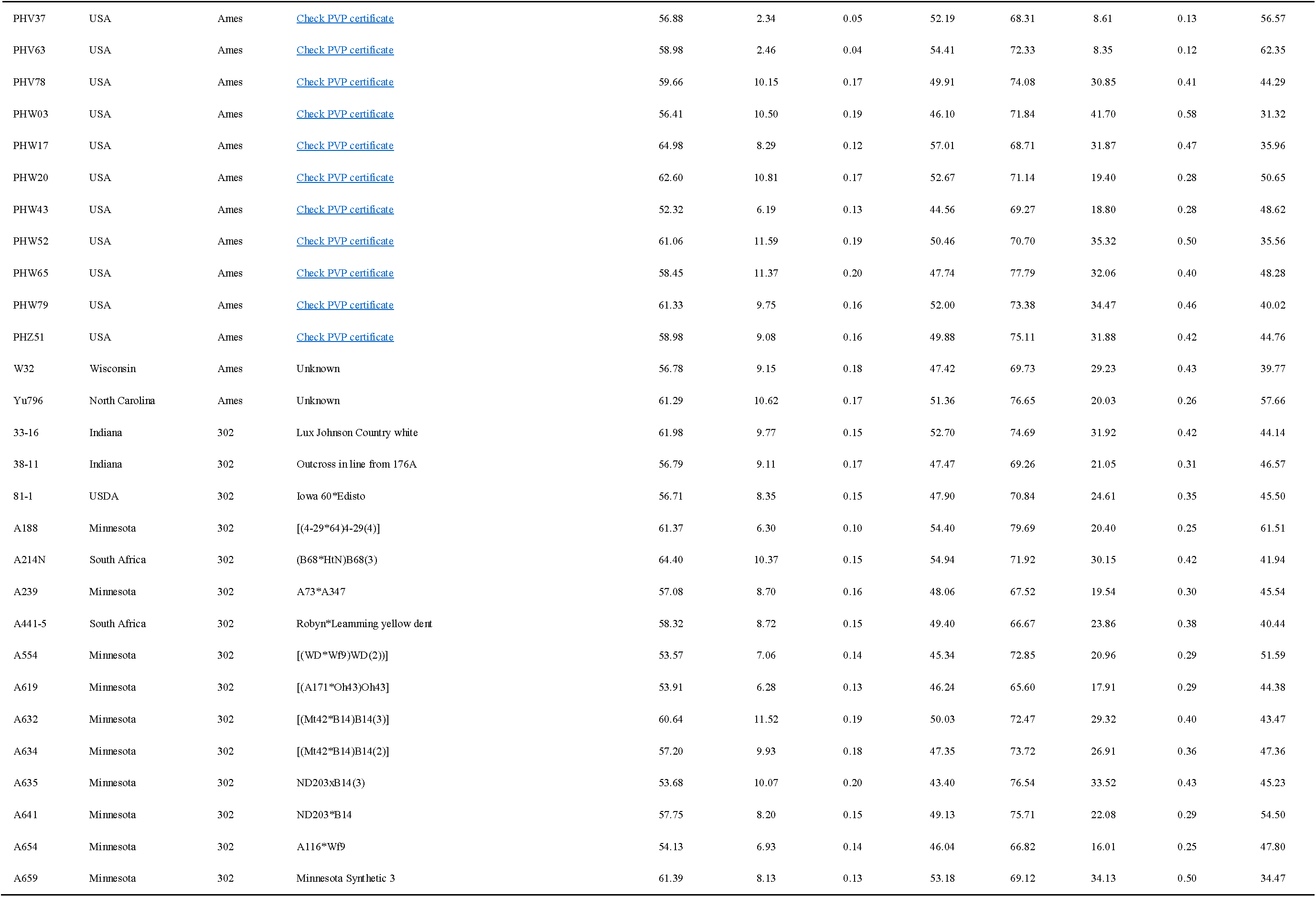
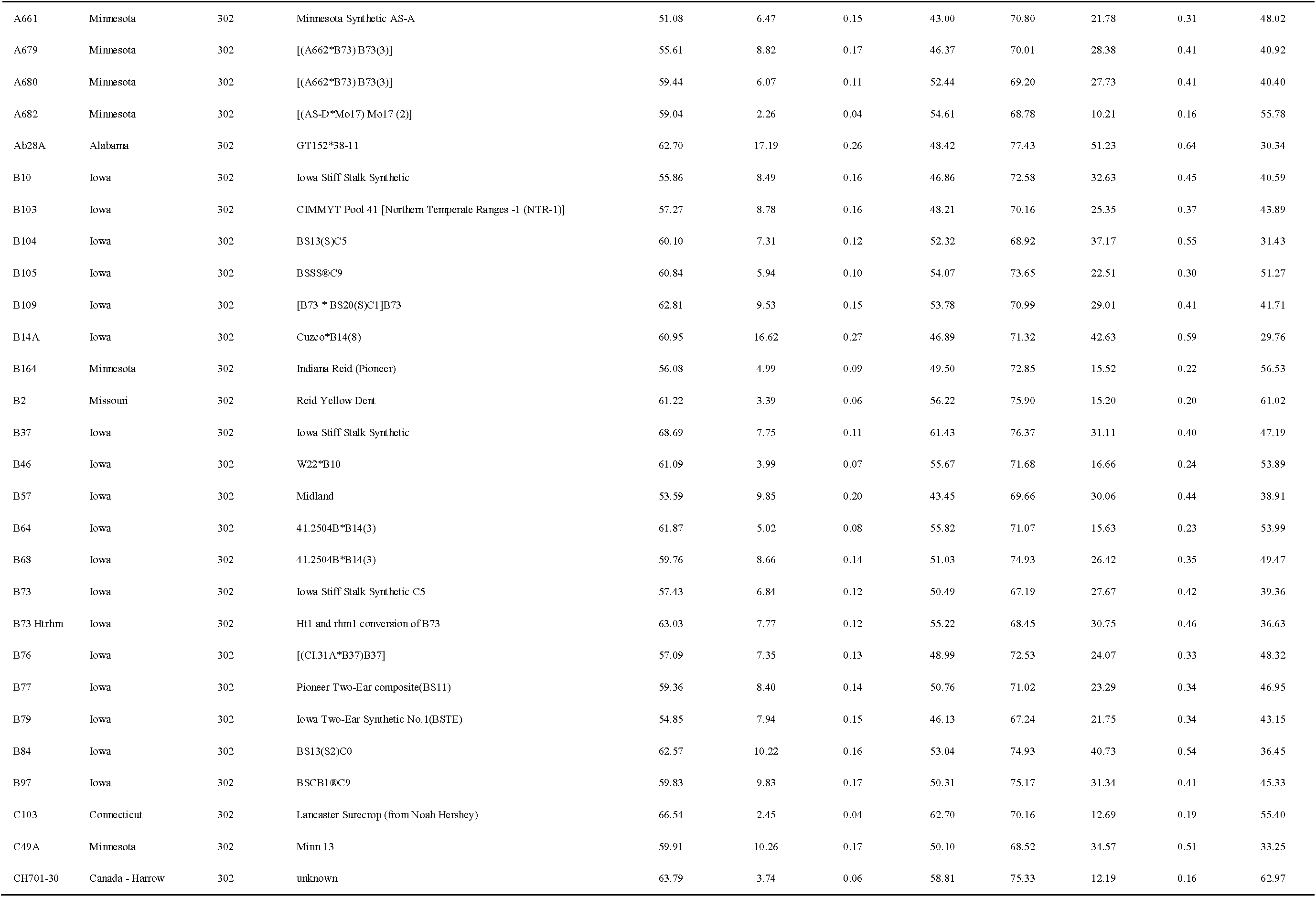
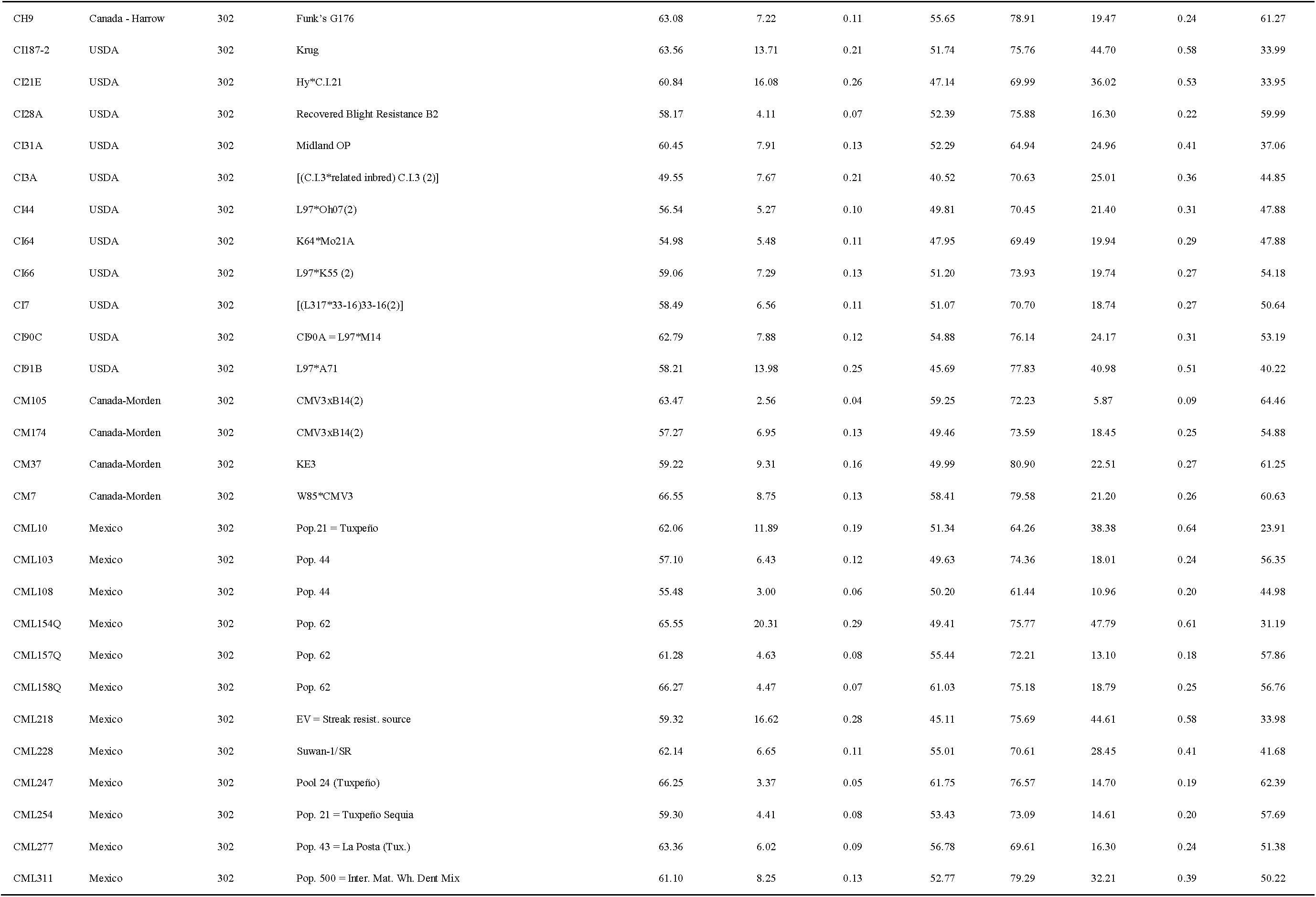
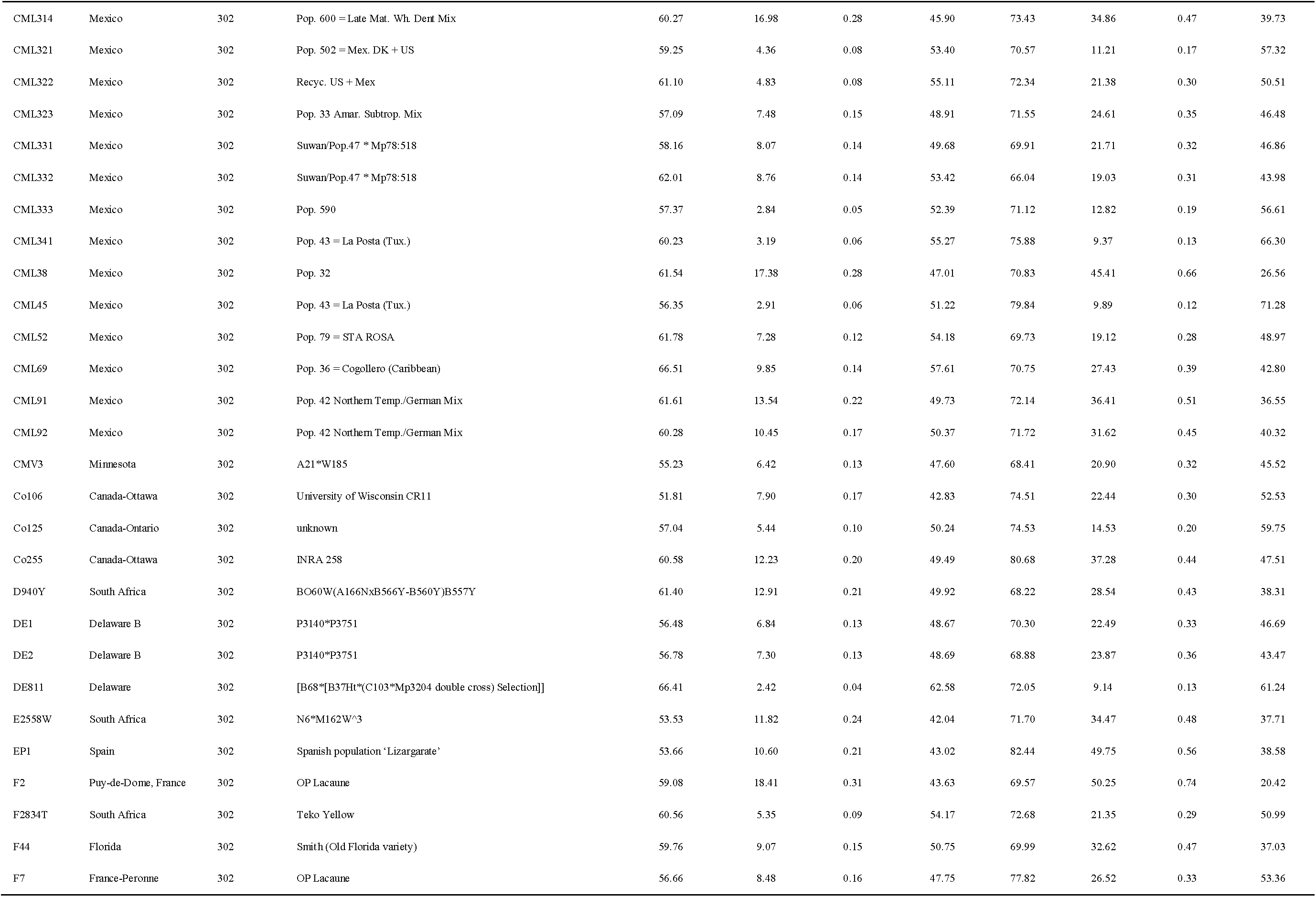
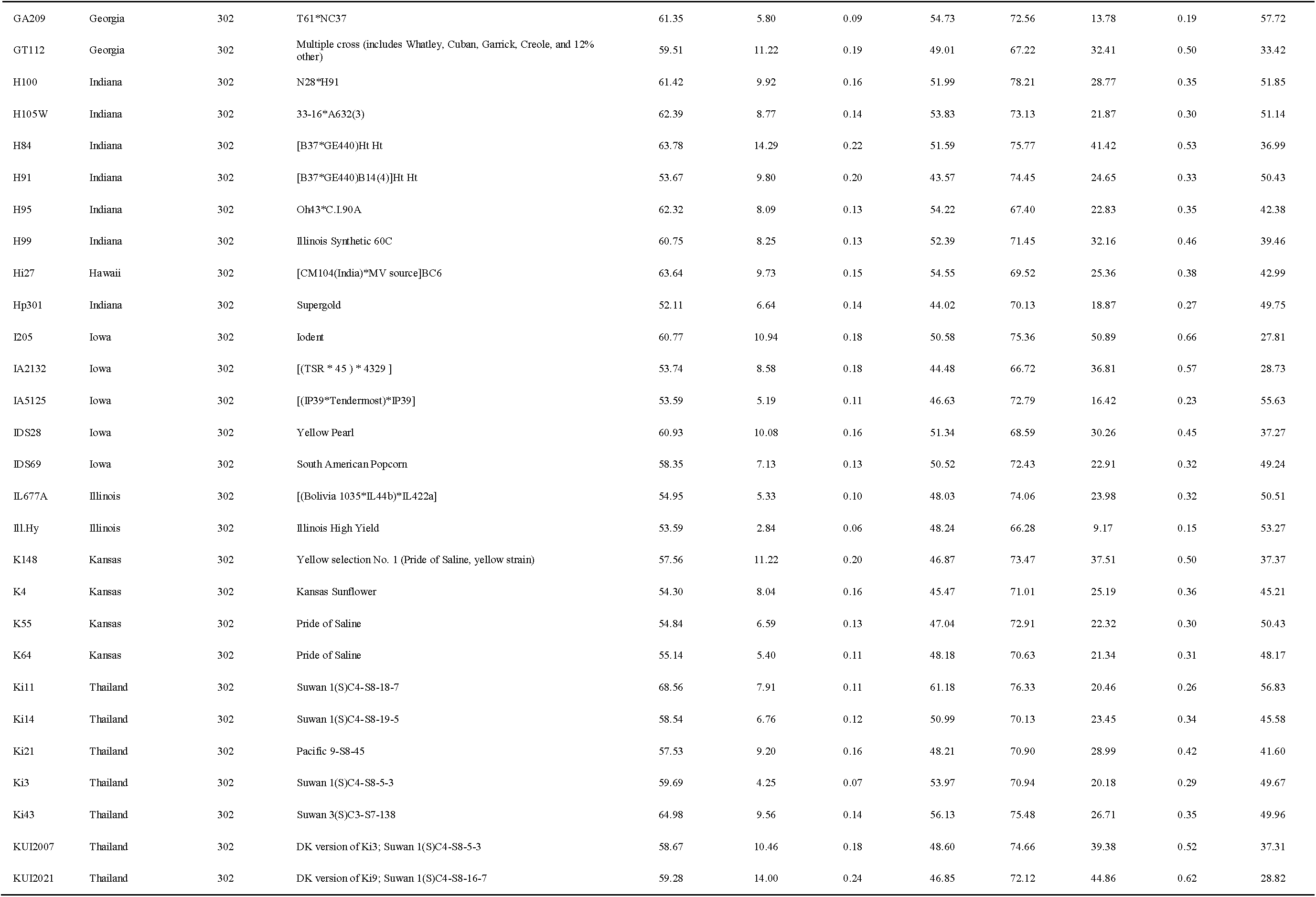
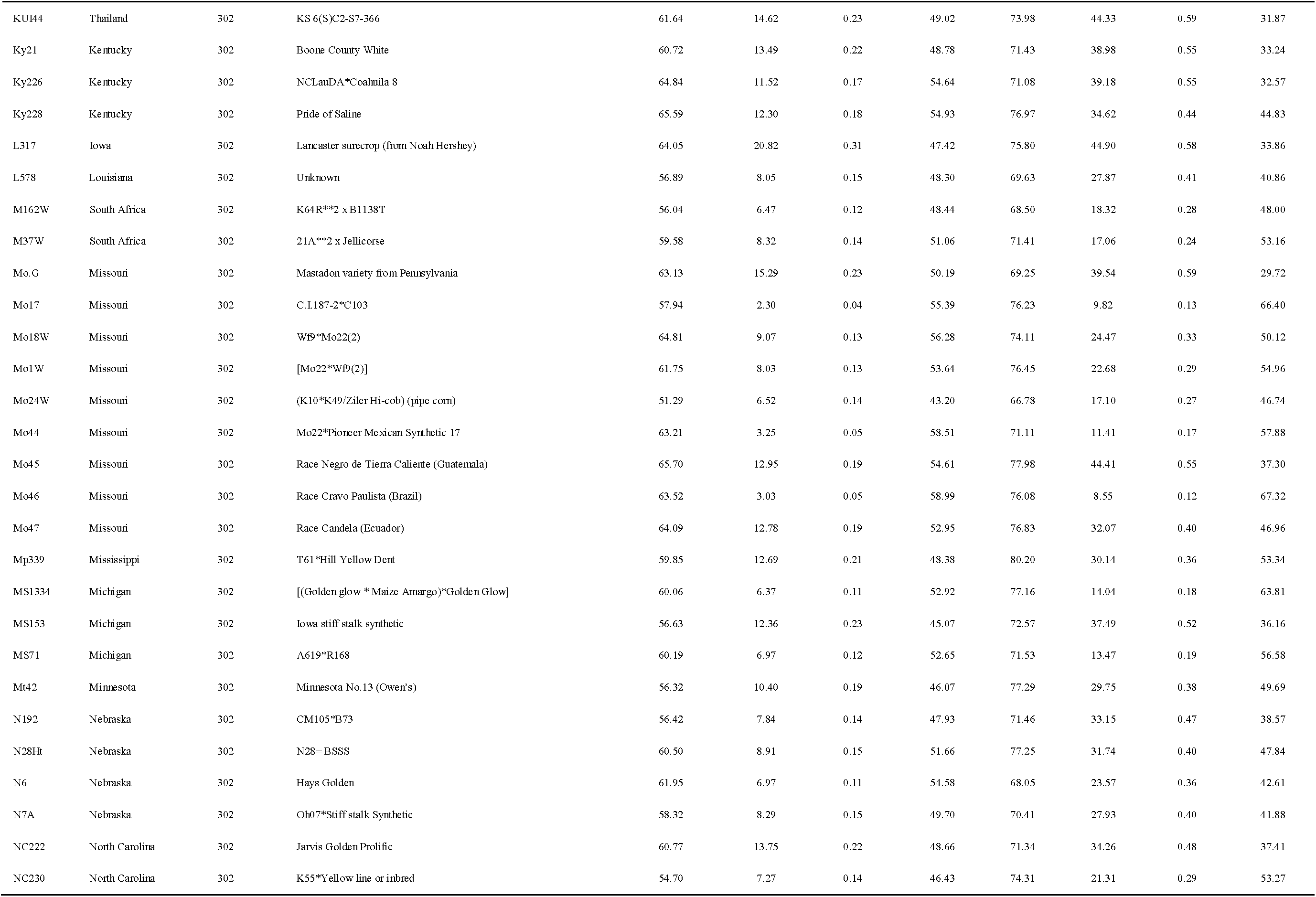
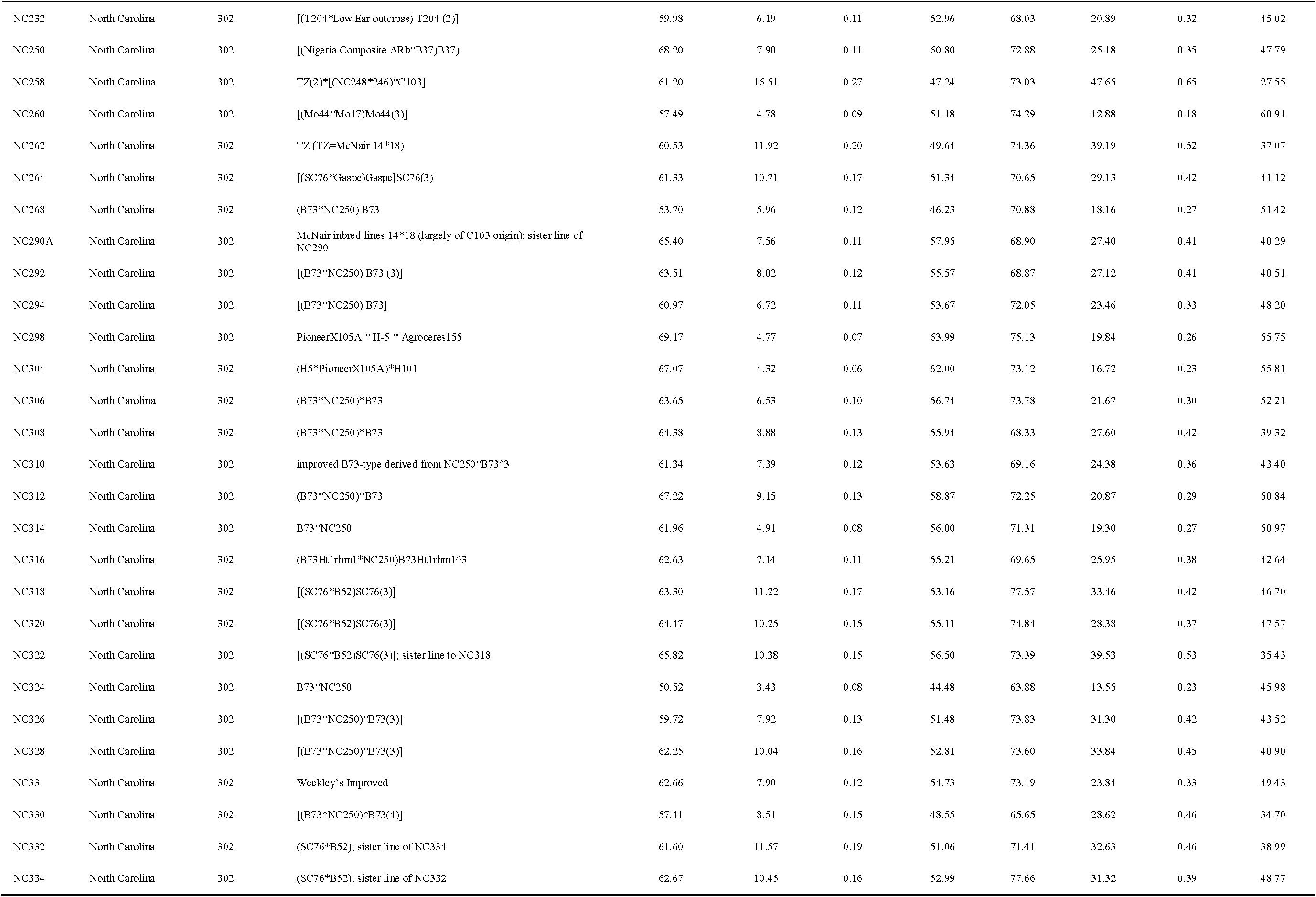
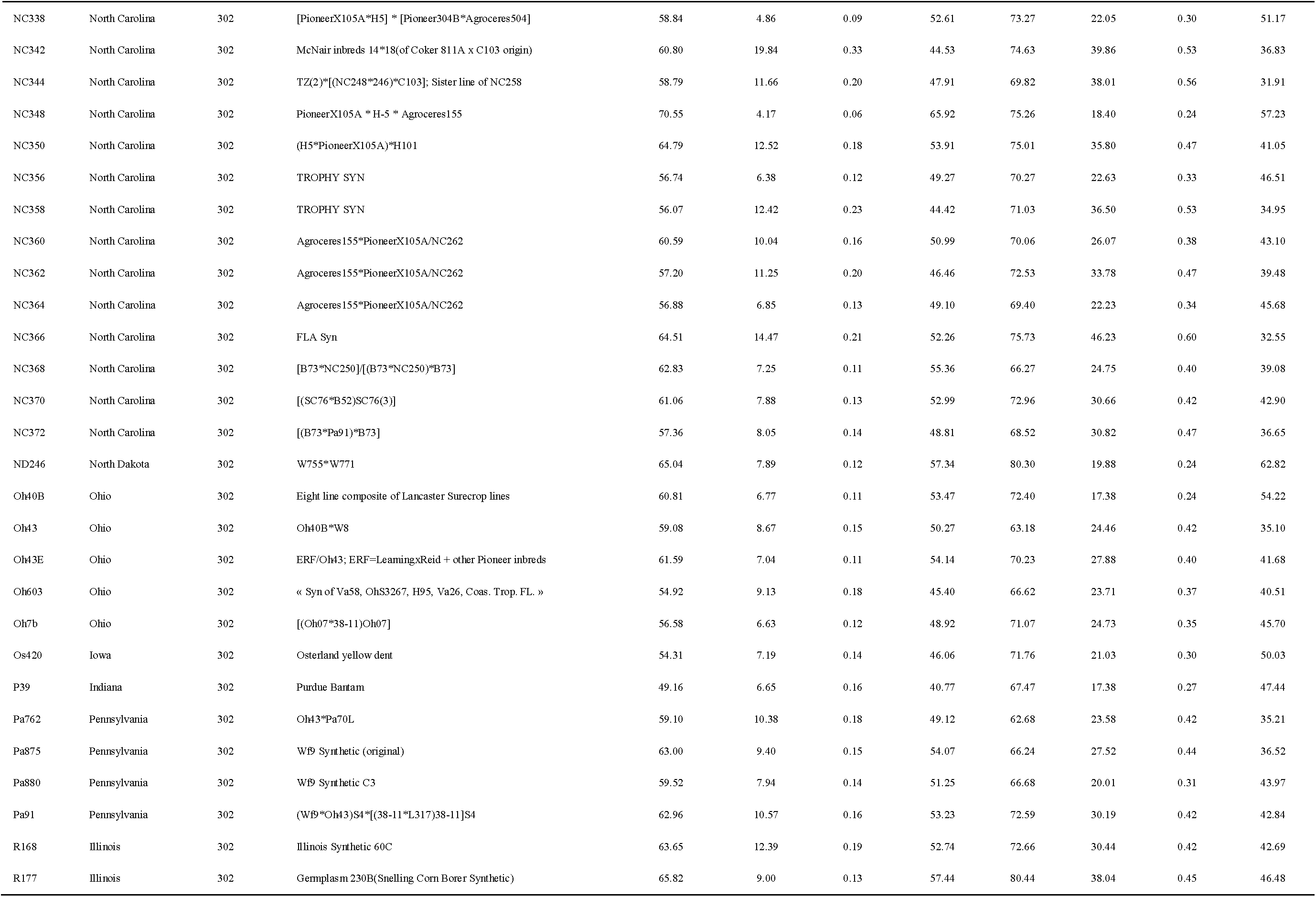
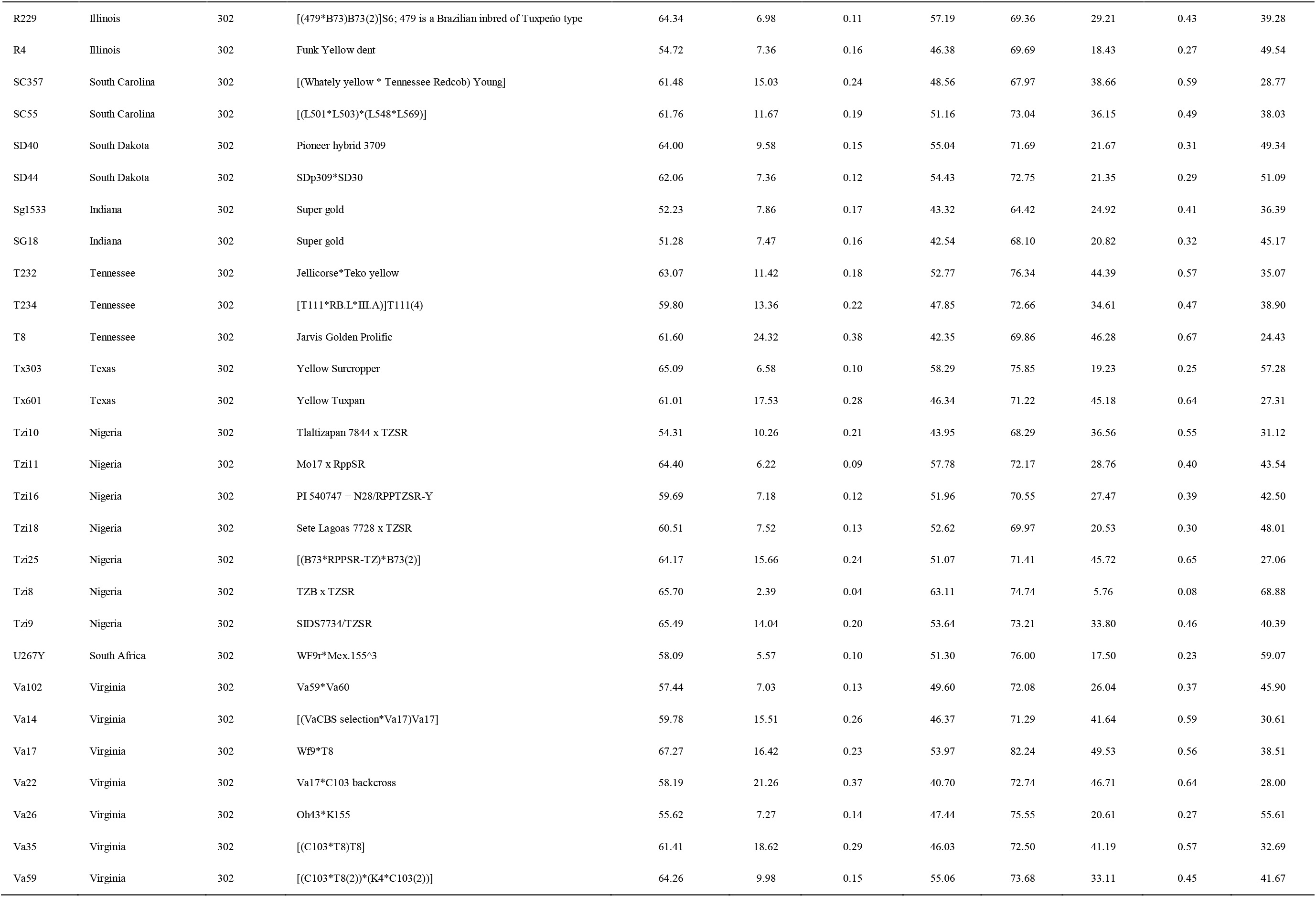
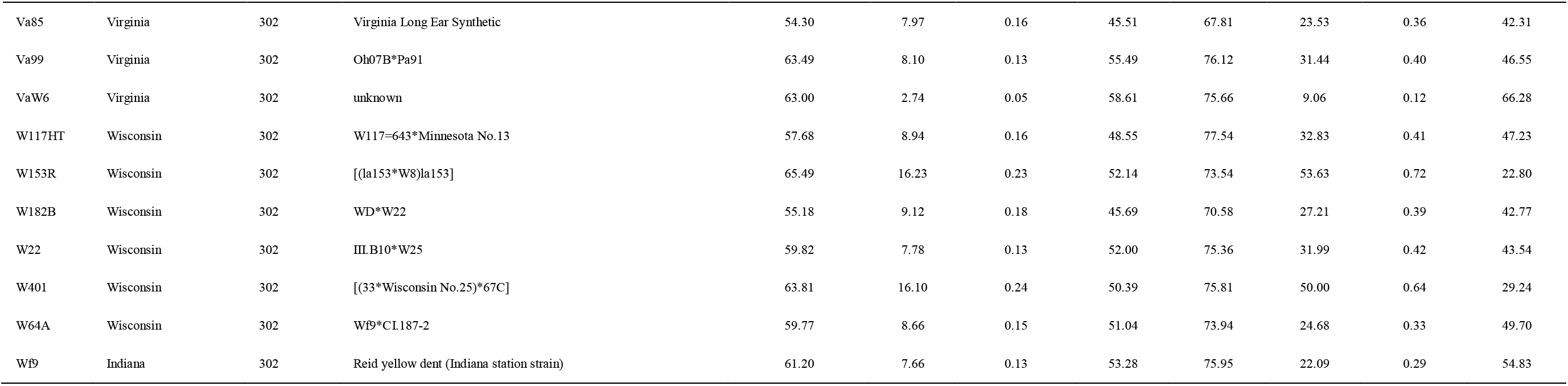
The BLUP values of the wild-type (WT) and mutant (MT) siblings of the F_1_ hybrids of *Oy1*-*N1989*/*oy1*:B73 with respective maize diversity lines (MDL). Information on the inbred lines from maize association panel (referred to as 302) was adapted from Flint-Garcia *et al.* 2005.

**Table S5.** The summary of the HapMap3 variants before and after filtering to remove SNPs with minor allele frequency < 0.05 (5%) and missing > 0.1 (10%).

| Chromosome | Sites_before_filtering | Sites_after_filtering |
| --- | --- | --- |
| 1 | 12550106 | 2900472 |
| 2 | 9620462 | 2247603 |
| 3 | 9415581 | 2208140 |
| 4 | 9862608 | 2146099 |
| 5 | 8226794 | 1855240 |
| 6 | 6502278 | 1430027 |
| 7 | 6902443 | 1632102 |
| 8 | 6619311 | 1510768 |
| 9 | 6112165 | 1528972 |
| 10 | 5875644 | 1327869 |
| **Total** | **81687392** | **18787292** |

**Table S6.**
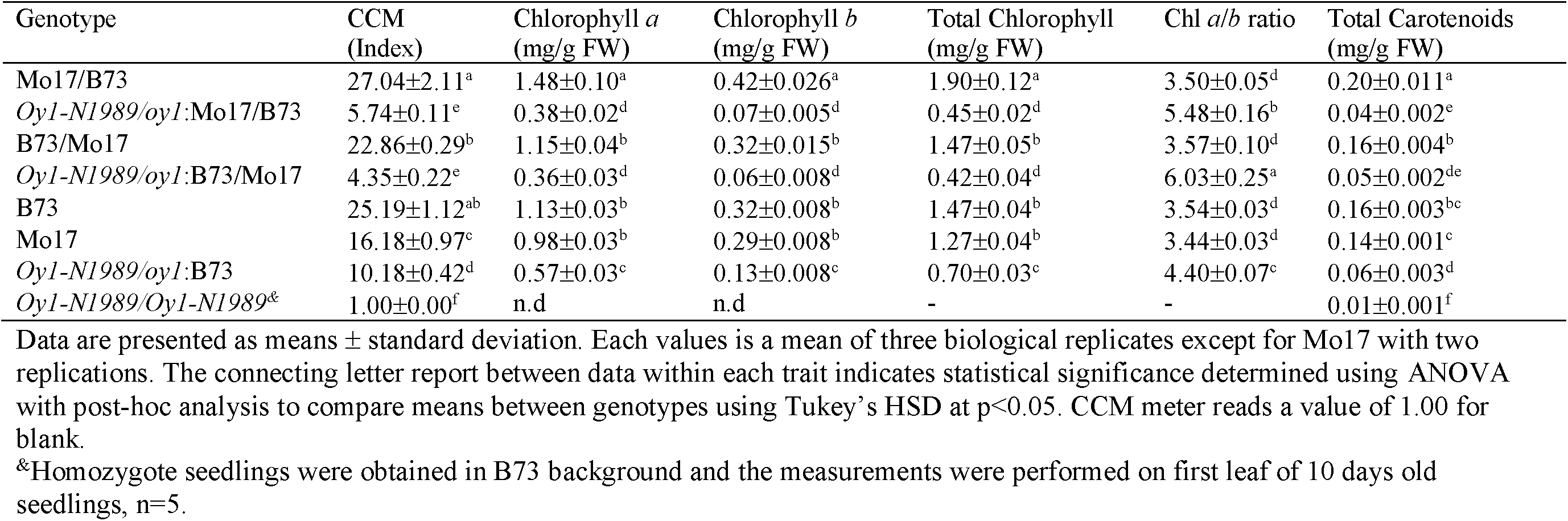
The chlorophyll accumulation in the third fully-expanded leaf at the V3 stage of greenhouse-grown maize seedlings.

**Table S7.** Means and standard deviation of pigment absorbance (index) from mutant (*Oy1-N1989/oy1*) and wild-type plants grown at the Purdue Agronomy Farm.

| Genotype | CCMI | CCMII |
| --- | --- | --- |
| B73 | 57.3±8.0<sup>a</sup> | 67.0±5.2<sup>a</sup> |
| Mo17/B73 | 57.8±7.3<sup>a</sup> | 76.4±5.6<sup>b</sup> |
| *Oy1-1989/oy1*:B73 | 6.8±1.3<sup>b</sup> | 27.7±3.2<sup>c</sup> |
| *Oy1-1989/oy1*:Mo17/B73 | 2.3±0.5<sup>c</sup> | 9.8±1.7<sup>d</sup> |

The connecting letter report indicates statistical significance determined using ANOVA followed by mean comparisons between the genotypes using Tukey's HSD at $p < 0.01$ . Check materials and methods for trait descriptions.

**Table S8.** The trait correlations among the CCM traits in IBM x *Oy1-N1989/oy1*:B73 F_1_ hybrid populations.

|  | WT_CCM I | MT_CCM I | Ratio_CCM I | Diff_CCM I | WT_CCM II | MT_CCM II | Ratio_CCM II | Diff_CCM II |
| --- | --- | --- | --- | --- | --- | --- | --- | --- |
| WT_CCM I |  | 0.069 | -0.222 | 0.901 | 0.368 | 0.062 | -0.045 | 0.235 |
| MT_CCM I | 0.312 |  | 0.948 | -0.371 | 0.059 | 0.875 | 0.859 | -0.616 |
| Ratio_CCM I | 0.001 | <.0001 |  | -0.619 | -0.062 | 0.829 | 0.855 | -0.673 |
| Diff_CCM I | <.0001 | <.0001 | <.0001 |  | 0.317 | -0.323 | -0.416 | 0.487 |
| WT_CCM II | <.0001 | 0.39 | 0.363 | <.0001 |  | 0.136 | -0.156 | 0.664 |
| MT_CCM II | 0.361 | <.0001 | <.0001 | <.0001 | 0.046 |  | 0.947 | -0.651 |
| Ratio_CCM II | 0.513 | <.0001 | <.0001 | <.0001 | 0.021 | <.0001 |  | -0.835 |
| Diff_CCM II | 0.001 | <.0001 | <.0001 | <.0001 | <.0001 | <.0001 | <.0001 |  |

The upper half contains Pearson's correlation coefficient and the lower half contains correlation p-values for each pairwise trait comparison. Self-comparisons (diagonal) are left blank.

**Table S9.**
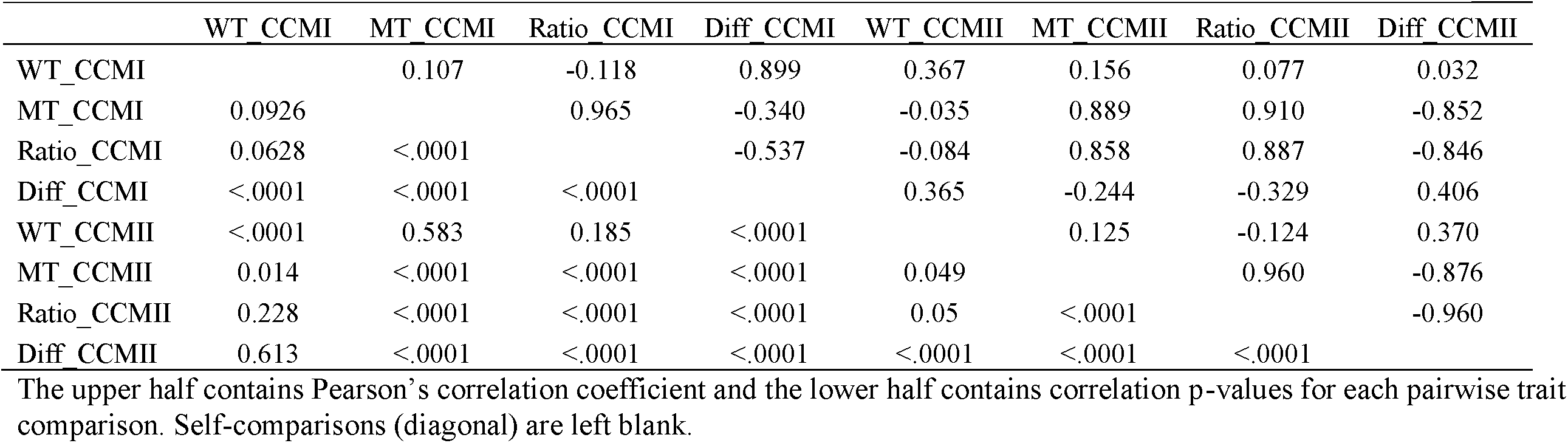
The trait Correlations among the CCM traits in Syn10 x *Oy1-N1989/oy1*:B73 F_1_ hybrid populations.

|  | WT_CCM I | MT_CCM I | Ratio_CCM I | Diff_CCM I | WT_CCM II | MT_CCM II | Ratio_CCM II | Diff_CCM II |
| --- | --- | --- | --- | --- | --- | --- | --- | --- |
| WT_CCM I |  | 0.107 | -0.118 | 0.899 | 0.367 | 0.156 | 0.077 | 0.032 |
| MT_CCM I | 0.0926 |  | 0.965 | -0.340 | -0.035 | 0.889 | 0.910 | -0.852 |
| Ratio_CCM I | 0.0628 | <.0001 |  | -0.537 | -0.084 | 0.858 | 0.887 | -0.846 |
| Diff_CCM I | <.0001 | <.0001 | <.0001 |  | 0.365 | -0.244 | -0.329 | 0.406 |
| WT_CCM II | <.0001 | 0.583 | 0.185 | <.0001 |  | 0.125 | -0.124 | 0.370 |
| MT_CCM II | 0.014 | <.0001 | <.0001 | <.0001 | 0.049 |  | 0.960 | -0.876 |
| Ratio_CCM II | 0.228 | <.0001 | <.0001 | <.0001 | 0.05 | <.0001 |  | -0.960 |
| Diff_CCM II | 0.613 | <.0001 | <.0001 | <.0001 | <.0001 | <.0001 | <.0001 |  |

The upper half contains Pearson's correlation coefficient and the lower half contains correlation p-values for each pairwise trait comparison. Self-comparisons (diagonal) are left blank.

**Table S10.** The summary of the QTL detected for CCM traits in IBM x *Oy1-N1989/oy1*:B73 F_1_ hybrid populations.

| Trait Identifier | QTL<sup>a</sup> | Chr<sup>b</sup> | LOD<sup>c</sup> | Position<sup>d</sup> | Left Marker | Right Marker | LOD-2 Interval<sup>e</sup> |  | PVE<sup>f</sup> | Mean±SE<sup>g</sup> |  | Greater Allele |
| --- | --- | --- | --- | --- | --- | --- | --- | --- | --- | --- | --- | --- |
|  |  |  |  |  |  |  | Left | Right |  | B73 | Mo17 |  |
| MT_CCM I | 1 | 10 | 46.9 | 139 | AI795367 | AY109994 | 137 | 141 | 53.68 | 8.81±0.188 | 4.16±0.226 | B73 |
| WT_CCM I | 1 | 1 | 3.6 | 523 | bnl5.59a | php20654 | 519 | 534 | 7.12 | 47.27±0.548 | 43.51±0.734 | B73 |
| MT_CCM II | 1 | 10 | 45.0 | 139 | AI795367 | AY109994 | 128 | 141 | 51.48 | 18.27±0.448 | 7.38±0.585 | B73 |
| WT_CCM II | 0 |  |  |  |  |  |  |  |  |  |  |  |
| Ratio_CCM I | 1 | 10 | 39.7 | 139 | AI795367 | AY109994 | 137 | 145 | 50.19 | 0.19±0.004 | 0.09±0.005 | B73 |
| Ratio_CCM II | 1 | 10 | 44.4 | 139 | AI795367 | AY109994 | 137 | 142 | 55.38 | 0.36±0.009 | 0.14±0.01 | B73 |
| Diff_CCM I | 1 | 10 | 6.6 | 145 | phi059 | isu85b | 126 | 153 | 13.18 | 36.89±0.576 | 42.24±0.724 | B73 |
| Diff_CCM II | 1 | 10 | 22.4 | 138 | AI795367 | AY109994 | 129 | 142 | 36.80 | 32.95±0.687 | 44.77±0.912 | B73 |

<sup>a</sup>Number of QTL detected for a given trait
<sup>b</sup>Chromosome location of a QTL
<sup>c</sup>LOD score at the peak of a given QTL
<sup>d</sup>Peak position and <sup>e</sup>LOD-2 interval of the QTL in terms of genetic position in centiMorgan(cM)
<sup>f</sup>PVE is percent of variance explained by the QTL at this position as estimated by regression and reported as an R<sup>2</sup> value\*100
<sup>g</sup>Mean and standard error of the trait with B73 and Mo17 genotype at the peak marker of the detected QTL

**Table S11.** The summary of the QTL detected from CCM traits in Syn10 x *Oy1-N1989/oy1*:B73 F_1_ hybrid populations.

| Trait | QTL<sup>a</sup> | Chr<sup>b</sup> | LOD<sup>c</sup> | Position<sup>d</sup> | Left Marker | Right Marker | LOD-2 Interval<sup>e</sup> |  | PVE<sup>f</sup> | Mean±SE<sup>g</sup> |  | Greater allele |
| --- | --- | --- | --- | --- | --- | --- | --- | --- | --- | --- | --- | --- |
|  |  |  |  |  |  |  | Left | Right |  | B73 | Mo17 |  |
| MT_CCM I | 1 | 10 | 59.9 | 189 | chr10.90.5 | chr10.93 | 189 | 193 | 65.2 | 7.00±0.117 | 2.65±0.174 | B73 |
| WT_CCM I | 0 |  |  |  |  |  |  |  |  |  |  |  |
| Ratio_CCM I | 1 | 10 | 54.0 | 189 | chr10.90.5 | chr10.93 | 189 | 192 | 62.1 | 0.14±0.002 | 0.05±0.003 | B73 |
| Diff_CCM I | 0 |  |  |  |  |  |  |  |  |  |  |  |
| MT_CCM II | 1 | 10 | 83.6 | 192 | chr10.94.5 | chr10.95.5 | 190 | 193 | 66.5 | 29.07±0.478 | 8.39±0.731 | B73 |
| WT_CCM II | 0 |  |  |  |  |  |  |  |  |  |  |  |
| Ratio_CCM II | 1 | 10 | 78.1 | 192 | chr10.94.5 | chr10.95.5 | 190 | 193 | 66.9 | 0.40±0.006 | 0.11±0.010 | B73 |
| Diff_CCM II | 1 | 10 | 52.1 | 189 | chr10.90.5 | chr10.93 | 189 | 191 | 60.3 | 42.46±0.623 | 63.39±0.926 | B73 |

<sup>a</sup>Number of QTL detected for a given trait
<sup>b</sup>Chromosome location of a QTL
<sup>c</sup>LOD score at the peak of a given QTL
<sup>d</sup>Peak position and <sup>e</sup>LOD-2 interval of the QTL in terms of genetic position in centiMorgan(cM)
<sup>f</sup>PVE is percent of variance explained by the QTL at this position as estimated by regression and reported as an R<sup>2</sup> value\*100
<sup>g</sup>Mean and standard error of the trait with B73 and Mo17 genotype at the peak marker of the detected QTL

**Table S12.** Recombinants within the *vey1* region derived from Syn10 x *Oy1-N1989/oy1*:B73 F_1_ populations.

| Line ID | MT_CCMI | Ratio_CCMI | MT_CCMII | Ratio_CCMII | chr10.90.5 | chr10.93 | chr10.94.5 | chr10.95.5 | Category<sup>1</sup> |
| --- | --- | --- | --- | --- | --- | --- | --- | --- | --- |
| IBM_31 | 7.0 | 0.13 | 33.2 | 0.46 | B | A | A | A | Suppressed |
| IBM_41 | 2.3 | 0.05 | 7.0 | 0.10 | B | B | A | A | Enhanced |
| IBM_149 | 1.7 | 0.03 | 5.7 | 0.09 | B | B | B | A | Enhanced |
| IBM_144 | 1.7 | 0.03 | 5.6 | 0.07 | B | B | B | A | Enhanced |
| IBM_199 | 8.9 | 0.20 | 30.1 | 0.41 | A | A | A | B | Suppressed |
| IBM_201 | 8.5 | 0.15 | 32.0 | 0.40 | A | A | A | B | Suppressed |
| IBM_214 | 1.8 | 0.03 | 5.5 | 0.07 | B | B | B | A | Enhanced |
| IBM_217 | 2.4 | 0.04 | 7.3 | 0.10 | B | B | B | A | Enhanced |
| IBM_229 | 9.7 | 0.17 | 34.8 | 0.49 | B | A | A | A | Suppressed |

The genotype code 'A' and 'B' for each marker denote B73 and Mo17 genotype, respectively.
<sup>1</sup>Severity of the mutation. Check the respective CCM trait value for quantitative assessment of severity.

**Table S13.** The trait correlations of various CCM traits using mean values of MDL x *Oy1-N1989/oy1*:B73 F_1_ hybrid populations.

|  | WT CCMI | MT CCMI | Ratio CCMI | Diff CCMI | WT CCMII | MT CCMII | Ratio CCMII | Diff CCMII |
| --- | --- | --- | --- | --- | --- | --- | --- | --- |
| WT_CCMI |  | 0.182 | -0.047 | 0.825 | 0.329 | 0.178 | 0.124 | 0.007 |
| MT_CCMI | 0.001 |  | 0.964 | -0.405 | 0.135 | 0.861 | 0.846 | -0.750 |
| Ratio_CCMI | 0.410 | <.0001 |  | -0.596 | 0.080 | 0.823 | 0.818 | -0.746 |
| Diff_CCMI | <.0001 | <.0001 | <.0001 |  | 0.231 | -0.329 | -0.372 | 0.438 |
| WT_CCMII | <.0001 | 0.012 | 0.142 | <.0001 |  | 0.171 | -0.028 | 0.351 |
| MT_CCMII | 0.002 | <.0001 | <.0001 | <.0001 | 0.002 |  | 0.976 | -0.862 |
| Ratio_CCMII | 0.030 | <.0001 | <.0001 | <.0001 | 0.601 | <.0001 |  | -0.942 |
| Diff_CCMII | 0.904 | <.0001 | <.0001 | <.0001 | <.0001 | <.0001 | <.0001 |  |

The upper half contains Pearson's correlation coefficient and the lower half contains correlation p-values for each pairwise trait comparison. Self-comparisons (diagonal) are left blank.

**Table S14.** The broad sense heritability and variance estimates of CCM traits measured in MDL x *Oy1-N1989/oy1*:B73 F_1_ hybrid populations.

| Trait | Heritability | LSD | Variance |
| --- | --- | --- | --- |
| WT_CCM I | 0.59 | 14.56 | 80.64 |
| MT_CCM I | 0.95 | 3.22 | 18.77 |
| Ratio_CCM I | 0.92 | 0.06 | 0.01 |
| Diff_CCM I | 0.65 | 14.53 | 88.81 |
| WT_CCM II | 0.63 | 11.74 | 57.20 |
| MT_CCM II | 0.96 | 7.36 | 126.66 |
| Ratio_CCM II | 0.94 | 0.12 | 0.02 |
| Diff_CCM II | 0.87 | 13.37 | 169.53 |

**Table S15.**
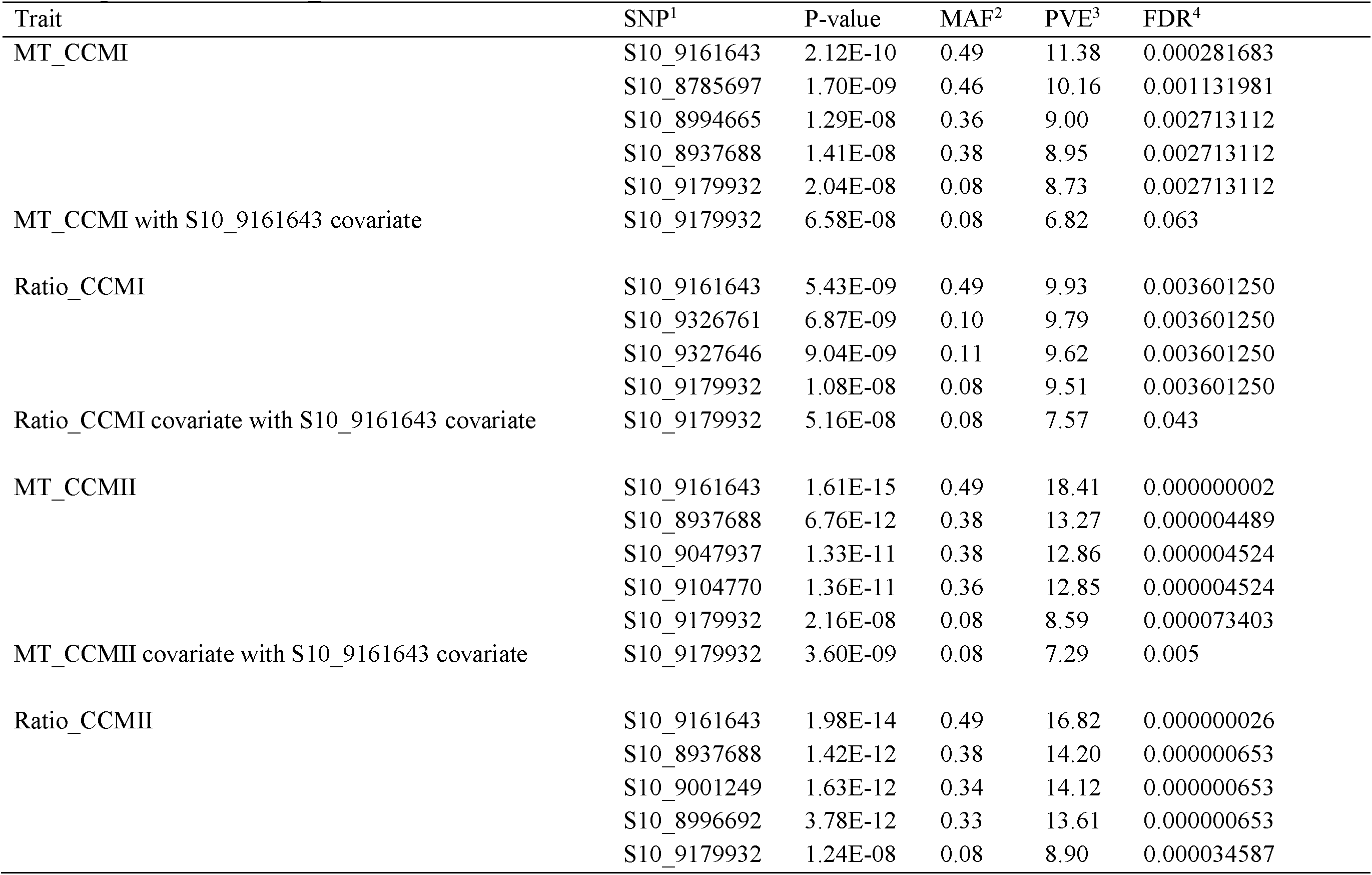
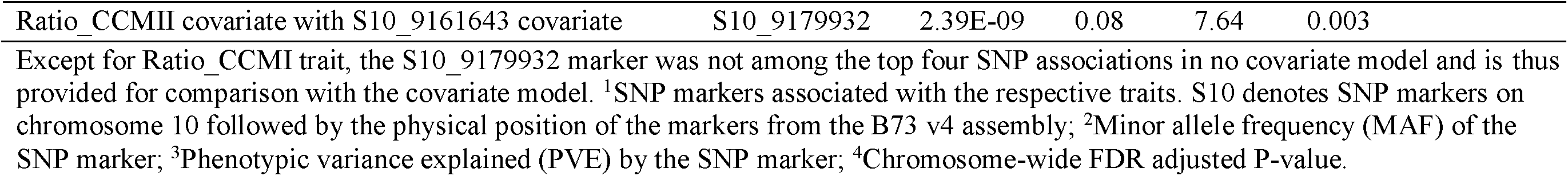
The summary of the top four statistically significant SNP markers associated with CCM traits by GWAS and top SNP following the addition of S10_9161643 as a covariate for each trait.

| Trait | SNP<sup>1</sup> | P-value | MAF<sup>2</sup> | PVE<sup>3</sup> | FDR<sup>4</sup> |
| --- | --- | --- | --- | --- | --- |
| MT_CCMI | S10_9161643 | 2.12E-10 | 0.49 | 11.38 | 0.000281683 |
|  | S10_8785697 | 1.70E-09 | 0.46 | 10.16 | 0.001131981 |
|  | S10_8994665 | 1.29E-08 | 0.36 | 9.00 | 0.002713112 |
|  | S10_8937688 | 1.41E-08 | 0.38 | 8.95 | 0.002713112 |
|  | S10_9179932 | 2.04E-08 | 0.08 | 8.73 | 0.002713112 |
| MT_CCMI with S10_9161643 covariate | S10_9179932 | 6.58E-08 | 0.08 | 6.82 | 0.063 |
| Ratio_CCMI | S10_9161643 | 5.43E-09 | 0.49 | 9.93 | 0.003601250 |
|  | S10_9326761 | 6.87E-09 | 0.10 | 9.79 | 0.003601250 |
|  | S10_9327646 | 9.04E-09 | 0.11 | 9.62 | 0.003601250 |
|  | S10_9179932 | 1.08E-08 | 0.08 | 9.51 | 0.003601250 |
| Ratio_CCMI covariate with S10_9161643 covariate | S10_9179932 | 5.16E-08 | 0.08 | 7.57 | 0.043 |
| MT_CCMII | S10_9161643 | 1.61E-15 | 0.49 | 18.41 | 0.000000002 |
|  | S10_8937688 | 6.76E-12 | 0.38 | 13.27 | 0.000004489 |
|  | S10_9047937 | 1.33E-11 | 0.38 | 12.86 | 0.000004524 |
|  | S10_9104770 | 1.36E-11 | 0.36 | 12.85 | 0.000004524 |
|  | S10_9179932 | 2.16E-08 | 0.08 | 8.59 | 0.000073403 |
| MT_CCMII covariate with S10_9161643 covariate | S10_9179932 | 3.60E-09 | 0.08 | 7.29 | 0.005 |
| Ratio_CCMII | S10_9161643 | 1.98E-14 | 0.49 | 16.82 | 0.000000026 |
|  | S10_8937688 | 1.42E-12 | 0.38 | 14.20 | 0.000000653 |
|  | S10_9001249 | 1.63E-12 | 0.34 | 14.12 | 0.000000653 |
|  | S10_8996692 | 3.78E-12 | 0.33 | 13.61 | 0.000000653 |
|  | S10_9179932 | 1.24E-08 | 0.08 | 8.90 | 0.000034587 |
| Ratio_CCMII covariate with S10_9161643 covariate | S10_9179932 | 2.39E-09 | 0.08 | 7.64 | 0.003 |

Except for Ratio\_CCMI trait, the S10\_9179932 marker was not among the top four SNP associations in no covariate model and is thus provided for comparison with the covariate model. <sup>1</sup>SNP markers associated with the respective traits. S10 denotes SNP markers on chromosome 10 followed by the physical position of the markers from the B73 v4 assembly; <sup>2</sup>Minor allele frequency (MAF) of the SNP marker; <sup>3</sup>Phenotypic variance explained (PVE) by the SNP marker; <sup>4</sup>Chromosome-wide FDR adjusted P-value.

**Table S16.** Haplotypes at two SNPs at *vey1* locus associated with *Oy1*-*N1989* suppression and its effect on CCM traits in MDL x *Oy1*-*N1989*/*oy1*:B73 F_1_ populations.

| Haplotype<sup>&</sup> | Sample (n) | WT_CCMI | MT_CCMI | Ratio_CCMI | WT_CCMII | MT_CCMII | Ratio_CCMII |
| --- | --- | --- | --- | --- | --- | --- | --- |
| AG | 153 | 60.86±3.69<sup>a</sup> | 10.00±4.13<sup>a</sup> | 0.16±0.06<sup>a</sup> | 72.57±3.45 | 30.99±9.19<sup>a</sup> | 0.43±0.12<sup>a</sup> |
| CG | 121 | 59.00±4.21<sup>b</sup> | 7.65±2.42<sup>b</sup> | 0.13±0.04<sup>b</sup> | 72.04±3.97 | 22.65±6.69<sup>b</sup> | 0.32±0.09<sup>b</sup> |
| AA | 8 | 58.56±2.43<sup>ab</sup> | 3.93±1.17<sup>c</sup> | 0.07±0.02<sup>c</sup> | 73.78±3.74 | 15.28±4.90<sup>c</sup> | 0.21±0.07<sup>c</sup> |
| CA | 23 | 60.53±4.45<sup>ab</sup> | 2.78±0.99<sup>c</sup> | 0.05±0.02<sup>c</sup> | 72.99±3.08 | 10.63±2.82<sup>c</sup> | 0.15±0.03<sup>c</sup> |

<sup>&</sup>The first nucleotide of the haplotype denotes variant (A/C) at SNP S10\_9161643 and the second nucleotide of the haplotype denotes variant (G/A) at SNP S10\_9179932. The physical positions of these SNPs are from the B73 v4 assembly.

**Table S17.** The chlorophyll quantification of plants segregating for the allelic interaction between *Oy1-N1989* and *oy1-yg* alleles at *oy1*.

| Cross | Days after planting | *Oyl-N1989/oyl-yg* | *Oyl-N1989/+* | *oyl-yg/+* | *oyl-yg/oyl-yg* |
| --- | --- | --- | --- | --- | --- |
| (Mo17 x *oyl-yg/oyl-yg*) x *Oyl-N1989/+*:B73 | 21 | 1.7±0.4<sup>a</sup> | 3.0±0.6<sup>a</sup> | . | . |
| (B73 x *oyl-yg/oyl-yg*) x *Oyl-N1989/+*:B73 | 21 | 2.9±0.5<sup>b</sup> | 5.1±0.8<sup>b</sup> | . | . |
| (Mo17 x *oyl-yg/oyl-yg*) x *Oyl-N1989/+*:B73 | 40 | 3.7±0.6<sup>c</sup> | 7.0±0.3<sup>c</sup> | . | . |
| (B73 x *oyl-yg/oyl-yg*) x *Oyl-N1989/+*:B73 | 40 | 7.5±0.9<sup>d</sup> | 17.1±3.3<sup>d</sup> | . | . |
| (Mo17 x *oyl-yg/oyl-yg*) x *oyl-yg/oyl-yg* | 21 | . | . | 27.6±5.7<sup>a</sup> | 7.6±2.8<sup>a</sup> |
| (B73 x *oyl-yg/oyl-yg*) x *oyl-yg/oyl-yg* | 21 | . | . | 27.8±6.8<sup>a</sup> | 8.1±2.8<sup>a</sup> |
| (Mo17 x *oyl-yg/oyl-yg*) x *oyl-yg/oyl-yg* | 40 | . | . | 32.2±4.9<sup>b</sup> | 16.7±5<sup>b</sup> |
| (B73 x *oyl-yg/oyl-yg*) x *oyl-yg/oyl-yg* | 40 | . | . | 38.7±6.1<sup>b</sup> | 21±6.2<sup>b</sup> |

Data are presented as means and standard deviations. The sample size in each category varies from 5-20 plants. Comparisons to declare statistical significance were done only between the crosses with B73 and Mo17 as a parent within each genotype class of the progenies quantified on the same day (i.e. within 21 or 40 days after planting). The connecting letter report between the two samples indicates the statistical significance at $p < 0.05$ using student's t-test.

**Table S18.** The summary of the average CCM value of the F_1_ hybrids of inbred lines crossed with *Oy1-N1989/oy1*:B73, and allelic state at the 6 bp (two amino acids) indel in the coding sequence of OY1 transcript in the respective parental inbred line.

| Inbred | Insertion<sup>a</sup> | MT CCM I | WT CCM I | Ratio CCM I | MT CCM II | WT CCM II | Ratio CCM II | Category<sup>b</sup> |
| --- | --- | --- | --- | --- | --- | --- | --- | --- |
| NC350 | AT | 12.7 | 68.1 | 0.19 | 36.2 | 76.6 | 0.48 | sup |
| NC358 | AT | 12.6 | 53.3 | 0.24 | 36.9 | 70.3 | 0.54 | sup |
| CML69 | AT | 9.9 | 71.0 | 0.14 | 27.5 | 69.9 | 0.39 | sup |
| B97 | AS | 9.9 | 59.7 | 0.17 | 31.5 | 76.9 | 0.41 | sup |
| Mo18W | AS | 9.1 | 68.1 | 0.13 | 24.4 | 75.2 | 0.33 | sup |
| Oh43 | AS | 8.7 | 58.4 | 0.15 | 24.3 | 57.9 | 0.42 | sup |
| W22 | AT | 8.5 | 64.4 | 0.13 | 37.8 | 74.6 | 0.51 | sup |
| IL14H | AT | 7.7 | 50.3 | 0.15 | 22.5 | 68.8 | 0.33 | sup |
| B73 | - | 7.0 | 57.2 | 0.12 | 26.3 | 69.4 | 0.38 | sup |
| MS71 | AS | 6.9 | 60.3 | 0.11 | 12.9 | 71.1 | 0.18 | sup |
| P39 | AT | 6.5 | 41.6 | 0.16 | 16.9 | 64.7 | 0.26 | sup |
| Oh7b | AS | 6.5 | 54.2 | 0.12 | 24.6 | 70.4 | 0.35 | sup |
| CML103 | - | 6.3 | 55.1 | 0.12 | 17.6 | 75.6 | 0.23 | sup |
| CML322 | - | 4.6 | 61.9 | 0.07 | 21.1 | 72.4 | 0.29 | enh |
| Ki3 | AT | 4.0 | 59.5 | 0.07 | 19.9 | 70.2 | 0.28 | enh |
| CML247 | AT | 3.1 | 70.6 | 0.04 | 14.1 | 79.1 | 0.18 | enh |
| Tzi8 | AT | 2.4 | 71.0 | 0.03 | 5.4 | 78.8 | 0.07 | enh |
| Mo17 | AS | 2.0 | 53.7 | 0.04 | 9.6 | 76.5 | 0.13 | enh |
| **Linear Fit (R<sup>2</sup>)<sup>c</sup>** |  | **0.009** | **0.00003** | **0.005** | **0.002** | **0.008** | **0.0009** |  |
| **P-value<sup>d</sup>** |  | **0.49** | **0.66** | **0.56** | **0.56** | **0.87** | **0.64** |  |

Data were derived from three replications planted in a RCBD.
<sup>a</sup>Coding sequence polymorphism in the third exon of OY1 protein with three alternate alleles. Abbreviated symbols are A:Alanine, T:Threonine, S:Serine, -:Deletion of both amino acids.
<sup>b</sup>Category of the genotypes in terms of the severity of *Oy1-N1989/oy1*:B73/inbred F<sub>1</sub> mutant individuals assigned based on CCM I and Ratio\_CCM I trait value. Abbreviated symbols are sup:suppressed mutants, enh:enhanced/severe mutants.
<sup>c</sup>R<sup>2</sup> of the linear regression model using Indel polymorphism as an explanatory variable onto a given trait as a response variable.
<sup>d</sup>P-value of the analysis of variance for effect of Indel polymorphism on the respective trait value.

**Table S19.** The linear regression of the top *vey1* linked marker (isu085b) and CCM traits from wild-type and mutant siblings of IBM x *Oy1-N1989/oy1*:B73 F_1_ populations on to OY1 expression (RPKM values) of the respective IBM line (n=74).

| Trait/Marker | R<sup>2</sup> (%) | P-value |
| --- | --- | --- |
| isu085b | 19.2 | <0.0001 |
| MT_CCM I | 25.2 | <0.0001 |
| WT_CCM I | 4.8 | 0.06 |
| MT_CCM II | 27.3 | <0.0001 |
| WT_CCM II | 0.9 | 0.40 |
| Ratio_CCM I | 31.3 | <0.0001 |
| Ratio_CCM II | 22.5 | <0.0001 |

**Table S20.** The allele expression bias at *oy1* in leaf tissue from the top fully-expanded leaf at the V3 stage.

| Genotype | SNP_252<sup>1</sup> | SNP252_Ref<sup>2</sup> | SNP_252_Alt<sup>3</sup> | SNP_317 | SNP317_Ref | SNP317_Alt | Ratio_SNP252<sup>ψ</sup> | Ratio_SNP317<sup>+</sup> | Average<sup>%</sup> |
| --- | --- | --- | --- | --- | --- | --- | --- | --- | --- |
| *oyl/oyl*:B/M<sup>&</sup> | . | . | . | . | . | . | . | . | 1.19±0.07 |
| *oyl/oyl*:B/M | C/C | . | . | C/T | 2635±51.60 | 2430±67.28 | . | 1.08±0.01<sup>a</sup> | 1.08±0.01<sup>a</sup> |
| *Oyl-N1989/oyl*:M/B | C/T | 2044±149.77 | 2291±152.87 | C/T | 2782±120.79 | 2522±67.29 | 1.12±0.01<sup>a</sup> | 1.10±0.02<sup>a</sup> | 1.11±0.01<sup>a</sup> |
| *Oyl-N1989/oyl*:B/M | C/T | 2138±142.51 | 2456±122.46 | C/T | 2677.33±99.71 | 2425±95.50 | 1.15±0.03<sup>a</sup> | 1.10±0.01<sup>a</sup> | 1.13±0.02<sup>a</sup> |
| *Oyl-N1989/oyl*:B | C/T | 2261±109.33 | 2290±76.62 | C/C | . | . | 1.01±0.02<sup>b</sup> | . | 1.01±0.02<sup>b</sup> |

The connecting letter report in all columns indicate statistical significance calculated using ANOVA with post-hoc analysis using Tukey's HSD with p<0.01.
<sup>1</sup>SNP position 252 with two alternate alleles. C corresponds to the wild-type/reference allele and T corresponds to the mutant/alternate allele. Same applies to SNP\_317 column, except this SNP is polymorphic only between B73 and Mo17; *Oyl-N1989* allele is monomorphic with B73 at SNP\_317.
<sup>2</sup>Mean ± standard deviation of allele count for the reference allele using three biological replications. Same applies to SNP317\_Ref column. Same applies to SNP317\_Ref column.
<sup>3</sup>Mean ± standard deviation of allele count for the alternate allele using three biological replications. Same applies to SNP317\_Alt column. Same applies to SNP317\_Alt column.
<sup>&</sup>Data obtained from Waters *et al.* 2017. Allele bias at *oyl* locus for plants grown under control condition.
<sup>ψ</sup>Mean ± standard deviation of the ratios of the read count from reference allele to the alternate allele at SNP252.
<sup>+</sup>Mean ± standard deviation of the ratios of the read count from reference allele to the alternate allele at SNP317.
<sup>%</sup>Mean ± standard deviation of the average of the ratios at SNP position 252 and 317.

**Table S21.** The chlorophyll approximation (using CCM) from the middle of the third leaf at the V3 stage on greenhouse-grown maize seedlings from a cross of B73, Mo17, and PH207 inbred lines (ear-parents) with *Oy1-N1989/oy1*:B73 plants (pollen-parent).

| Pedigree | Genotype | Sample size | CCM (Index) |
| --- | --- | --- | --- |
| B73 | wild-type | 8 | 21.16±2.65<sup>a</sup> |
| B73 | mutant | 5 | 7.95±0.45<sup>b</sup> |
| PH207/B73 | wild-type | 5 | 28.20±1.57<sup>c</sup> |
| PH207/B73 | mutant | 6 | 5.53±0.23<sup>d</sup> |
| Mo17/B73 | wild-type | 5 | 15.62±1.83<sup>e</sup> |
| Mo17/B73 | mutant | 5 | 3.90±0.12<sup>d</sup> |

Data are presented as mean ± standard deviation. The connecting letter report indicates statistical significance calculated using ANOVA with post-hoc analysis using Tukey's HSD with $p < 0.05$ .

**Table S22.** The distribution of normalized OY1 expression in the emerging shoot tissue of maize diversity lines (Kremling *et al.* 2018) at two SNPs associated with suppression of *Oy1*-*N1989* phenotype in MDL x *Oy1*-*N1989*/*oy1*:B73 F_1_ populations.

| Variant | Allele/Haplotype | Sample Size | OY1 (count) | R<sup>2</sup> (%)<sup>a</sup> |
| --- | --- | --- | --- | --- |
| S10_9161643 | A | 138 | 3.94 ± 0.24* | 7.14 |
|  | C | 119 | 3.79 ± 0.30 |  |
| S10_9179932 | A | 15 | 3.62 ± 0.20* | 4.80 |
|  | G | 242 | 3.89 ± 0.28 |  |
| Haplotypes | AG | 132 | 3.95 ± 0.24<sup>a</sup> | 11.24 |
|  | CG | 110 | 3.81 ± 0.30<sup>b</sup> |  |
|  | AA | 6 | 3.68 ± 0.17<sup>bc</sup> |  |
|  | CA | 9 | 3.58 ± 0.23<sup>c</sup> |  |

Data are presented as mean ± standard error. An asterisk and the connecting letter report denotes the significant statistical difference between the means in each variant category determined using ANOVA, followed by mean comparisons using student's t-test at p<0.05.
<sup>a</sup>Variation explained by a given variant (first column) in OY1 expression. All the linear regression models were significant at p<0.001.

**Table S23.** The pairwise trait correlations between OY1 transcript abundance in the emerging shoots of maize inbred lines and the CCM traits of corresponding F_1_ hybrids with *Oy1*-*N1989*/*oy1*:B73 for the 198 inbred lines common between the current study and Kremling *et al*. 2018.

|  | OY1 (count) | WT_CCM I | MT_CCM I | Ratio_CCM I | WT_CCM II | MT_CCM II | Ratio_CCM II |
| --- | --- | --- | --- | --- | --- | --- | --- |
| OY1 (count) |  | 0.110 | 0.219 | 0.196 | -0.075 | 0.169 | 0.191 |
| WT_CCM I | 0.133 |  | 0.231 | 0.004 | 0.295 | 0.212 | 0.162 |
| MT_CCM I | 0.003 | 0.001 |  | 0.967 | 0.154 | 0.865 | 0.853 |
| Ratio_CCM I | 0.007 | 0.958 | <.0001 |  | 0.093 | 0.837 | 0.837 |
| WT_CCM II | 0.303 | <.0001 | 0.030 | 0.194 |  | 0.209 | -0.007 |
| MT_CCM II | 0.020 | 0.003 | <.0001 | <.0001 | 0.003 |  | 0.973 |
| Ratio_CCM II | 0.009 | 0.023 | <.0001 | <.0001 | 0.921 | <.0001 |  |

The upper half contains Pearson's correlation coefficient and the lower half contains correlation p-values for each pairwise trait comparison. Self-comparisons (diagonal) are left blank.

## Supplemental figure

**Figure S1.**
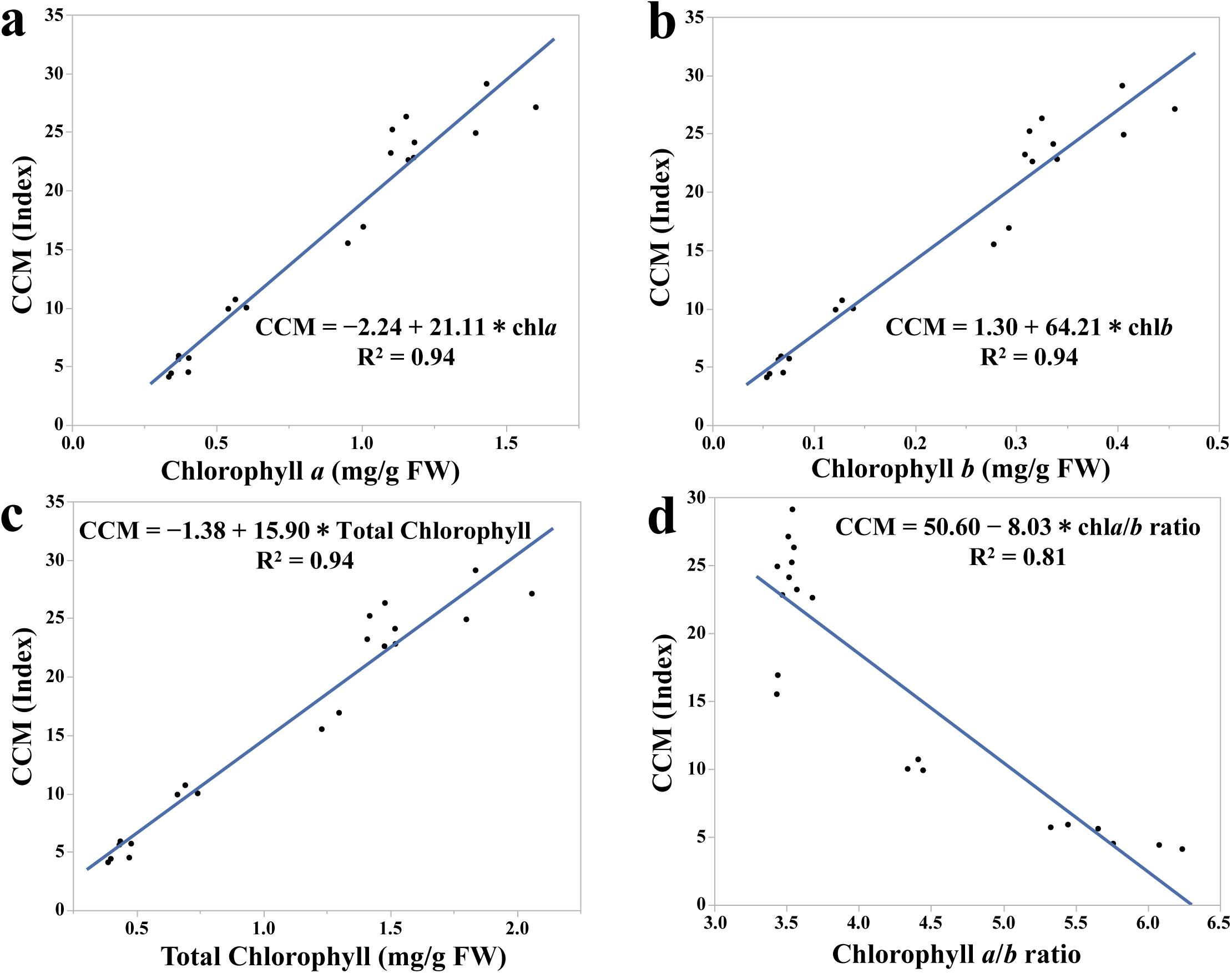
The linear regression of the chlorophyll pigment measurements using non-destructive CCM-200 plus meter (expressed as CCM index) and absolute chlorophyll pigment quantification using the spectrophotometric method from the same leaf.

**Figure S2.**
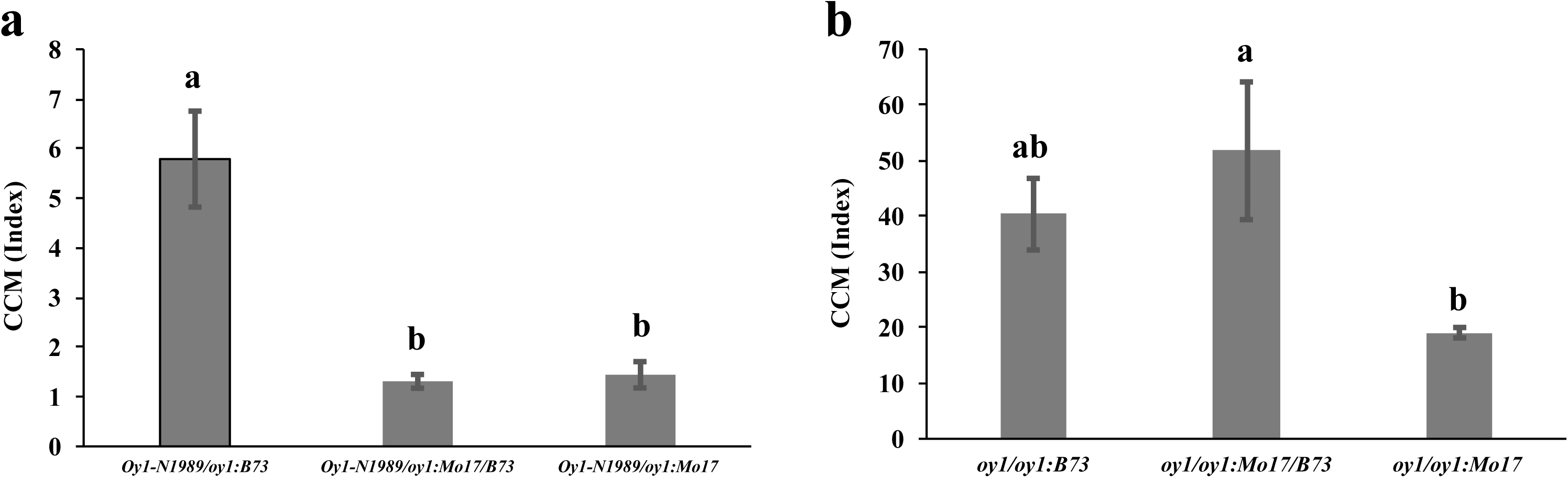
The CCM quantification of the (a) mutant (*Oy1-N1989/oy1*), and (b) wild-type (*oy1/oy1*) siblings in B73, Mo17 x B73, and Mo17 (BC_6_ generation) genetic background at 30 days after planting.

**Figure S3.**
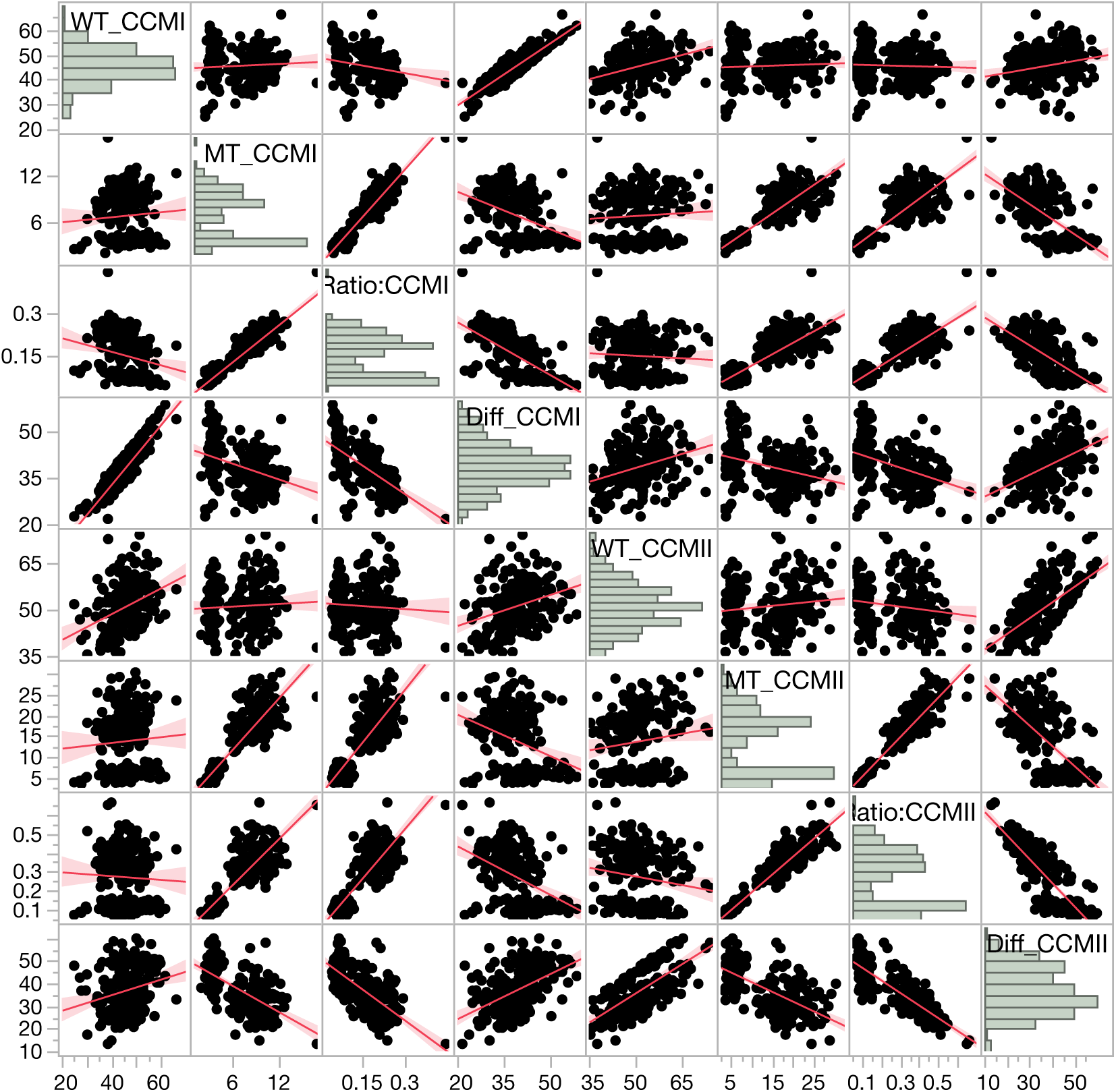
The pairwise scatter plot of primary trait measurements in IBM x *Oy1-N1989/oy1*:B73 F_1_ populations.

**Figure S4.**
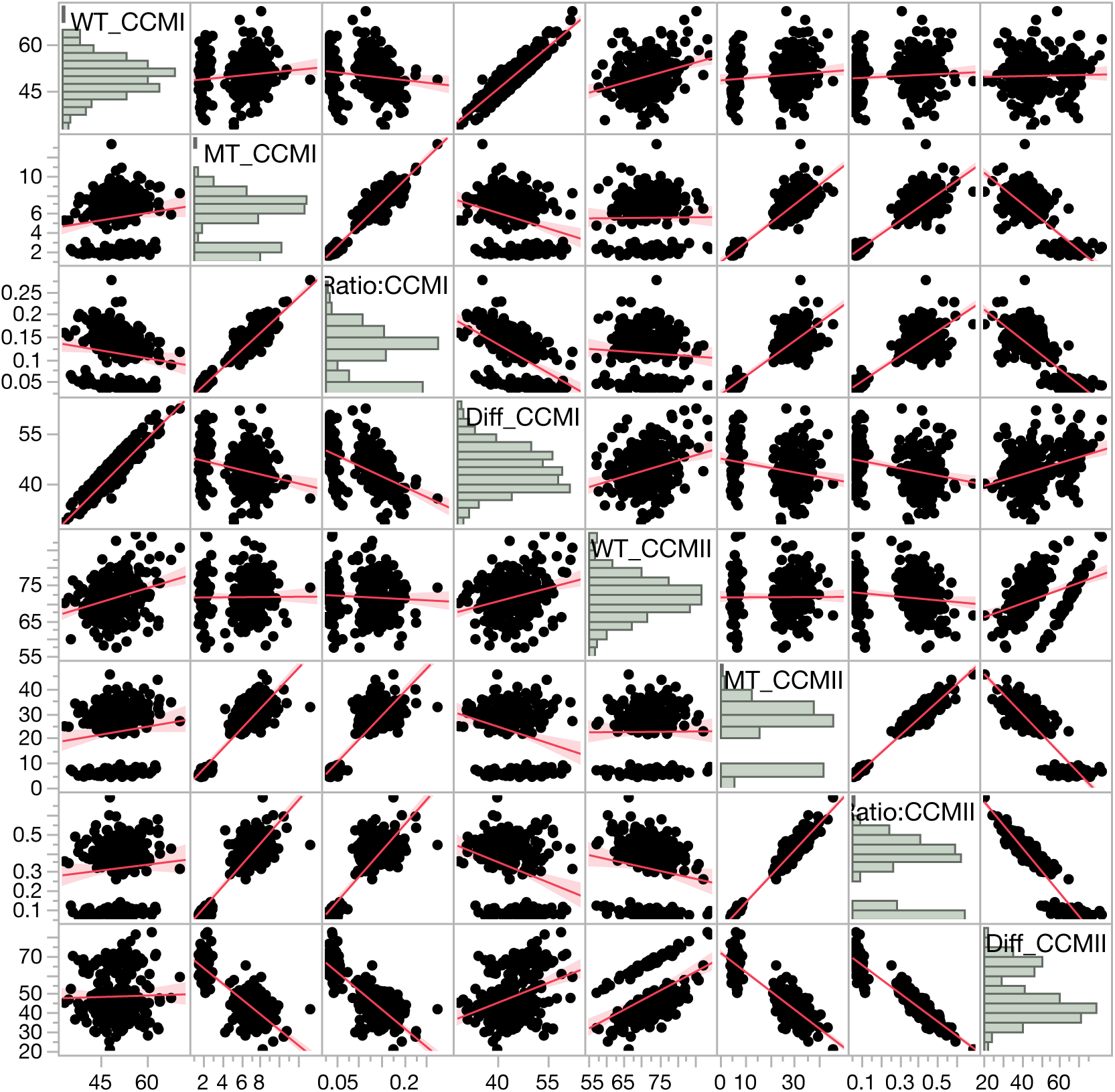
The pairwise scatter plot of primary trait measurements in Syn10 x *Oy1-N1989/oy1*:B73 F_1_ populations.

**Figure S5.**
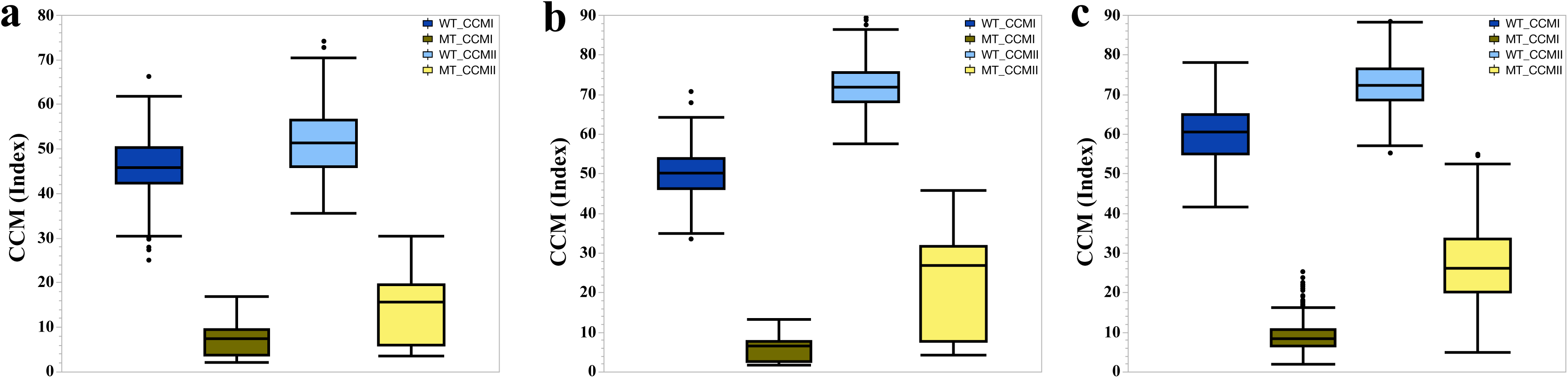
The CCMI and CCMII distribution in the wild-type (WT) and mutant (MT) siblings of (a) IBM x *Oy1-N1989/oy1*:B73 F_1_ populations, (b) Syn10 x *Oy1-N1989/oy1*:B73 F_1_ populations, and (c) MDL x *Oy1-N1989/oy1*:B73 F_1_ populations.

**Figure S6.**
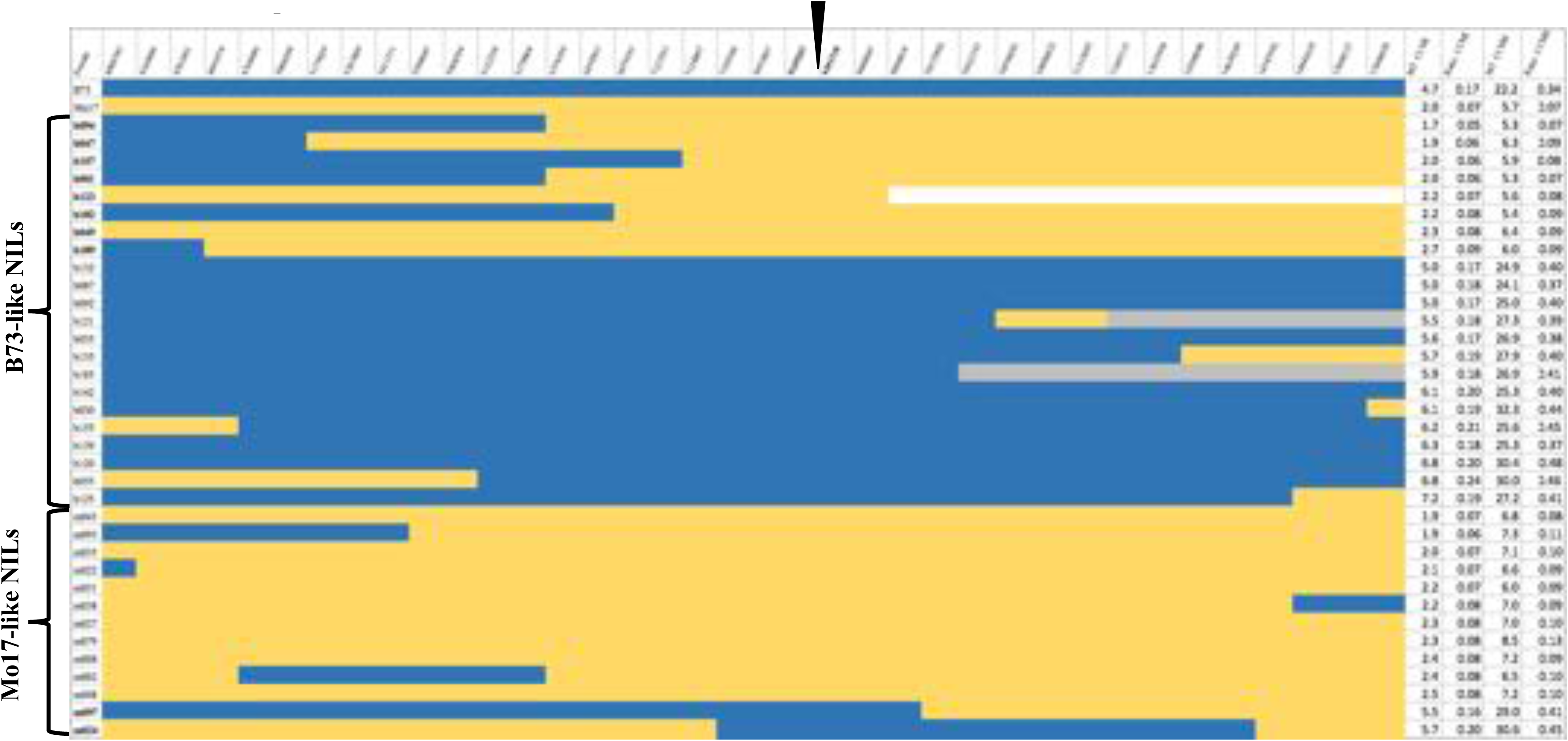
The cartoon showing *vey1* validation in BM-NILs x *Oy1-N1989/oy1*:B73 F_1_ populations. The first column shows the female parent of each cross, Colored figure shows the genotypes (B73, Mo17, Heterozygous, and missing colored as blue, golden, grey and white respectively) at a given SNP position (X-axis of the left figure; physical position from B73 RefGen v2); and position of *oy1* locus (between the two SNPs that are highlighted by a black arrow). The average (five replications) of CCM trait values in mutant siblings and their ratios (mutant/wild-type) are shown on the extreme right (last four columns) of the figure. The parental (B73 and Mo17) crosses with *Oy1-N1989/oy1*:B73 that were planted as checks in this experiment are shown in the first two rows for comparison.

**Figure S7.**
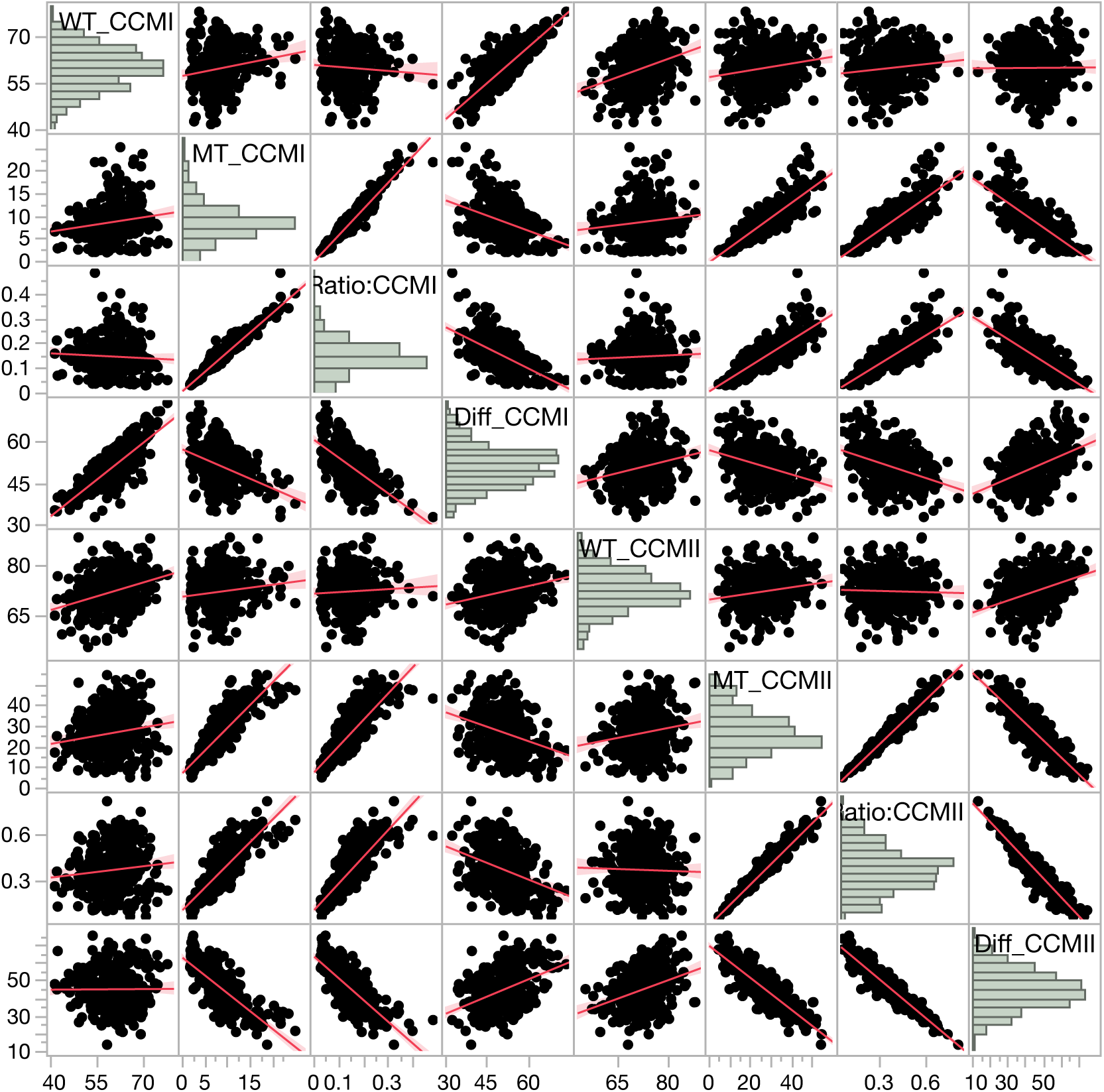
The pairwise scatter plot of primary trait measurements in MDL x *Oy1-N1989/oy1*:B73 F_1_ populations.

**Figure S8.**
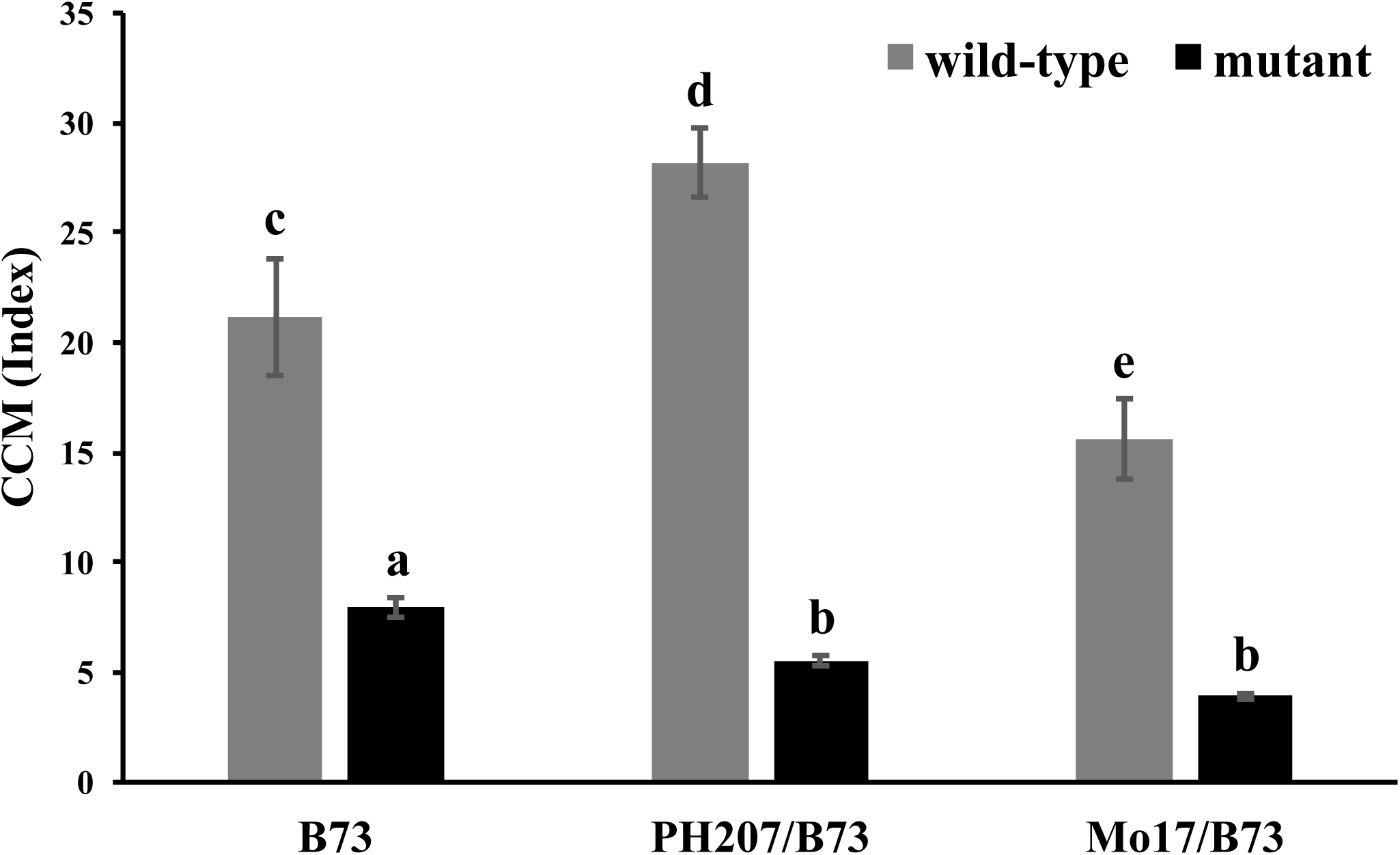
The chlorophyll approximation (using CCM) from the middle of the third leaf in the greenhouse grown F_1_ maize seedlings from a cross of B73, Mo17, and PH207 inbred lines (ear-parents) with *Oy1-N1989/oy1*:B73 plants (pollen-parent) at the V3 developmental stage. The CCM values are presented as mean with standard deviation (error bars). The connecting letter report indicates the statistical significance calculated using ANOVA with post-hoc analysis using Tukey’s HSD with p<0.05 among all genotypes. The sample size (n) for each genotype group varied from five to eight plants. Check supplemental table for details.

**Figure S9.**
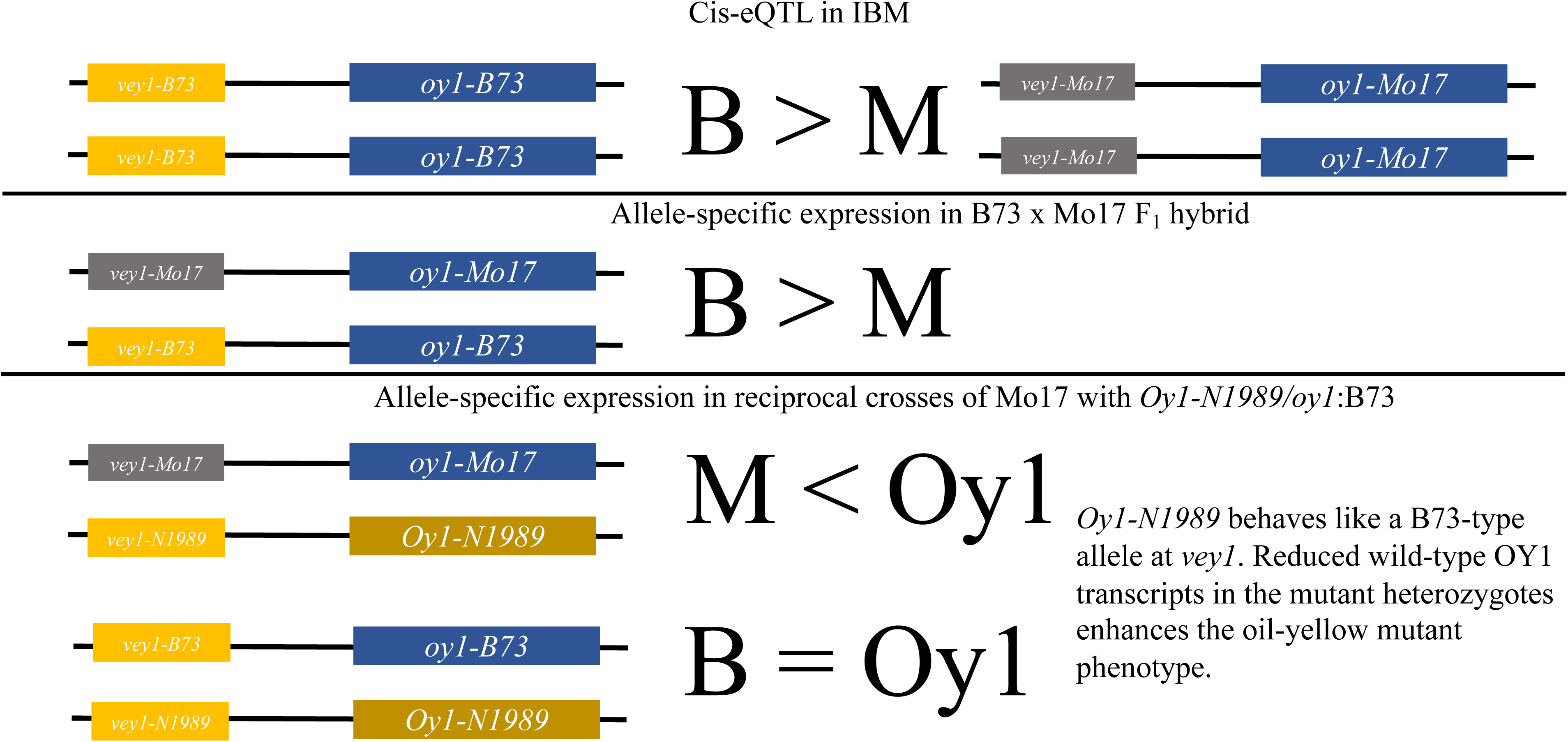
The proposed model for cis-acting regulatory variation as the basis of *vey1.*

## Supplemental files

1. S1_IBM_F1_Rqtl_input: CSV file with average CCM values and genotypic data of IBM x *Oy1-N1989/oy1*:B73 F_1_ population formatted for R/qtl.

2. S2_Syn10_F1_Rqtl_input: CSV file with BLUP value of CCM traits and genotypic data of Syn10 x *Oy1-N1989/oy1*:B73 F_1_ population formatted for R/qtl.

